# Directed evolution of a bacterial leucyl tRNA in mammalian cells for enhanced noncanonical amino acid mutagenesis

**DOI:** 10.1101/2024.02.19.581038

**Authors:** Rachel L. Huang, Delilah Jewel, Rachel E. Kelemen, Quan Pham, Shu Wang, Soumya Jyoti Singha Roy, Zeyi Huang, Samantha D. Levinson, Bharathi Sundaresh, Suyen Espinoza Miranda, Tim Van Opijnen, Abhishek Chatterjee

## Abstract

The *E. coli* leucyl-tRNA synthetase (EcLeuRS)/tRNA^EcLeu^ pair has been engineered to genetically encode a structurally diverse group of enabling noncanonical amino acids (ncAAs) in eukaryotes, including those with bioconjugation handles, environment-sensitive fluorophores, photocaged amino acids, and native post-translational modifications. However, the scope of this toolbox in mammalian cells is limited by the poor activity of tRNA^EcLeu^. Here, we overcome this limitation by evolving tRNA^EcLeu^ directly in mammalian cells using a virus-assisted selection scheme. This directed evolution platform was optimized for higher throughput such that the entire acceptor stem of tRNA^EcLeu^ could be simultaneously engineered, which resulted in the identification of several variants with remarkably improved efficiency for incorporating a wide range of ncAAs. The advantage of the evolved leucyl tRNAs was demonstrated by expressing ncAA mutants in mammalian cells that were challenging to express before using the wild-type tRNA^EcLeu^, by creating viral vectors that facilitated ncAA mutagenesis at a significantly lower dose, and by creating more efficient mammalian cell lines stably expressing the ncAA-incorporation machinery.

## Introduction

Site-specific incorporation of noncanonical amino acids (ncAAs) into proteins in mammalian cells, using an engineered nonsense-suppressing aminoacyl-tRNA synthetase (aaRS)/tRNA pairs, is a powerful technology with innumerable potential applications.^*1-4*^ To avoid cross-reaction with host counterparts, these engineered pairs are typically imported into mammalian cells from a distant domain of life, such as bacteria or archaea.^*1-3*^ However, an unwanted consequence of this strategy is the poor activity of the heterologous tRNAs in the new host due to their suboptimal interaction with foreign components, such as tRNA biogenesis and processing machinery, elongation factors, and the ribosome.^*5-7*^ To overcome such poor intrinsic efficiency, it is necessary to overexpress them by delivering a large number of tRNA genes into the host cell.^*5-8*^ Although this can be achieved through transient transfection of plasmids encoding multiple copies of the tRNA gene, this approach becomes more difficult to implement in cells that are not easily transfected. Moreover, this requirement poses significant challenges for the creation of mammalian cells that stably integrate the machinery for ncAA incorporation, which is necessary for both commercial and advanced research applications.^*8, 9*^ Previous studies have revealed that many hundreds of tRNA genes must be integrated into the host genome to achieve efficient ncAA incorporation in stable cell lines, which significantly complicates their development as well as maintenance of stability during propagation.^*7, 8*^

The *E. coli* leucyl-tRNA synthetase (EcLeuRS)/tRNA pair is one of the most versatile platforms for site-specific ncAA mutagenesis in eukaryotes.^*1-3, 10-19*^ EcLeuRS has been engineered to genetically encode an impressive variety of structurally diverse ncAAs with both aliphatic and aromatic side chains (**Figure S1**).^*1-3, 10-19*^ Many useful ncAAs have been genetically encoded using this pair, including those with bioorthogonal conjugation handles,^*14, 15*^ environment-sensitive fluorophores,^*12, 16*^ photocaged versions of serine, cysteine,^*10*^ and selenocysteine,^*17*^ as well as native post-translational modifications such as citrulline.^*18*^ This pair can also be combined with pyrrolysyl and *E. coli* tryptophanyl pairs to enable site-specific incorporation of multiple distinct ncAAs into proteins expressed in mammalian cells.^*15, 20, 21*^ The ability to develop engineered variants of tRNA^EcLeu^ with improved activity in mammalian cells will expand the scope of such established applications, as well as develop new ones. The factors that limit the performance of such heterologous tRNAs in mammalian cells is poorly understood, and their discovery is complicated by the complex, multifaceted biology of tRNAs involving, expression, maturation, post-transcriptional modification, aminoacylation, and subsequent interaction with components of the translation system (e.g., elongation factors and the ribosome).

Directed evolution has been used with much success to improve the activities of suppressor tRNAs in *E. coli* for ncAA mutagenesis,^*22-32*^ but technical challenges have precluded the implementation of analogous approaches in higher eukaryotes. Recently, we developed a virus-assisted directed evolution strategy for engineering tRNAs (VADER) in mammalian cells (**Figure 1**), that linked the activity of the *M. mazei* pyrrolysyl (tRNA^Pyl^) to the proliferation of adeno-associated virus (AAV).^*5, 33*^ Using VADER, we could identify mutants of tRNA^Pyl^ with significantly higher activities from naïve synthetic libraries. Here, we extend the use of VADER for the directed evolution of tRNA^EcLeu^ and developed mutants with remarkably enhanced activities in mammalian cells. An optimized VADER workflow with higher capacity was combined with a novel library generation scheme, that uses insights from the consensus sequences of homologous bacteria-derived leucyl tRNAs, as well as a pairwise randomization scheme to generate smarter acceptor stem library of tRNA^EcLeu^ with a higher density of functional mutants. The engineered leucyl tRNAs enabled improved genetic incorporation of many useful ncAAs in mammalian cells, and facilitated expression of ncAA mutants that were challenging to achieve before. Its advantage was further demonstrated by developing a viral delivery vector that is effective at a much lower multiplicity of infection (MOI; virus-to-cell ratio), as well as significantly more efficient generation a stable mammalian cell line for ncAA mutagenesis using the EcLeu pair. These leucyl tRNAs for improved genetic incorporation of ncAAs (LeuIGIs) will be beneficial for improving the robustness and the scope of this toolbox in mammalian cells.

**Figure 1.**
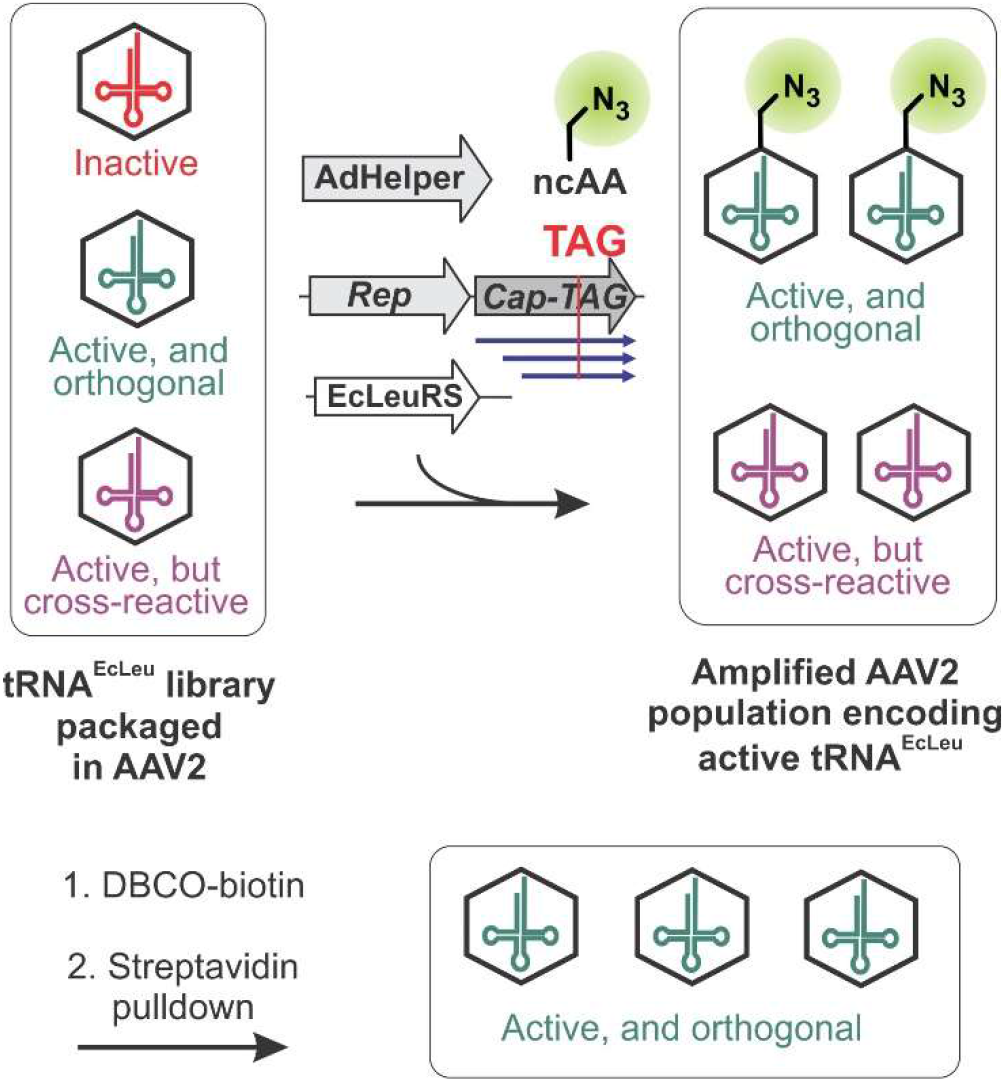
The VADER Selection Scheme. AAV2 encoding the synthetic nonsense-suppressing tRNA library are used to infected HEK293T cells at low multiplicity of infection (MOI). Additional genetic elements (AAV Rep-Cap genes, helper genes from adenovirus, a cognate aaRS to charge the tRNA) needed to support AAV replication are supplied *in trans* by transient transfection. The Cap gene encodes a nonsense codon such that only active tRNA members that suppress it can generate progeny virus. Orthogonal tRNAs that interact with its cognate aaRS facilitate the incorporation of an azido-ncAA into a surface-exposed site of AAV capsid, allowing their selective enrichment, following bioorthogonal attachment of biotin.

## Results and discussion

### Adapting VADER to enable the directed evolution of tRNA^EcLeu^

The VADER scheme subjects an AAV-encoded tRNA mutant library to two selective steps: selective amplification, and bioorthogonal capture (**Figure 1**).^*5*^ For conditional amplification, the gene encoding AAV capsid proteins (Cap) is inactivated with a TAG stop codon and is supplied *in trans* to support virus replication.^*5, 34-36*^ Consequently, only those incoming AAV particles encoding an active TAG-suppressing mutant tRNA are able to facilitate the expression of full-length capsid proteins and produce progeny virus. The tRNA library may contain undesirable mutants that suppress the TAG codon, but cross-react with a host aaRS. To remove such cross-reactive mutants, which would survive the selective amplification step, VADER uses an azide-containing ncAA as a substrate for the tRNA being engineered, and introduces it at a permissive surface-exposed site on the AAV capsid. Only active and non-cross-reactive tRNA mutants will facilitate the incorporation of the azide-ncAA onto the progeny virus particles, enabling their selectively isolation through bioorthogonal biotin attachment followed by avidin pulldown.^*5*^ To use VADER for the directed evolution of tRNA^EcLeu^, we needed to ensure that: A) the EcLeuRS/tRNA^EcLeu^ pair can suppress a TAG codon at the surface exposed site of AAV-Cap to allow efficient virus production, and B) this pair can incorporate a ncAA containing an azide (or another suitable bioorthogonal conjugation handle) into the AAV-Cap to facilitate the bioorthogonal capture step.

We have previously reported a polyspecific EcLeuRS mutant (PLRS1) that efficiently charges several ncAAs with linear aliphatic side-chains containing different conjugation handles, including an azide.^*15*^ We attempted to use the PLRS1/tRNA^EcLeu^ pair to decode a TAG codon at the surface-exposed T454 site of AAV-Cap and produce ncAA-decorated functional AAV (**Figure S2**). Surprisingly, no functional virus was produced when using the azide-containing ncAA LCA, one of the most efficient substrates of PLRS1. However, robust virus production was observed using AzK, a structurally similar ncAA that is a considerably less efficient substrate of PLRS1.^*15*^ This is likely due to the delicate nature of the multifunctional virus capsid proteins and their complex assembly process, which often respond in unpredictable ways upon alterations. Nonetheless, successful packaging of AAV-Cap-454-AzK, facilitated by the TAG-suppressing PLRS1/tRNA^EcLeu^ pair, makes it possible to apply VADER to engineer tRNA^EcLeu^.

We then used a mock-selection experiment to confirm that active tRNA^EcLeu^ mutants can be enriched using VADER (**Figure S3**). A defined mixture of two AAV vectors, one encoding tRNA^EcLeu^ and a wild-type mCherry reporter, and another encoding tRNA-pyrrolysyl (Pyl) and a wild-type EGFP reporter, in a 1:1,000 ratio, was subjected to VADER. We have previously shown that tRNA^Pyl^ is orthogonal to EcLeuRS and would be inactive under the conditions of this experiment. The composition of the resulting progeny virus mixture was evaluated by FACS analysis to reveal ∼28,000-fold enrichment of the EcLeu encoding AAV, further confirming that VADER can be used for engineering tRNA^EcLeu^.

### Virus-assisted directed evolution of tRNA^EcLeu^

To identify variants of tRNA^EcLeu^ that exhibit higher efficiency in mammalian cells, we focused on engineering its acceptor stem, which directly interacts with various components of the translation system. This strategy has been used with much success for tRNA evolution in both *E. coli* and in mammalian cells.^*5, 22-25, 28*^ Using our first-generation VADER protocol, it was possible to process up to a few thousand mutants per selection. Consequently, we were only able to simultaneously randomize up to 3 base pairs in the stem regions of tRNA^Pyl^. Even though we were still able to identify beneficial mutations that resulted in modest improvements in tRNA activity, the inability to simultaneous randomize larger segments would preclude the identification of synergistic distal mutations providing further enhancement. Using systematic optimization of the VADER workflow, we have now significantly improved its capacity, allowing the selection of libraries containing up to 10^5^ mutants. However, this improved capacity was still insufficient to allow simultaneous randomization/selection of the entire acceptor stem, complete randomization of which would create approximately 2.7×10^8^ distinct mutants. We used two key strategies to rationally reduce the size of the acceptor-stem library, without significantly compromising its functional diversity: A) Traditional tRNA stem libraries are generated by independently randomizing each of the base-paired strands. Consequently, such libraries contain a large number of mutants that fail to base pair, and are typically inactive. Using oligo-pool synthesis technology, we employed a novel pairwise-randomization scheme, where base pairing between distal residues was maintained in all library members. Using this approach, the library size can be drastically reduced without compromising functional diversity: while traditional full-randomization of the acceptor stem would generate a 2.7×10^8^ possible mutants, the pairwise randomization creates only 2.8×10^5^ (including the G:U/U:G ‘wobble’ pairs). B) We also used sequence alignment of 682 bacterial leucyl tRNAs to identify strongly conserved sequence elements (**Figure 2a**) that may be functionally important (e.g., those important for aaRS recognition), and restricted alterations to these conserved regions (**Figure 2b**). Together, these two strategies allowed us to restrict the size of the tRNA^EcLeu^ acceptor stem library to 7,776 mutants.

**Figure 2.**
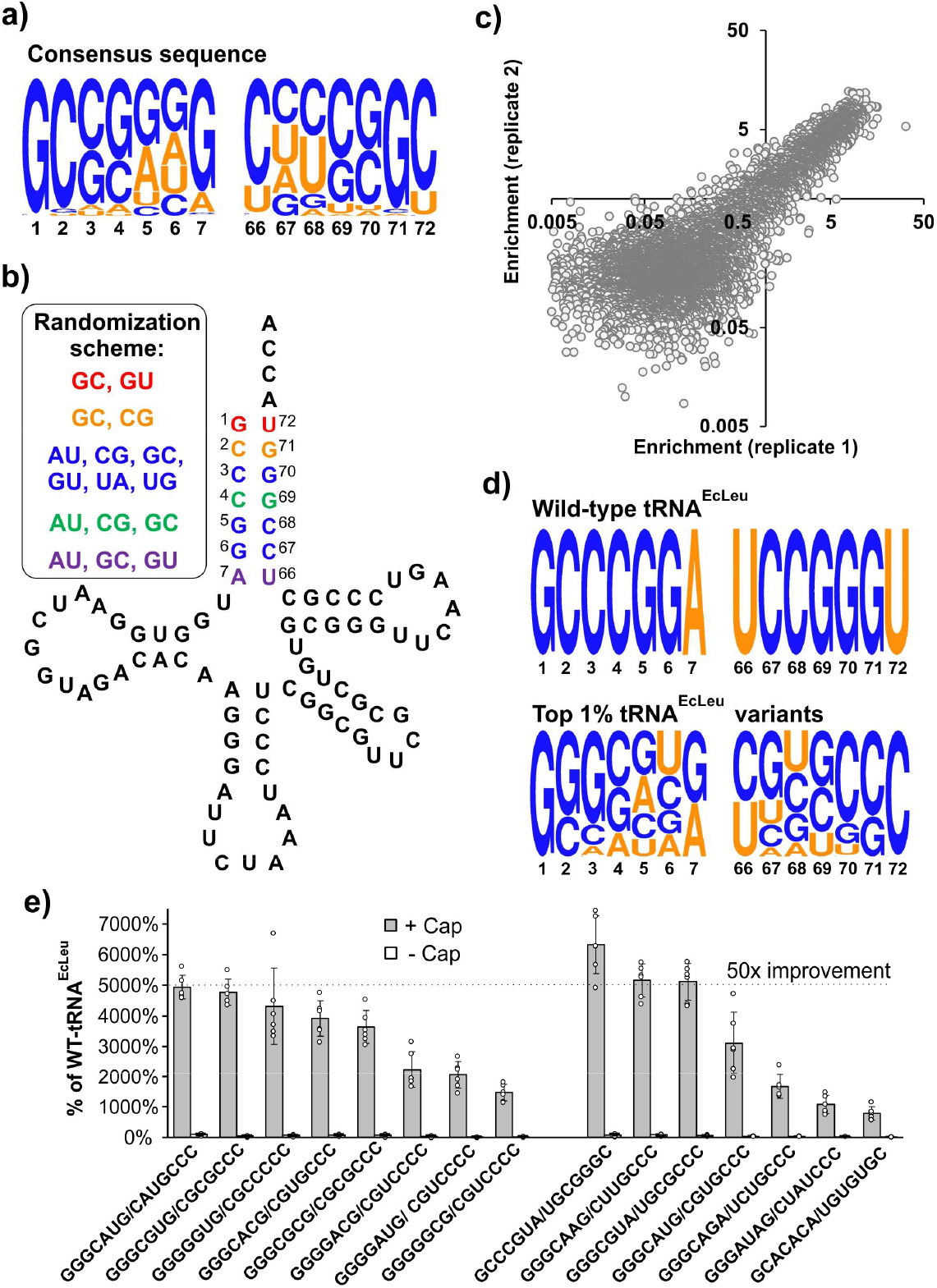
tRNA^Leu^ Library Design and Next Generation Sequencing Selection Data. **a)** WebLogo representation of the sequence alignment of the acceptor stem from 682 bacterial tRNAs, demonstrating relative abundance of each residue at each position. The numbering scheme is relative to tRNA^EcLeu^ (panel b). **b)** Cloverleaf structure of wild-type *E. coli* tRNA_CUA_^Leu^ showing the library generation scheme by randomizing the acceptor stem base pairs. Base pairing was maintained in all library members by introducing pairwise substitutions at each position according to the colored scheme shown in the box. Randomization was restricted to maintain conserved sequence elements (panel a). **c)** Enrichment of each mutant in the library upon VADER selection was measured using their normalized abundance in the library after and before the selection (abundance after/abundance before). Two biological replicates of the selection were performed, and the enrichment of each mutant observed in these two replicates were plotted against each other. **d)** Consensus sequence of top 1% (most enriched) sequences is shown. The sequence of the WT-tRNA^EcLeu^ is provided as reference. **e)** The activity of each tRNA^EcLeu^ mutant was tested by co-transfection into HEK293T cells with PLRS1 and EGFP-39TAG in the presence or absence of 1 mM Cap (also see **Figure S5**). Expression of EGFP-39TAG was measured in cell-free extract, normalized relative to mCherry expression (present in the tRNA-expressing plasmid), and shown relative to the activity of wild-type tRNA^EcLeu^.

This library was cloned into our virus packaging plasmid with a 185-fold sequence coverage, and was packaged into wild-type AAV2 particles. AAV-encoded tRNA^EcLeu^ library was subjected to the VADER selection scheme using PLRS1 and AzK as the cognate aaRS and ncAA, respectively. The composition of the library was characterized by Illumina MiSeq both before and after the selection (in duplicate), which identified the presence of nearly all of the theoretical mutants (**Figure S4**). The degree of enrichment for each mutant was calculated from their relative abundance in the library before and after the selection. The observed enrichment factors for the library members were largely consistent between the two replicates (**Figure 2c**). A large number of mutants showed high levels of enrichment relative to wild-type tRNA^EcLeu^ (**Figure S4b**), indicating significant plasticity of its acceptor stem. Analysis of the top 1% most enriched mutants revealed significant deviation from the wild-type sequence (**Figure 2d**). We constructed 15 mutants based on their high degree of enrichment, as well as homology with the consensus sequence of the top 1% most enriched mutants (**Figure 2e, Supplementary Table S2**). These mutants were tested by co-transfecting them into HEK293T cells with plasmids encoding PLRS1 and EGFP-39-TAG in the presence or absence of the ncAA **Cap**. The tRNA-encoding plasmid also contained a wild-type mCherry reporter to provide an internal control (**Figure S5**). All of the tested mutants showed remarkable enhancement in activity relative to wild type tRNA^EcLeu^, with the most improved ones showing >60-fold improvement (**Figure 2e**). In these engineered leucyl-tRNAs for improved genetic incorporation (LeuIGIs) of ncAAs, the first base-pair in the acceptor stem changed from G:U to G:C, which is frequently observed in wild-type bacterial leucyl tRNAs (**Figure 2a**). Interestingly, some of the most active mutants also harbored a U at the 6 position in the acceptor stem, base-paired with either an A or a G.

### Further characterization of tRNA^EcLeu^ mutants with additional ncAAs

Next, we investigated how the evolved LeuIGIs improve the incorporation efficiency of a broader series of ncAAs genetically encoded using this platform (**Figure 3**). One of the most active tRNA^EcLeu^ mutants (LeuIGI1; **Figure 3a**) identified above was co-transfected into HEK293T cells along with different engineered EcLeuRS mutants and an EGFP-39TAG reporter in the presence of a variety of different known substrates of this polyspecific synthetase. Identical control experiments using WT-tRNA^EcLeu^ was also set up in parallel. The use of the evolved tRNA^EcLeu^ was associated with significant improvement of incorporation of all ncAAs tested (**Figure 3b**). A second tRNA^EcLeu^ mutant (LeuIGI2; **Figure 3a**) with similar activity, but a distinct acceptor stem sequence, was also tested in the same manner to yield similar results (**Figure S6**). Interestingly, although LeuIGIs enabled enhanced incorporation in all cases, the degree of improvement was variable across different ncAAs. In particular, the improvements observed for linear aliphatic ncAA substrates were significantly higher than those observed for aromatic ncAA counterparts (**Figure 3b**). Although the reason behind this observation is unclear, it may stem from elongation factor (EF) binding to the aminoacylated-tRNA; poor ncAA substrates for EF may enjoy a higher benefit from an engineered tRNA variant that compensates for their suboptimal affinity by enhanced EF-binding. Indeed, canonical amino acids such as aspartate that bind EF poorly, are naturally assigned to tRNAs that associate with EF-Tu with high efficiency.^*37*^

**Figure 3.**
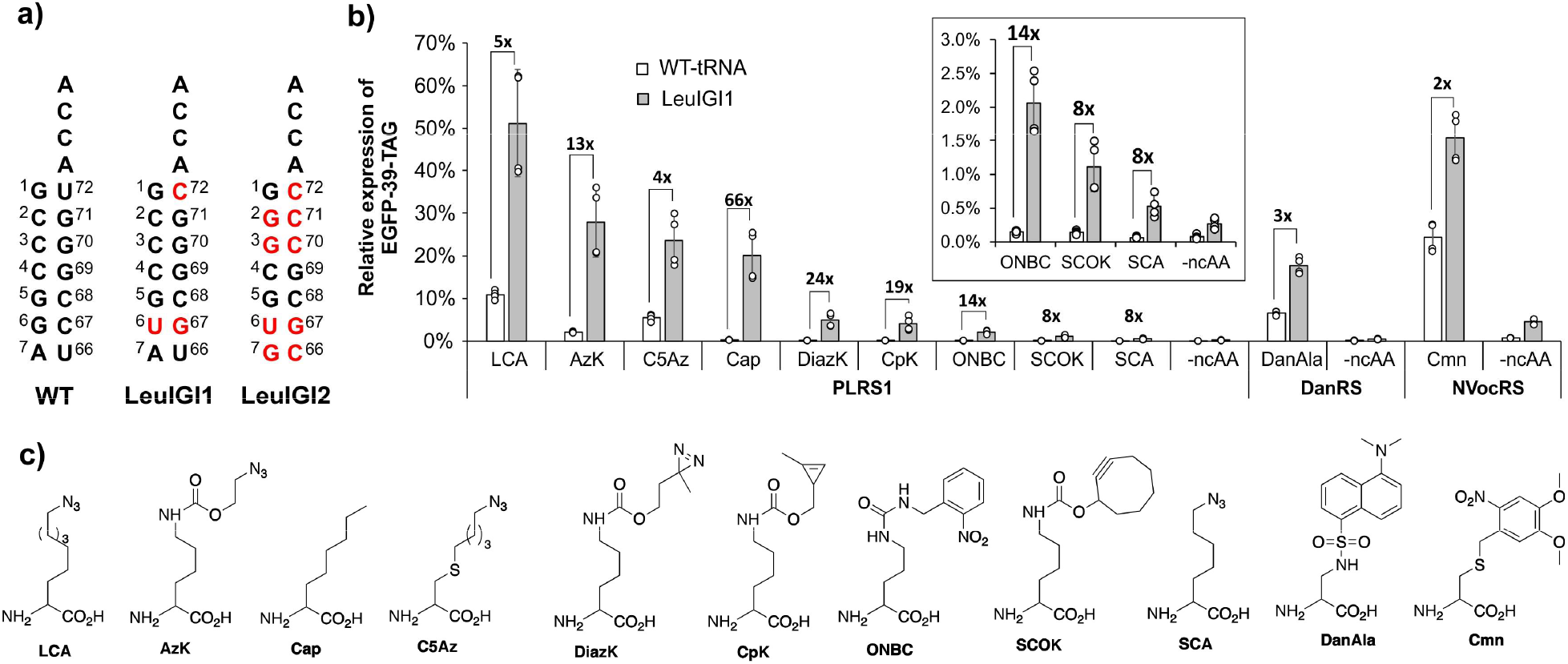
Substrate scope of engineered tRNA^EcLeu^ mutants. **a)** Sequences of the acceptor stems of wild-type tRNA^EcLeu^ LeuIGI1 and LeuIGI2. **b)** Incorporation efficiency of various ncAAs using wild-type tRNA^EcLeu^ and LeuIGI1. Measurement was performed by co-transfecting pAAV-ITR-CMV-mCherry-1xU6-LtR (either WT-tRNAEcLeu or LeuIGI1) tRNA-plasmid into HEK293T cells with plasmids encoding the appropriate pIDTSMART-CMV-EcLeuRS (PLRS1, DanRS, or NvocRS) and pAcBac1-CMV-EGFP(39TAG) in the presence or absence of 0.2 mM ncAA. Expression of EGFP-39-TAG was normalized relative to mCherry (encoded in the tRNA-expressing plasmid; **Figure S5**), and shown relative to the expression of a wild-type EGFP reporter. **c)** Structures of the ncAAs used in panel b. The characterization of LeuIGI2 activity is shown in **Figure S6**.

### LeuIGI1 activity upon controlled delivery by a viral vector

Using Northern blot analysis, we demonstrated that WT-tRNA^EcLeu^ and LeuIGI1 have similar expression levels in HEK293T cells upon transient transfection of identical plasmids (**Figure S7**). This observation shows that the improved efficiency of LeuIGI1 does not stem from a higher cellular abundance, and that it is likely intrinsically more active. We have previously found that overexpression of the tRNA by transient transfection can obscure the functional difference between mutants with varying activities due to saturation effects.^*5, 7*^ Instead, testing the activities of the tRNA variants across a variety of expression levels, which can be achieved by controlled delivery using a BacMam vector by simply changing the virus-to-cell ratio,^*5, 7, 38*^ provides a deeper insight into their comparative performance in mammalian cells. To use this assay for comparing the activities of LeuIGI1, and WT-tRNA^EcLeu^, we created two BacMam vectors (**Figure 4a**): 1) one encoding a single copy of either tRNA, and a wild-type mCherry reporter, and 2) the second encoding PLRS1 and an EGFP-39-TAG reporter. The 2^nd^ BacMam vector is used at a constant multiplicity of infection (MOI) to express the PLRS1 and the EGFP-39-TAG at a steady level, whereas the expression of the tRNA mutants is systematically increased by using higher MOI of the 1^st^ BacMam vector. We used the expression of the encoded mCherry to ensure comparable delivery of the two distinct tRNAs across all MOIs (**Figure 4b, S8**). LeuIGI1 exhibited robust TAG suppression activity even at the lowest MOI tested (5% of WT-EGFP at MOI 1), while WT-tRNA^EcLeu^ did not show detectable reporter expression until higher MOI of 5 was used (**Figure 4b, S8**). At MOI 7, LeuIGI1 showed >40% TAG suppression, while the WT-tRNA^EcLeu^ afforded <1%; a difference of ∼42-fold. These experiments highlight the superior performance of LeuIGI1, particularly under tRNA-limiting conditions. Furthermore, we have previously demonstrated that BacMam is one of the most effective viral delivery platforms for GCE applications across a broad range of mammalian cells.^7,38^ LeuIGI1 would significantly expands its scope by allowing robust TAG suppression at a lower MOI.

**Figure 4.**
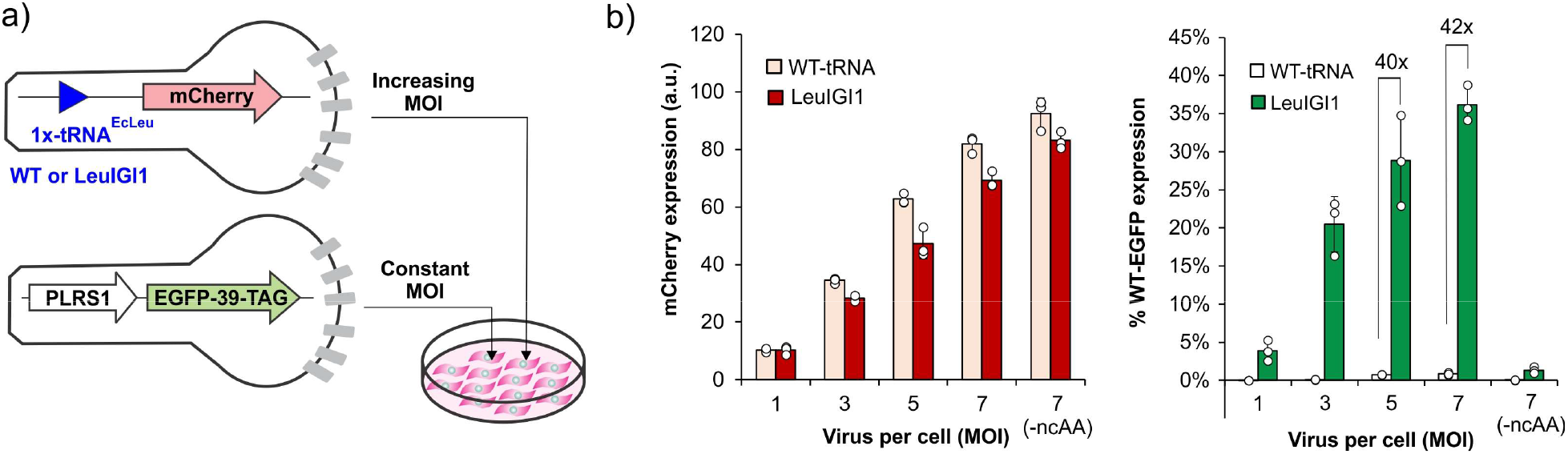
Relative activities of tRNA^EcLeu^ and LeuIGI1 upon controlled delivery using Bacmam vectors. **a)** Scheme of the experiment for using BacMam vectors, where the PLRS1/EGFP-39-TAG virus is delivered at a constant MOI of 100, whereas the tRNA-mCherry virus is used at increasing MOI. (b) Expression of EGFP-39-TAG as increasing MOI of tRNA-BacMam vector is used (green; right). Expression of mCherry (red; left) in the same experiments shows equivalent delivery of both tRNAs, which increases proportionately with higher MOI. Expression of the fluorescent reporters was measured in HEK293T cell-free extract. PLRS1 and Cap was used in these experiments. **Figure S8** shows representative images.

### LeuIGIs enable expression of previously challenging targets

To demonstrate that the improved efficiency of LeuIGIs can be beneficial beyond EGFP-39-TAG reporter expression, we focused on two specific applications. Citrullination is an important post-translational modification (PTM) of mammalian proteins, with important roles in health and disease.^*39-43*^ Due to the promiscuous activities of protein-arginine deiminases (PADs), enzymes responsible for installing this PTM, it is challenging to install this modification at specific sites of target proteins in a homogeneous manner to study its impact on structure and function. Recently, using an engineered EcLeuRS/tRNA^EcLeu^ pair, we genetically encoded a photocaged citrulline derivative ONBC (**Figure 3c**) in mammalian cells, allowing its site-specific incorporation followed by post-translational photo-decaging to generate homogeneously citrullinated protein samples.^*18*^ Although enabling, the incorporation efficiency was somewhat modest for this ncAA, and the expression levels of certain mutants (e.g., PAD4-372-TAG, an auto-citrullination site of PAD4) were particularly low. To explore if the use of LeuIGIs would improve the expression of such previously challenging citrulline mutants, we transfected HEK293T cells with plasmids encoding PAD4-372-TAG, the EcLeuRS mutant, and 4 copies of either the WT-tRNA^EcLeu^ or LeuIGI1. As a control, we also transfected the cells with the wild-type PAD4. Subsequent Western blot analysis revealed barely detectable signal for the PAD4-372-TAG mutant expressed using WT-tRNA^EcLeu^, but robust expression was observed using the LeuIGI1 variant (**Figure S9**), underscoring its utility for improving the scope of this technology.

### Facile development of a stable mammalian cell line for ncAA incorporation using LeuIGI1

The ability to develop mammalian cell lines stably expressing engineered aaRS/tRNA pairs for ncAA mutagenesis is crucial for basic research applications, including extension of this technology to animal models, as well as in biotechnology to enable expression of ncAA-proteins in scale.^*8, 9, 44, 45*^ Several strategies have been developed for generating such stable cell lines, including the genomic incorporation of the transgenes using random integration,^*8, 9*^ or transposon-mediated integration,^*44*^ as well as the use of an Eppstein-Barr virus (EBV) vector that is maintained as an episome.^*45*^ Previous reports have shown that to generate stable mammalian cell lines with robust ncAA incorporation efficiency, hundreds of tRNA genes/cell are needed.^*8*^ The need for such a large copy number significantly complicates both the generation and the maintenance of such cell lines. Given LeuIGI1 exhibits significantly higher activity at similar expression level as WT-tRNA^EcLeu^, it should enable more facile generation of stable cell line for ncAA incorporation. We sought to explore this possibility using the EBV stable episome system, as it avoids the need for genomic integration and the associated context effects that cannot be replicated across experiments.^*45*^ Two EBV vectors were created, each encoding 4 copies of either WT-tRNA^EcLeu^ or LeuIGI1, and a CMV-driven polycistronic gene cassette encoding the EGFP-39-TAG reporter, the puromycin-N-acetyltransferase (Pac), and PLRS1 (**Figure 5a**). Transfection followed by puromycin selection was used to establish stable polyclonal pools for both constructs using the HEK293T cell line (**Figure S10a**). FACS analysis showed that only ∼6% of cells in the pool generated using WT-tRNA^EcLeu^ exhibit robust EGFP-39-TAG reporter expression relative to the control (untransfected) population, compared to >33% cells in the LeuIGI1-derived pool (**Figure 5b, S10b**). The average fluorescence of the LeuIGI1 pool was also significantly brighter, demonstrating its clear advantage for constructing stable mammalian cell lines for ncAA incorporation.

**Figure 5.**
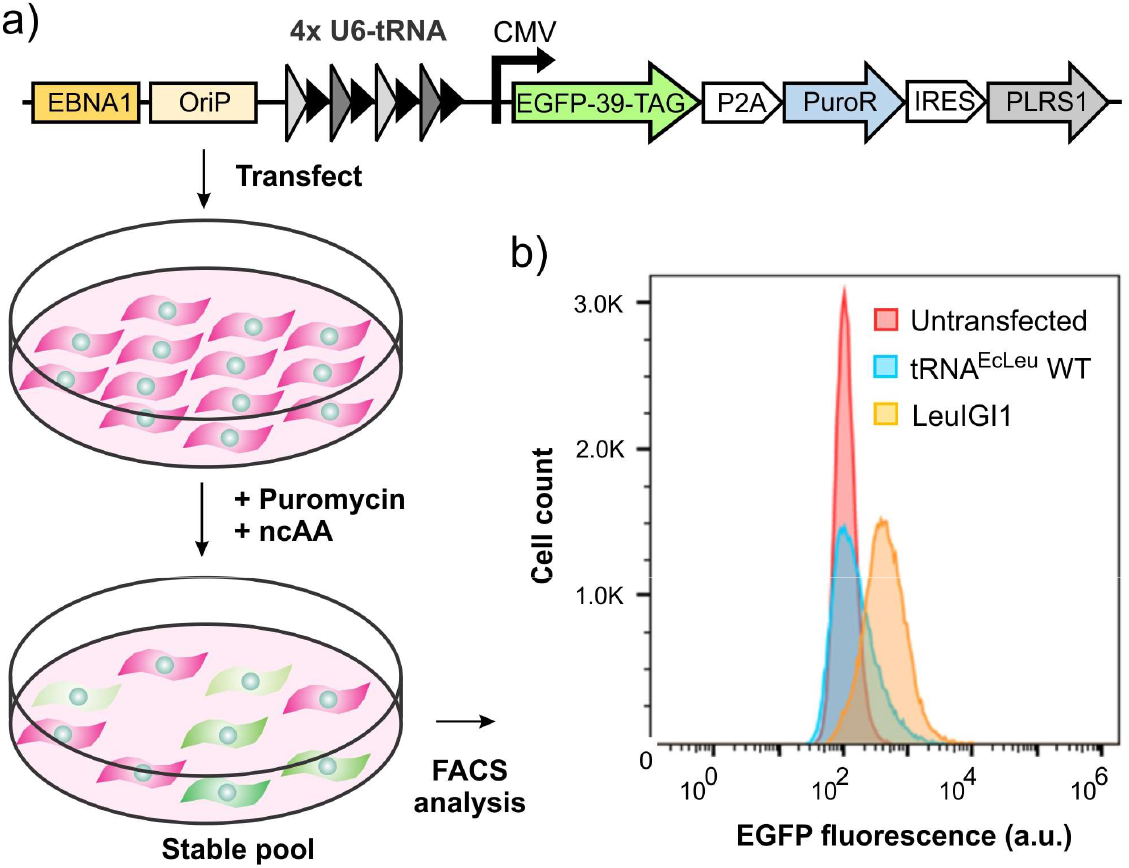
Generation of stable mammalian cell line using wild-type tRNA^EcLeu^ and LeuIGI1. **a)** The map of the EBV plasmid, and the experimental scheme, used to generate stable mammalian cell lines expressing PLRS1, tRNA^EcLeu^ (WT or LeuIGI1), an EGFP-39-TAG reporter, and the puromycin resistance gene. **b)** FACS histogram analysis (EGFP fluorescence) of the stable pool generated using puromycin selection for both WT-tRNA^EcLeu^ and LeuIGI1, and an untransfected control. Further FACS characterization is shown in Figure S10.

## Conclusions

The ncAA mutagenesis technology rely on heterologous tRNAs to minimize the chances of cross-reactivity with host aaRSs. A downside of this strategy is that such heterologous tRNAs are not optimized to interact with the numerous host machinery involved in tRNA biogenesis or translation. Although directed evolution of tRNAs has been used to overcome this limitation in *E. coli*, similar strategies were not available in higher eukaryotes. We recently developed VADER to enable systematic directed evolution of tRNAs in mammalian cells, and used it to engineer the *M. mazei* pyrrolysyl tRNA. Here we show that the VADER platform can be adapted to allow the directed evolution of tRNA^EcLeu^, which has been used to incorporate numerous useful ncAAs into proteins in mammalian cells. Simultaneous engineering of the entire acceptor stem of tRNA^EcLeu^ was made possible by: A) an improved VADER workflow capable of processing more variants, and B) a smarter library design that eliminates unpaired mutants, and uses lessons from the natural tRNA^Leu^ sequence diversity in bacteria to avoid losing important identity elements. The tRNA^EcLeu^ variants (LeuIGIs) identified here showed remarkably improved performance in several different contexts, which will be beneficial for the further application of this technology. Additionally, these advances further highlight the efficacy of VADER for tRNA engineering in mammalian cells.

## Data availability statement

The data described this study are available in the article and in the supporting information. Code used to analyze sequencing data are available from GitHub (http:github.com/chatterjeelab2022/VADER). Sequencing data is available via BioProject: https://www.ncbi.nlm.nih.gov/sra/?term=prjna1061141. Further data will be shared upon reasonable request to the corresponding author.

## Acknowledgements

We thank the National Science Foundation (MCB-1817893), and National Institutes of Health (R35GM136437 to A.C. and U01 AI124302 to T.v.O.) for financial support.

## Competing interest

A patent application has been submitted on the improved tRNA mutants on which A.C. in an inventor. A.C. is a senior advisor at BrickBio, Inc.

## Supplementary data

Experimental methods, nucleotide sequences, supplementary figures and tables. (PDF)

## Notes

https://www.ncbi.nlm.nih.gov/sra/?term=prjna1061141

