## Supporting information for "Directed evolution of a bacterial leucyl tRNA in mammalian cells for enhanced noncanonical amino acid mutagenesis"

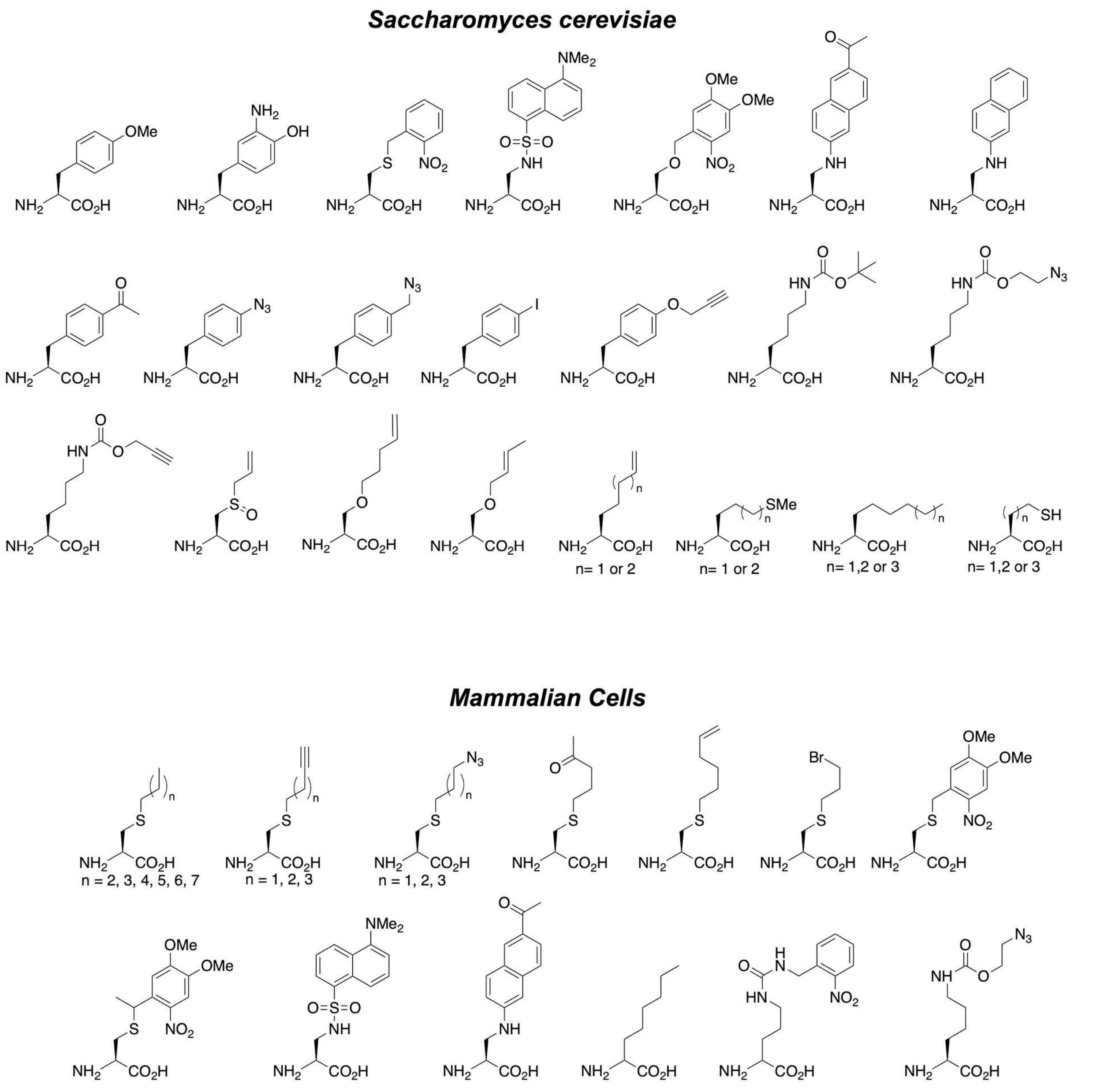


**Figure S1**. Examples of ncAAs genetically encoded in eukaryotes using the EcLeuRS/tRNA$^{EcLeu}$ pair.

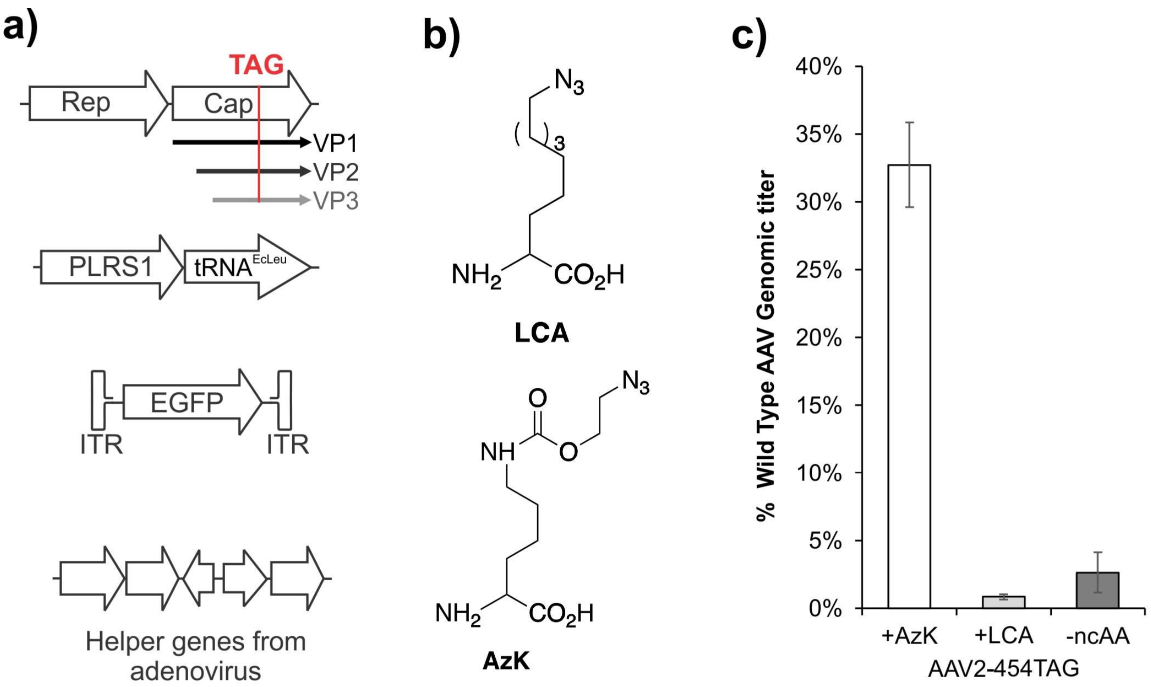


**Figure S2**. EcLeuRS/tRNA$^{EcLeu}$ pair can be used to incorporate an azide containing ncAA into the capsid of AAV. **a**) Scheme of the genetic components used for virus packaging: Rep, Cap-454-TAG (a TAG codon is introduced at the 454 position of the capsid gene), PLRS1/tRNA$^{EcLeu}$ pair, a wild-type EGFP gene encoded within the inverted terminal repeats (ITR; packaging signal), and necessary helper genes from adenovirus. **b**) Structures of LCA and AzK. **c**) AAV-454-TAG can be successfully packaged using the PLRS1/tRNA$^{EcLeu}$ pair in the presence of AzK, but not LCA. Virus production was measured by quantifying packaged genome copies by qPCR and normalized to wild-type AAV titers.

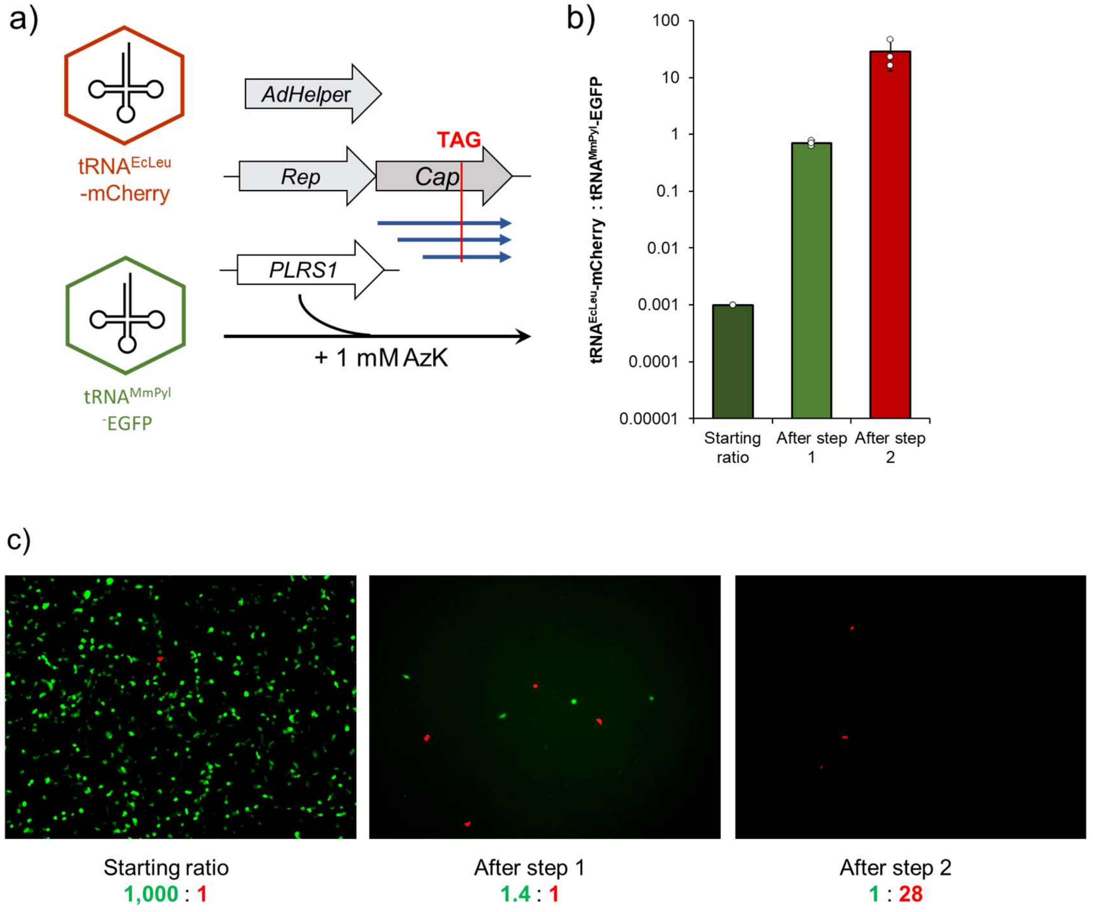


**Figure S3**. A mock selection experiment showing that VADER enables enrichment of tRNA$^{EcLeu}$ relative to tRNA$^{MmPyl}$ from a mixed population. **a)** Scheme for the mock selection experiment using two AAV vectors: one encoding tRNA$^{EcLeu}$ and an mCherry reporter (the active tRNA), and the other encoding tRNA$^{MmPyl}$ with an EGFP reporter (the inactive tRNA). Mock selection was performed in triplicates. **b)** FACS analysis of mock selection showed that >24,000-fold cumulative enrichment of tRNA$^{EcLeu}$-mCherry after a single round comprising two steps. **c)** Representative fluorescence microscopy images of cells infected with the mixed AAV population from different stages of the selection: input virus (1,000:1 ratio of inactive tRNA$^{MmPyl}$ -EGFP to active tRNA$^{EcLeu}$-mCherry), after VADER step 1 (selective amplification), and step 2 (bioorthogonal capture). The ratios below images are the average ratio of three mock selections.

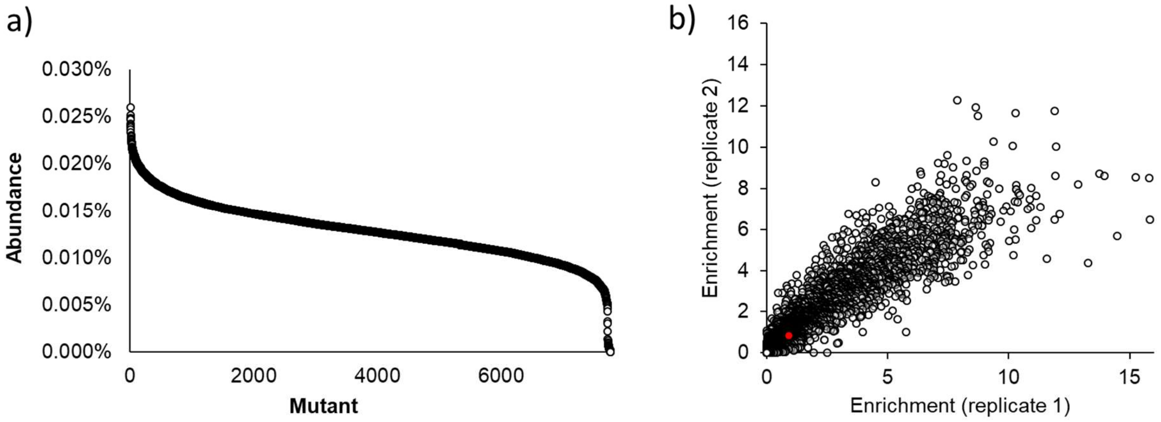


**Figure S4. a)** Distribution of unique mutants in the input library as assessed by NGS analysis. It revealed the presence of >99.9% of all designed mutants (7,773/ 7,776). **b)** Enrichment of each mutant in the $tRNA^{EcLeu}$ library upon undergoing the VADER selection scheme in two replicates are plotted against each other. The enrichment profile of the wild-type $tRNA^{EcLeu}$ is shown in red.

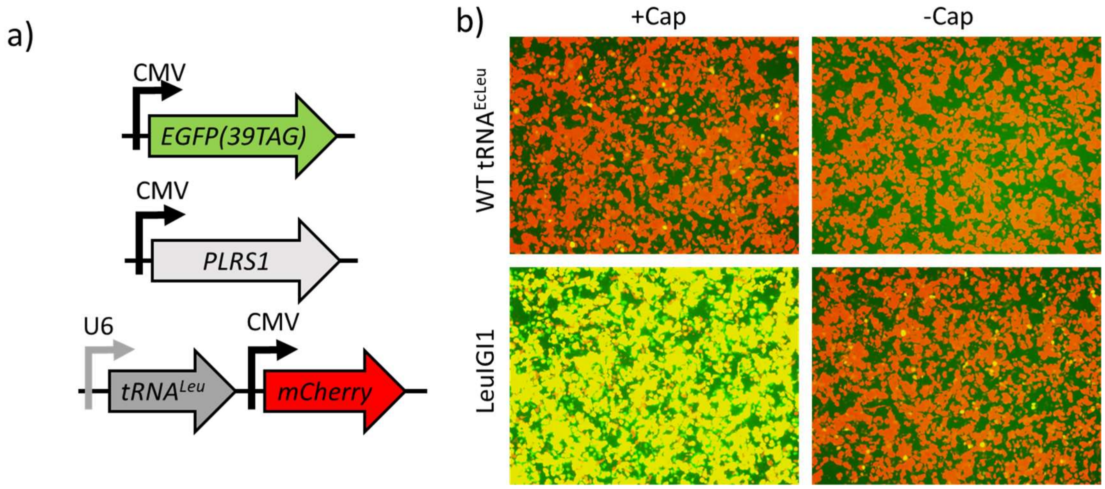


**Figure S5**. **a)** The three-plasmids used for transient transfection in HEK293T to test the activities of different tRNA variants in the presence or absence of 1 mM Cap: i) pAcBac1-CMV-EGFP-39-TAG reporter plasmid, ii) pIDTSmart-CMV-PLRS1 plasmid, and iii) pAAV plasmid harboring the tRNA and a wild-type mCherry reporter. Expression of mCherry and EGFP-39-TAG were measured by fluorescence in cell-free extract. The fluorescence of each tRNA was normalized relative to wild-type mCherry expression and then plotted as a percentage of the normalized activity of wild-type $tRNA^{EcLeu}$. **b)** Representative fluorescence microscopy images of cells transfected with wild-type $tRNA^{EcLeu}$ and LeuIGI1 in the presence or absence of 1 mM Cap according to the previously described transfection scheme in (a).

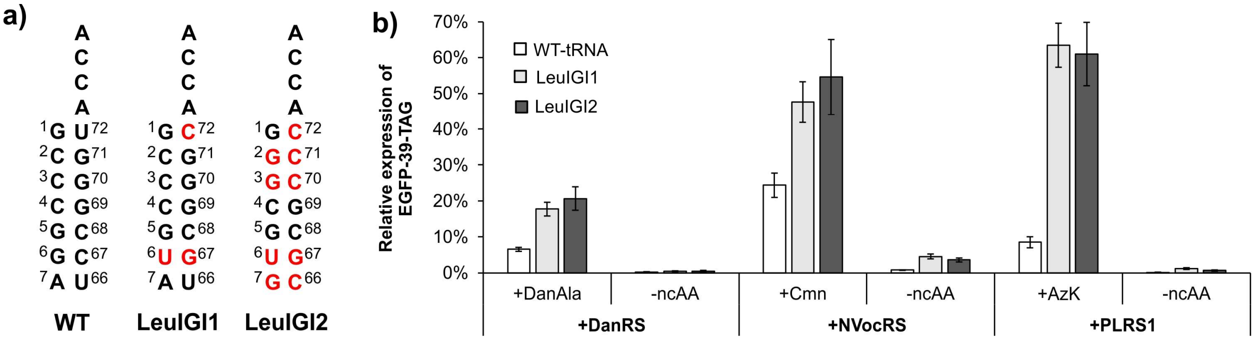


**Figure S6. a)** Acceptor stem sequences of WT tRNA$^{EcLeu}$ and LeuIGI1, and LeuIGI2 with mutations shown in red. **b)** Incorporation efficiency of various ncAAs using WT tRNA$^{EcLeu}$, LeuIGI1 or LeuIGI2. HEK293T cells were co-transfected with plasmids that encode a particular tRNA with wild-type mCherry, an engineered EcLeuRS variant, and EGFP-39-TAG in the presence or absence of 0.2 mM ncAA. The activity of the tRNAs was measured by the expression of EGFP-39-TAG in cell-free extract, normalized relative to wild-type mCherry expression and then reported relative to wild-type EGFP expression.

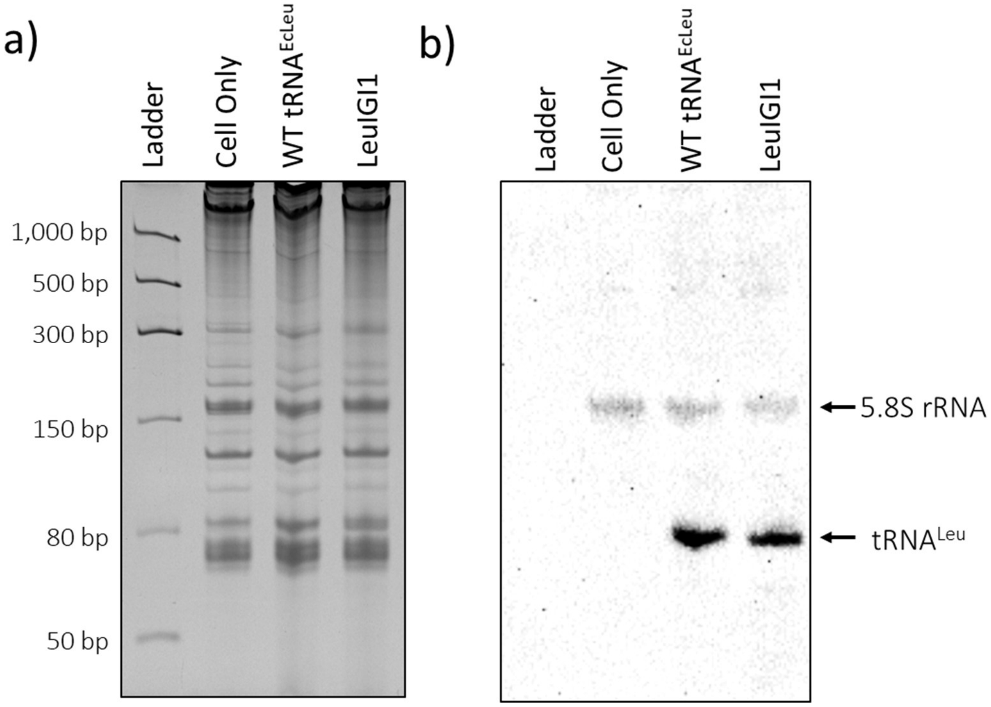


**Figure S7**. Northern blot analysis shows that tRNA$^{EcLeu}$ and LeuIGI1 expresses at similar levels. **a)** 5 µg total RNA from cells transfected with tRNA$^{EcLeu}$ or control cells (untransfected) was resolved in 6.5% acrylamide 8 M urea denaturing gel, then stained with ethidium bromide. **b)** The Northern blot was performed using 20 pM of the tRNA$^{Leu}$ digoxigenin (DIG)-labeled probe, LtR-Vloop-DIG (from LtR-Vloop-NB-R oligonucleotide), and 2.5 pM of control digoxigenin (DIG)-labeled probe, 5.8S-DIG (from 5.8S-NB-RR oligonucleotide). Expected bands for both tRNA$^{Leu}$ (84 bp) and 5.8S ribosomal control (156 bp) noted on the left with an arrow.

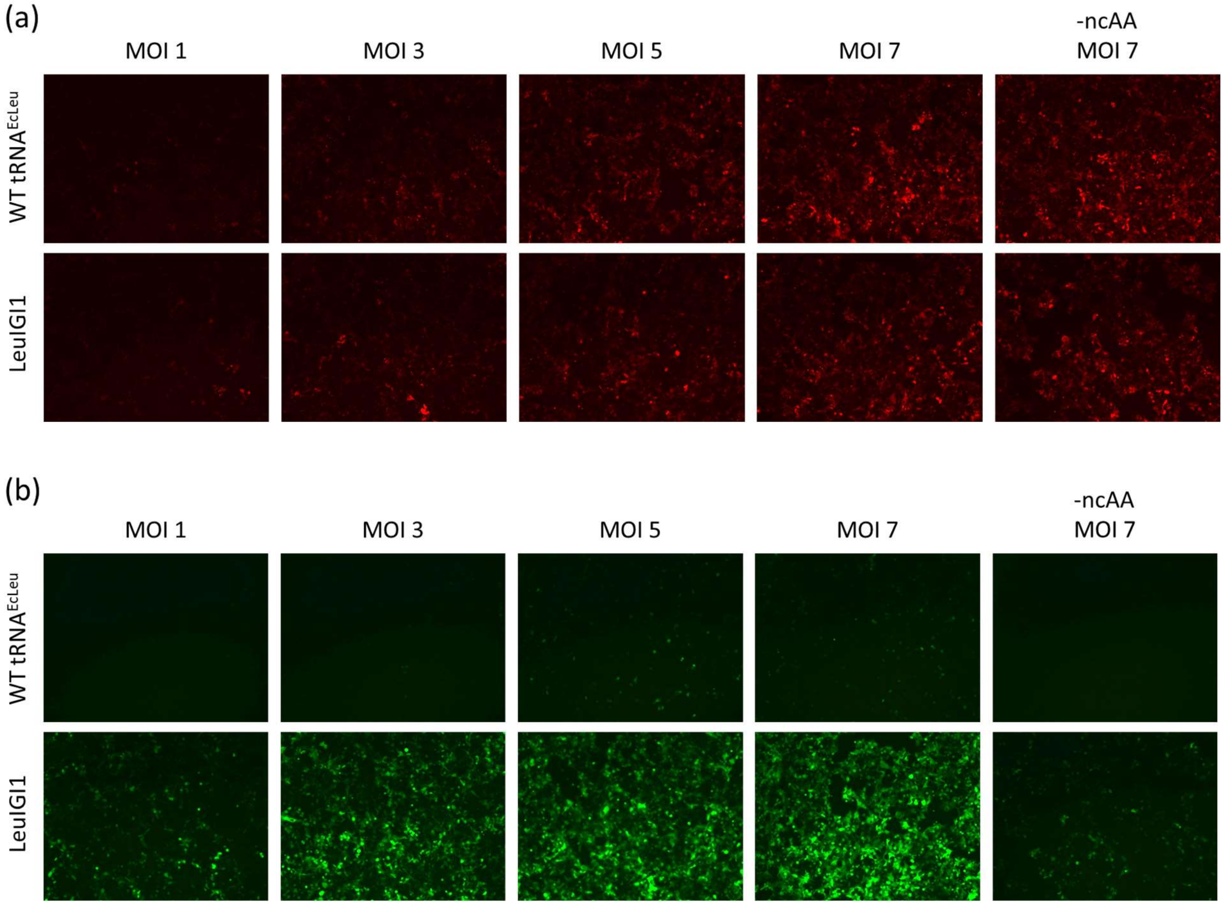


**Figure S8.** Representative fluorescence microscopy images of **(a)** mCherry and **(b)** EGFP-39-TAG expressing cells transduced with the BacMam vectors with increasing MOI of the tRNA-mCherry BacMam.. These experiments were performed using PLRS1 and Cap.

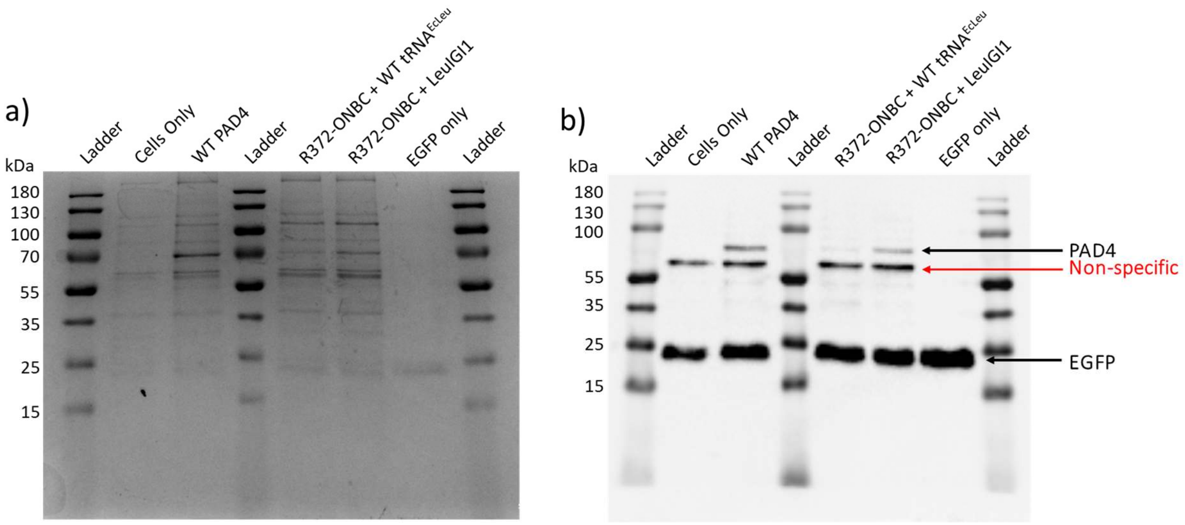


**Figure S9**. LeuIGI1 rescues the expression of PAD4-R372-TAG incorporating photocaged citrulline derivative ONBC (Figure 3c). **a)** Wild-type PAD4, or PAD4-R372-ONBC was expressed in HEK293T cells using the PLRS1/tRNA$^{EcLeu}$ pair. Cell-free extracts from each expression was adjusted to same total protein, and spiked with 400 ng of purified wild-type EGFP with a polyhistidine tag to serve as an internal control during the following Ni-NTA enrichment step. The expressed protein was enriched using Ni-NTA chromatography using a C-terminal polyhistidine tag. After loading, and washing, the histidine-tagged proteins were eluted using 300 mM imidazole-containing buffer. The samples were resolved using SDS-PAGE and stained using Coomassie blue. **(b)** Western blot analysis of the gel from (**a**) using an anti-polyhistidine tag antibody show similar EGFP levels in all samples. PAD4-372-ONBC expression was much lower relative to wild-type PAD4, when expressed using wild-type tRNA$^{EcLeu}$. However, the use of LeuIGI1 significantly enhanced its expression level.

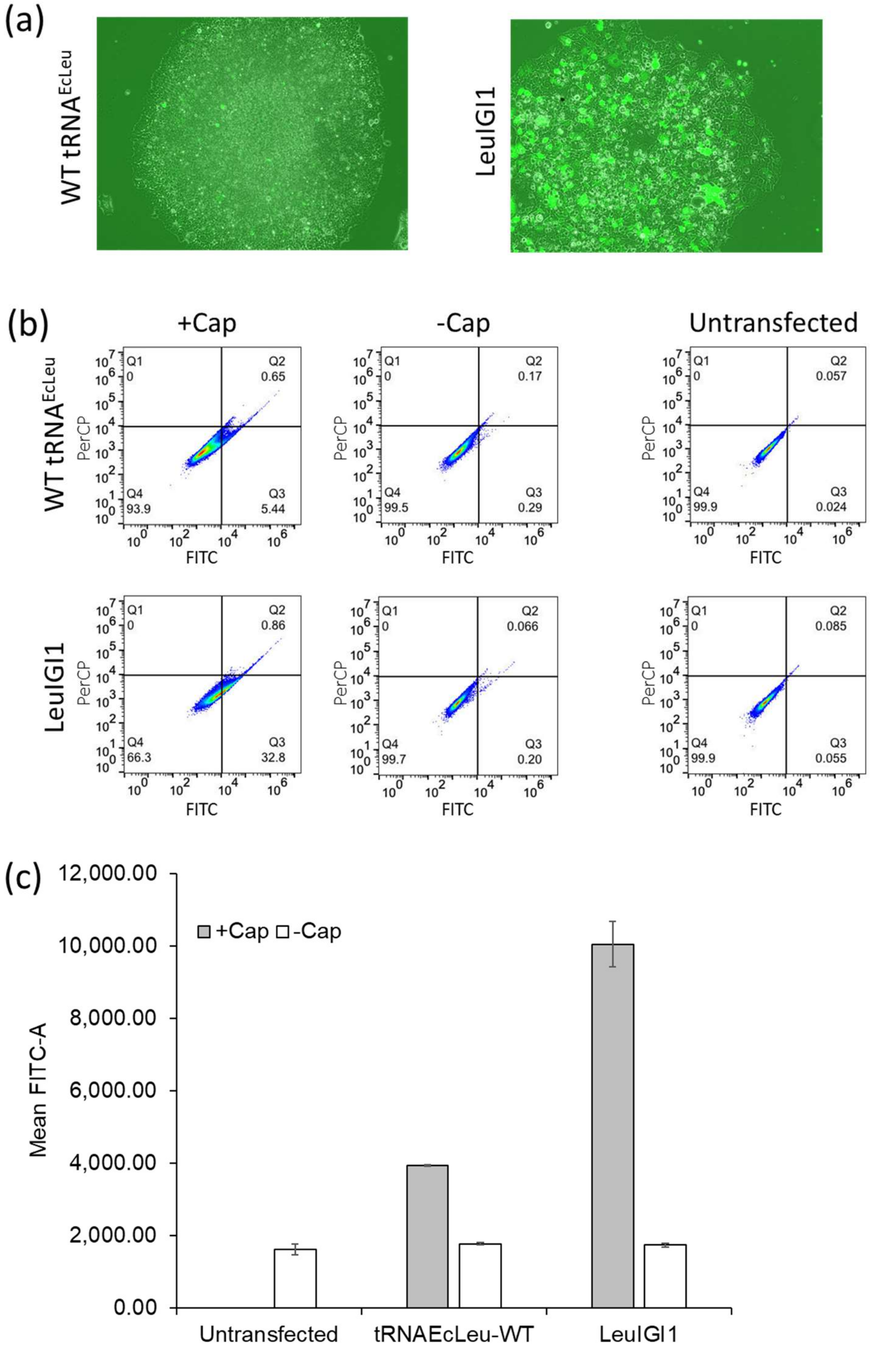


**Figure S10. a)** Representative fluorescence microscopy images of colony formation during stable EBV cell line generation under puromycin selection with EGPF-39-TAG and 4xU6-tRNA$^{EcLeu}$ of either wild-type or LeuIGI1. **b)** FACS analysis of the polyclonal EBV pools in the presence and absence of 1 mM Cap. The LeuIGI1 containing cell pool shows significantly more EGFP-expressing cells in the presence of ncAA than the wild-type tRNA$^{EcLeu}$. **c**) Mean fluorescence intensities of these pools.

### Methods

**Cell culture.**
HEK293T (ATCC) cell cultures were maintained at 37 °C and at 5% $CO_2$ in DMEM-high glucose (HyClone) supplemented with 10% fetal bovine serum (Corning) and 100 U/mL antibiotic antimycotic penicillin/streptomycin/fungiezone (HyClone).

**Noncanonical amino acids.**
Azido-lysine (AzK) was purchased from Iris Biotech GMBH (Germany). Diazirine-lysine (DiazK), strained cyclooctyne-L-lysine (SCOK), and Cyclopropene-L-lysine (CpK) were purchased from Sirius Fine Chemicals (Germany). Aminocaprylic acid (Cap) was obtained from TCI America (Portland, Oregon). Long chain azide (LCA) [1], short chain azide (SCA)[1], cysteine-5-azide (C5Az)[1], o-nitrobenzylcitruline (ONBC)[2], dansyl alanine (DanAla)[3] and photocaged cysteine (Cmn)[4] were synthesized using previously established routes.

**General Cloning.**
All plasmid propagation and cloning were performed in the *E. coli* TOP10 strain grown in LB. Restriction enzymes and T4 DNA ligase were from New England Biolabs (NEB). Gibson assembly was carried out using NEBuilder HiFi DNA Assembly Master Mix (NEB). DNA oligos were purchased from Integrated DNA Technologies (IDT) and Genewiz and are listed in in **Supplementary Table 1.** For ligation cloning, correct size digestion products were purified by agarose gel purification and a 50 ng ligation was performed with at least 5:1 insert to vector ratio. Sanger sequencing was performed by Eton Bioscience and Genewiz.

<u>Hit Cloning</u>
A total of 15 $tRNA^{Leu}$ sequences, seen in **Supplementary Table 2,** were selected from the top enriched or consensus sequences of enriched $tRNA^{EcLeu}$ from the selection and were purchased as oligos from Twist Bioscience. These oligos were 300 bp long that included the desired $tRNA^{EcLeu}$ sequence with flanking BsaI sticky ends designed to be used for Golden Gate Assembly for $tRNA^{Leu}$ insert. The oligos were PCR amplified using PrimeSTAR Max DNA Polymerase (Takara Bio) with forward and reverse primers PytR-Twist-GGA-F and PytR-Twist-GGA-R. After PCR amplification, each tRNA construct, or HTS, was isolated using agarose gel purification and golden gate assembly was performed with pAAV-ITR-mCherry-GGA tRNA library as the receiving vector, using BsaI-HFv2 for scar-less introduction of $tRNA^{Leu}$ HTS.

<u>Additional aaRS cloning</u>
pIDTSmart-CMV plasmids containing previously evolved DanRS or NvocRS were constructed to test incorporation of either DanAla[3] or Cmn[4,5], respectively. For pIDTSmart-CMV-DanRS construction, the DanRS insert underwent PCR with Phusion Hot Start II DNA Polymerase (Thermo Scientific) according to the manufacturer's protocol. DanRS-HindIII-F and DanRS-XhoI-R primers were used to add HindIII and XhoI restriction sites. The DanRS insert was then digested with HindIII and XhoI while the pIDTSmart-CMV-PLRS1 backbone was digested with AgeI, HindIII and XhoI to ensure the correct band to exchange the PLRS1 with DanRS. For pIDTSmart-CMV-NVocRS construction, restriction digestion of pAcBac2-CMV-NVocRS-4xU6-LtR with HindIII and XhoI was performed to remove the NVocRS and insert it into pIDTSmart-

CMV-PLRS1 backbone. The pIDTSmart-CMV-PLRS1 backbone was once again digested with with AgeI, HindIII and XhoI to ensure the correct band to exchange the PLRS1 with NVocRS.

BacMam vector cloning

All PCRs for cloning for BacMam constructs was carried out using Phusion Hot Start II DNA Polymerase (Thermo Scientific) according to the manufacturer's protocol. For pAcBac1-CMV-PLRS1 cloning, PCR amplification PLRS1 was amplified from pIDTSmart-CMV-PLRS1 with CMV-iF and polyLeuRS-ins-EcoRI-R then digested with restriction enzymes EcoRI and NheI to inserted into an empty pAcBac1 vector. For pAcBac2-CMV-PLRS1 with EGFP(wt) or EGFP(39TAG), the EGFP inserts was amplified with RS-pIDT-AvrII-F and RS-pIDT-AvrII-R primers from their respective pAcBac1-CMV-EGFP vector and then digested with AvrII and SpeI and inserted into pAcBac1-CMV-PLRS1 at the single AvrII location. Orientation of the insert was confirmed by analytical digestion and Sanger sequencing. Since EGFP(39TAG) insert has an AvrII cute site at the 39 position, partial digestion of AvrII was performed. For tRNA-mCherry-BacMam, pAcBac vectors had MTH-mCherry and either the wild-type $tRNA^{Leu}$ or the highly active LeuIGI1 (HTS-9 $tRNA^{Leu}$). First, pAcBac1-1xU6-LtR-wt or pAcBac1-1xU6-LeuIGI1 were constructed from PCR of tRNA in either pAAV-ITR-CMV-mCherry-1xU6-LtR-wt or pAAV-ITR-CMV-mCherry-1xU6-LeuIGI1 (HTS-9 $tRNA^{Leu}$) with AvrII-ins-KpnI-F and PytR-Twist-GGA-R. The tRNA insert were digested with AvrII and NheI and inserted into an empty pAcBac1 vector single AvrII location. MTH-mCherry insert was added by digestion of pAcBac1-MTH-mCherry and pAcBac1-1xU6-LtR-wt or pAcBac1-1xU6-LeuIGI1 with SbfI and NotI.

Eppstein–Barr virus (EBV) plasmid cloning

EBV plasmid constructs were made using 4 fragment Gibson Assembly, to include the aaRS, EGFP(39TAG)-P2A-Puro-IRES cassette[6], the tRNA of interest, and the EBV plasmid backbone.[6] The first fragment was PCR amplified aaRS from pIDTSmart-CMV-PLRS1 using RS-PCR-F and RS-PCR-R. The second fragment was PCR amplified EGFP-puromycin from pEBV1 with EGFP-PCR-F and EGFP-PCR-R. The third fragment was PCR amplified from pIDTSmart-4xU6-LtR, which contain 4 copies of either wild-type tRNA or LeuIGI1 and was PCR amplified using primers tRNA-PCR-F and tRNA-PCR-R. The last fragment was generated by restriction enzyme digestion of pEBV1 plasmid with EcoNI and KpnI. Before Gibson assembly each PCR fragment underwent DpnI digestion and were purified by agarose gel extraction. Gibson assembly was performed per manufacturer's instructions in 1:1:1:1 ratio with 0.50 pmol DNA total. PCR fragments for the tRNA cassette were performed with PrimeSTAR Max DNA Polymerase (Takara Bio) and while the EGFP(39TAG)-P2A-Puro-IRES cassette were performed with Phusion Hot Start II DNA Polymerase (Thermo Scientific) according to the manufacturer's protocols.

PAD4 plasmid cloning

Final PAD4 plasmids were made by digesting the previously made[2] starting vectors of pAcBac3-CMV-PLRS1-CAG-PAD4-wt-12xHisTag or pAcBac3-CMV-PLRS1-CAG-PAD4-R372TAG-12xHisTag with AvrII only and inserting 4 copies of either wild-type tRNA or LeuIGI1 from AvrII and NheI digestion of pIDTSmart-4xU6-LtR plasmids.

**Leucyl tRNA library design and plasmid construction.**

The sequences of all known bacterial leucyl tRNA ($tRNA^{Leu}$) was obtain from transfer RNA database (http://trnadb.bioinf.uni-leipzig.de/DataOutput/Search). Of the 688 $tRNA^{Leu}$, two sequences with 8-base pair acceptor stem and four sequences with 6-base pair acceptor stem were excluded from library design. A Weblogo was created with remaining 682 7-base pair acceptor stem $tRNA^{Leu}$ sequences, and an acceptor stem base pair library was designed based on the consensus from these results (https://weblogo.threeplusone.com/create.cgi). Based on the library design, a custom program (Twist oligo program) was run to make a list 7,776 sequences of 150 base pooled oligos to be order from Twist Bioscience. Each oligo was a unique library member with flanking BsaI sites designed to be used for Golden Gate Assembly for tRNA library member insertion. The Twist oligo pools were PCR amplified with PytR-Twist-GGA-F and PytR-Twist-GGA-R forward and reverse primers. The PCR was performed at a low cycle number (15 cycles) and high-fidelity DNA polymerase PrimeSTAR Max DNA Polymerase (Takara Bio) to prevent scrambling of library members. Large scale golden gate assembly was performed with ~1 µg library vector pAAV-ITR-CMV-mCherry-GGA tRNA Library Vector, using BsaI-HFv2 for scar-less introduction of tRNA library members. Golden gate assembly reactions were concentrated by ethanol precipitation with yeast tRNA (Ambion) and transformed into electrocompetent TOP10 *E. coli*. >$1.4 \times 10^6$ transformants were plated (>100-fold library coverage). These colonies were pooled and their DNA was miniprepped for packaging into AAV.

**AAV2 Production and Infective Titer of mock and library tRNAs into AAV (wild-type capsid)**

Various cargo was packaged as previously described.[7-9] In brief, 8 µg each of the appropriate cargo plasmid (pAAV-ITR-tRNA-fluorescent protein), pHelper, and pAAV-RC2 were transfected in a 10 cm tissue culture dish with 84 µL of polyethylenimine (PEI, Sigma, 1 mg/mL) in 420 µL DMEM (no FBS) and incubated for 10 min. Media was exchanged for fresh DMEM 24 hours after transfection and then AAV semi-purified using PEG precipitation 72 hours after transfection. Infectious titer was calculated from flow cytometry at a low MOI of infected cells as previously described.[7] A 12-well plate was seeded at 0.7 million cells per well with HEK293T cells and infected at confluence the next day. Infectivity was boosted by the addition of a total of 5 mM sodium butyrate (Sigma). Two days post infection cells were trypsinized, washed with PBS and then analyzed using flow cytometry to count the fluorescent population. Flow cytometry was performed on S3e cell sorter (Bio-Rad).

**Virus-Assisted Directed Evolution of tRNA (VADER).**

VADER was performed as previously described[7] with the $tRNA^{Leu}$ library and pIDTSMART-CMV-PLRS1-AAV-RC2-TAG454 as the selection pressure plasmid at an apparent MOI of 2 (the actual MOI is substantially reduced in the presence of PEI, the transfection reagent) in three separate HEK293T 10 cm plates. After selection, viral DNA was recovered and then amplified as described.[7] The amplified tRNA from the viral was cloned into pAAV-ITR-mCherry-no-tRNA-GGA vector using a large-scale Golden Gate Assembly as described in library cloning method.

**Production of AAV RC2-454-ncAA**

The incorporation of ncAA to site T454TAG of AAV2 capsid was tested in 12 well-plates as previously described.[10] In brief, 1.5 µg/per well plasmids containing AdHelper genes, AAV genes, and an PLRS1/tRNA pair were mixed with polyethyleneimine (Sigma) in serum-free media

(DMEM, HyClone) at RT. After 15 minutes of incubation, the mixture was added to HEK293T cells at 70% confluency. NcAA (AzK (H-L-Lys(EO-N3)-OH, Iris biotech GMBH) or LCA was added to the final concentration of 0.2 mM for the production of AAV2-containing ncAA. Viruses were harvested 72 h post-transfection using the AAVPro Extraction Solution Kit (Takara) and titered using the AAVpro® Titration Kit (for Real-Time PCR) Ver.2 (Takara), following the manufacturer's instructions.

**Mock selections using LeucyltR-mCherry and PytR-GFP**

The mock selection was performed as described in VADER protocol with 1:1,000 mixture of active to inactive virus made from pAAV-ITR-mCherry-1xU6-LtR-wt(TAG) and pAAV-ITR-EGFP-1xU6-PytR-wt(TAG), respectively. Enrichment of active virus was analyzed by mCherry and GFP fluorescence via flow cytometry to determine red to green ratio. Cells were infected in 12-well plate as described in virus titer protocol with 0.5 μL input virus, 50 μL of positive selection virus, and 250 μL of negative selection virus. Flow cytometry was performed on S3e cell sorter (Bio-Rad).

**Illumina sample preparation, and high throughput sequencing**

Illumina sequencing sample preparation was done for the library input and outputs DNA from replicate selections using two rounds of PCR with PrimeSTAR Max DNA Polymerase for the addition of Illumina adapters based on sequences of TruSeq DNA HT Sample Prep Kit (Illumina). The first round of PCR had forward and reverse primers which contained primer-binding regions flanking the tRNA, diversity sequence bases and half of the TruSeq adapters. The diversity sequence for these primers was used for cluster identification during Illumina sequencing runs. The forward primer was made by combining four primers in 1:1:1:1 fashion with 9 base sequence variation for base sequence diversity (Illumina-PytR-X1a-F, Illumina-PytR-X1b-F, Illumina-PytR-X1c-F, Illumina-PytR-X1d-F = ACACTCTTTCCCTACACGACG CTCTTCCGATCT[X1a-d)]TTATATATCTTGTGGAAAGGACGAAAC). The reverse primer was combined in a similar fashion to allow 7 base variation for diversity (Illumina-PytR-X2a-R, Illumina-PytR-X2b-R, Illumina-PytR-X2c-R, Illumina-PytR-X2d-R = GTGACTGGAGTTCAG ACGTGTGCTCTTCCGATC[X2a-d]GCTCTCGTCGATCCGCTAGC). Products from the first round were then purified by agarose gel extraction and then underwent another round of PCR for the addition of the full TruSeq adapter with different barcodes for Illumina sequencing. The forward primer series of Illumina-i5-F (AATGATACGGCGACCACCGAGATCTACAC [i5]ACACTCTTTCCCTACACGACGC) variants contained different barcodes and a reverse primer Illumina-i7-R (CAAGCAGAAGACGGCATACGAGAT[D702]GTGACTGGAGTTCA GACGTGTGCTC) barcode variant. Both i5 and i7 barcode sequences are given in **Supplemental Table 3**. Unique combinations of i5 and i7 barcode sequences were applied to each sample to enable multiplex sequencing with only forward reads were used in data analysis. Samples were prepared for sequencing using the 300-cycle MiSeq Reagent Kit v2 (Illumina) per manufacturer's instructions and sequence on an Illumina MiSeq System with 25% Illumina PhiX control added.

**Illumina high-throughput-sequencing data processing**

Processing of sequencing data generated by the MiSeq System was performed as previously described[7] with a custom Python code available on GitHub.[11] Reads containing any bases with Q-scores lower than 30, corresponding to an error rate of 0.1%, were discarded. Besides Q-score filtering, the reads were filtered by "mismatch filter" that compared each read to the expected tRNA sequence within the fixed regions of the tRNA. Additionally, the minimum abundant library count was set to 1 with three acceptor stem library members (GGGGGUG/CACCCCC, GCCCUUA/UGAGGGC, and GCAAAUA/UAUUUGU) discarded for being too low in abundance in input library. A comma-separated file was created for all sequences that passed the filtering and counts for each sample were tallied to determine "fraction of total", "fold enrichment", and "enrichment factor". "Fraction of total" was calculated by dividing the counts of a given sequence by the total counts of all sequences in that sample, "fold enrichment" was calculated the ratio of counts in the selection output to counts in the selection input, and fold enrichment normalized to the most enriched hit, and "enrichment factor" was calculated by dividing the fold enrichment of a given hit by the fold enrichment of the most enriched hit in that sample. After these calculations each library member was sorted and ranked by the average enrichment factor that was determined from the two output enrichment factor values.

**Hit Activity and substrate scope**

Initial hit analysis was conducted by transfecting HEK293T cells in 12-well plates with 0.5 µg each of a constructed hit pAAV-ITR-CMV-mCherry-1xU6-LtR-HTS plasmid, pIDTSmart-CMV-PLRS1, and pAcBac1- CMV-EGFP(39TAG) added in the presence and absence of 1 mM Cap. Transfections were done with polyethylenimine (PEI, Sigma Aldrich NC0704680) and DNA which was mixed at a ratio of 3.5 µL PEI (1 mg/mL) to 1 µg DNA in DMEM (17.5 µL per 1 µg of DNA), and incubated for 10 min. Fluorescence images and fluorescence protein expression analysis were performed two days post transfection. Images of each well were taken on Zeiss Axio Observer A1 microscope on brightfield, GFP and mCherry channels. Fluorescence analysis was obtained for both EGFP and mCherry by removal of media and cell lysis in 300 µL with CelLytic M buffer (Sigma) with 0.01 µL/mL Pierce universal nuclease (Fisher). Lysis solution was then incubated for 10 min while nutating and lysate was transferred into a clear bottom 96-well assay plate (Corning, Corning, NY). EGFP fluorescence was recorded at 480 nm excitation and 520 nm emission while mCherry fluorescence was recorded using 580 nm excitation and 620 nm emission using Molecular Devices SpectraMax M5 microplate reader (Molecular Devices, Sunnyvale, CA). The fluorescence values for untransfected wells were treated as background level fluorescence and was subtracted from each well. The each EGFP value were normalized to mCherry internal control.

The best $tRNA^{Leu}$ hit, HTS-9 or LeuIGI1 (GCCCGUA/ UGCGGGC), was selected for further activity testing with a variety of substrates that include AzK, CpK, C5Az, DiazK, LCA, ONBC, SCA, SCOK with PLRS1. *E. coli* leucyl specific aaRS DanRS and NVocRS, which were evolved to incorporate DanAla[3,5] and photocage cysteine[4], Cmn, respectively were also tested. An additional LeuIGI mutant, HTS-1 or LeuIGI2 (GGGCGUG/CGCGCCC) activity was tested for AzK, DanAla and Cmn, HEK293T cells were transfected using the same method as hit activity testing at 70-80% confluence with 0.5 µg pAAV-ITR-CMV-mCherry-LtR containing either the wild-type or evolved tRNA, 0.5 µg pIDTSmart-CMV-aaRS, and 0.5 µg pAcBac1-CMV-EGFP(39TAG) in a 12-well plate. Analysis of improved activity of LeuIGI1 and LeuIGI2 was assessed using same method as hit activity testing with a BioTek Synergy Neo2 Hybrid Multimode microplate reader.

**Baculovirus Production and Infection for LeuIGI1 characterization**

Vesicular stomatitis virus glycoprotein-pseudotyped (VSV-G) baculovirus vectors (BacMam) were generated and titered as previously described.[12,13] Sf9 cells (Life Technologies) that were maintained in a suspension culture at 28 °C in Sf-900 III SFM serum-free media (Fisher Scientific). BacPAK Baculovirus Rapid Titer Kit (Clontech) was used for titers per manufacturer's instructions for BacMam vectors, or baculovirus, containing PLRS1 with either wild-type EGFP or EGFP(39TAG). An infective HEK293T titer was used to determine for the tRNA-mCherry baculovirus by transduction of confluent HEKL293T cells, seeded in a 12-well plates at 0.7 million cells per well one day prior to transduction. The tRNA-mCherry baculoviruses were diluted to achieve low MOI conditions and 5 mM sodium butyrate was added to boost infectivity. Two days post transduction the cells were trypsinized, washed with PBS, and analyzed by flow cytometry to determine infectivity titer by the population count of mCherry fluorescence. Flow cytometry was performed on S3e cell sorter (Bio-Rad).

For tRNA activity testing HEK293T cells were seeded at 0.4 million cells per well in a 12-well plate. The next day the cells were co-transduced with EGFP reporter-PLRS1 and mCherry-tRNA baculovirus in triplicates at 30% confluence. All cells received 100 MOI of EGFP reporter-PLRS1 baculovirus and a range of 1-7 MOI HEK293T infectious titer of baculovirus vectors encoding mCherry-tRNA. 1 mM Cap was added dropwise at all tRNA MOI ranges. The highest infectious titer range on MOI of 7 also include a mCherry-tRNA control in the absence of ncAA. Two days post transduction, images of mCherry and GFP were taken and activity of tRNA was measures by fluorescence of EGFP and mCherry, as done with hit testing. EGFP expression was normalized to the EGFP value of cells transduced PLRS1-EGFP(wt) baculovirus at 100 MOI.

RNA isolation

HEK293T cells were seeded at 7.5 million in a 10 cm tissue culture dish and then transfected at 75% confluency with 24 µg of tRNA variant-containing pAAV-ITR-LtR-mCherry plasmid using polyethylenimine (PEI) (1 mg/mL). The day after transfection, culture media was exchanged for fresh DMEM. Two days (48 hours) after transfections, cells were harvested and RNA isolation with TRIzol Reagent (Thermo Fisher) following manufacturer's instructions was performed. The total RNA concentration was determined via Nanodrop Spectrophotometer. RNA integrity was assessed by A260/A280 value, as well as the presence of distinct, intact 28S and 18S ribosomal RNA bands on 1% agarose gel.

Northern blot probe preparation

Oligonucleotides design to bind to leucyl tRNA were 3'-end labeled with DIG using the DIG Oligonucleotide 3′-End Labeling Kit, 2nd Generation (Roche), and labeling efficiency was measured following manufacturer's instructions. The oligonucleotides used was LtR-Vloop-NB-R, which binds to the variable loop of the tRNA in order to be compatible with all A-stem variations, as well as 5.8S-NB-R probe to bind to 5.8S RNA as a positive control to assess overall RNA concentration. The sequences of the oligonucleotides can be found in **Supplemental Table 1**.

Northern blotting

A modified version sensitive non-isotopic northern blot[14] was performed as previously described[7] with 20 pM of the $tRNA^{Leu}$ digoxigenin (DIG)-labeled probe, LtR-Vloop-DIG (from LtR-Vloop-NB-R oligonucleotide), and 2.5 pM of control digoxigenin (DIG)-labeled probe, 5.8S-DIG (from 5.8S-NB-RR oligonucleotide).

**Puromycin selection and FACS analysis of EBV-based stable cell line.**
Two 10 cm tissue culture dishes, seeded at 7.5 million of HEK293T cells. The next day the cells were transfected at 70-80% confluence with 24 µg of EBV-EGFP-puromycin plasmid that contain either four copies of the wild-type $tRNA^{Leu}$ or LeuIGI1, 84 µL polyethylenimine (PEI) (1mg/mL) and 1 mM Cap. Media was changed with fresh DMEM media, 1 mM Cap and 3 µg/mL puromycin two days post transfection and then every day for 7 days during initial cell line expansion. Following the 7 days of continuous media change, fresh DMEM with 1 mM Cap and 3 µg/mL puromycin were changed every two days until only large cluster colonies with EGFP florescence cells formed, indicating cells with episomal maintenance of EBV plasmid for stable cell line. EBV stable cells were then trypsinized and added to T25 flask for larger production of stable cell line. The day after T25 transfer, fresh media with 1 mM Cap and 3 µg/mL puromycin was added, and media change continued every two days until cell confluence and then transferred to T125 flask. The day after T125 transfer, fresh media with 1 mM Cap and 3 µg/mL puromycin was added, and media change continued every two days until confluence. Confluent stable cell lines were then trypsized and underwent flow cytometry on S3e cell sorter (Bio-Rad) to determine EGFP fluorescence with and without the presence of Cap or were frozen for storage of stable cell line. Images of brightfield and EGFP fluorescence was taken before each media change, with EBV stable cell line of wild-type tRNA taking 47 days and LeuIGI1 EBV stable cell line taking 43 days post transfection media change.

**Protein arginine deiminase 4 (PAD4) Protein Purification and Western Blot**
7.5 million HEK293T cells/plate were seeded in 10 cm cell culture dishes, and then transfected in duplicates the next day at 70-80% confluence with 10 µg of either pAcBac1-EGFP-12xHisTag, pAcBac3-CMV-PLRS1-4xU6-LtR-wt-CAG-PAD4-R372TAG-12xHisTag, or pAcBac3-CMV-PLRS1-4xU6-LeuIGI1-CAG-PAD4-R372TAG-12xHisTag. To increase protein production, plasmids were transfected with 50 µL polyethylenimine (PEI) MAX and a total of 15 mM sodium butyrate. For the pAcBac3 containing tRNA, 1 mM of O-Nitrobenzoylcitruline (ONBC) was added during transfection. Two days after transfection, the media was removed from plate and cells were scraped off culture dishes with 2 mL 1x PBS. The cells were then spun down at 4,000 x g for 5 min and supernatant was discarded with the cell pellet stored at
-80 °C until purification. For purification, the duplicate cell pellets were combined and lysed with 600 µL cell lysis buffer containing celLytic M, protease inhibitor and universal nuclease solution at 10,000: 100: 1 ratio, and then nutated for 20 minutes. The cells were then centrifuged for 10 minutes 18,000 x g and supernatant of the proteins from total cell lysate was transferred to new eppies. Each PAD4-ONBC tRNA sample was normalized to lowest concentration, and 2x equilibrium buffer containing 20 mM $Na_2HPO_4$, 300 mM NaCl, pH 7.4, 25 mM Imidazole, pH 7.4, was added. Additionally, 400 ng of Ni-NTA nickel column purified EGFP (400 ng/µL) was spiked in as an internal control for Ni-NTA nickel resin (Thermo Scientific) pulldown. PAD4-ONBC tRNA and spiked EGFP samples were allowed to nutate for 1 hour at 4°C with 50 µL prepared nickel beads and then spun down 5,000 x g for 5 minutes to remove supernatant. Protein Ni-NTA nickel resin were then washed five times with 500 µL wash buffer (20 mM $Na_2HPO_4$, 300 mM NaCl, 50 mM Imidazole, pH 7.4), vortexed, and spun down at 5,000 x g for 5 minutes to remove supernatant. Protein was then eluted off the beads with 1x elution buffer (20 mM $Na_2HPO_4$, 300 mM NaCl, 300 mM Imidazole, pH 7.4), spun down at 5,000 x g for 5 minutes, and entire supernatant was collected and used to run western blot analysis using 6x-His Tag Monoclonal Antibody (Invitrogen).

**Supplementary Table S1. Primers used for PCR**

| Primer Name | Sequence |
|---|---|
| 5.8S-NB-R | CGCAAGTGCGTTCGAAGTGTCGATGATCAATGTG |
| AvrII-ins-KpnI-F | TTATTTACCTAGGGGTACCTCGGGCAGGAAGAGGGCC TATTTCCCATG |
| CMV-iF | GTAGGCGTGTACGGTGGGAGGTCTATATAAG |
| DanRS-HindIII-F | TAATTAAAGCTTGCCGCCACCATGGAAGAGCAATACCG CCCGG |
| DanRS-XhoI-R | TAATAACTCGAGTTAAACGGGCCCGCCAACGAC |
| EGFP-PCR-F | CCGGGCGGTATTGCTCTTCCATTCCGGACGCCATGGT TGTGG |
| EGFP-PCR-R | GCTGACTAGATAAACTGGCCGTCGTTTTACTCTAGAAT AGTAATCAATTACGGGGTCATTAGTTCATAGCCC |
| Illumina-i5-F | AATGATACGGCGACCACCGAGATCTACAC[D501]ACAC TCTTTCCCTACACGACGC, AATGATACGGCGACCACCGAGATCTACAC[D502]ACAC TCTTTCCCTACACGACGC, AATGATACGGCGACCACCGAGATCTACAC[D503]ACAC TCTTTCCCTACACGACGC |
| Illumina-i7-R | CAAGCAGAAGACGGCATACGAGAT[D702]GTGACTGGA GTTCAGACGTGTGCTC |
| Illumina-PytR-X1a-F | ACACTCTTTCCCTACACGACGCTCTTCCGATCTggaactc ctTTATATATCTTGTGGAAAGGACGAAAC |
| Illumina-PytR-X1b-F | ACACTCTTTCCCTACACGACGCTCTTCCGATCTtagcacat gTTATATATCTTGTGGAAAGGACGAAAC |
| Illumina-PytR-X1c-F | ACACTCTTTCCCTACACGACGCTCTTCCGATCTccttgagg aTTATATATCTTGTGGAAAGGACGAAAC |
| Illumina-PytR-X1d-F | ACACTCTTTCCCTACACGACGCTCTTCCGATCTatcgtgta cTTATATATCTTGTGGAAAGGACGAAAC |
| Illumina-PytR-X2a-R | GTGACTGGAGTTCAGACGTGTGCTCTTCCGATCgactacc GCTCTCGTCGATCCGCTAGC |
| Illumina-PytR-X2b-R | GTGACTGGAGTTCAGACGTGTGCTCTTCCGATCtctggat GCTCTCGTCGATCCGCTAGC |
| Illumina-PytR-X2c-R | TGACTGGAGTTCAGACGTGTGCTCTTCCGATCctgatggG CTCTCGTCGATCCGCTAGC |
| Illumina-PytR-X2d-R | GTGACTGGAGTTCAGACGTGTGCTCTTCCGATCagacct aGCTCTCGTCGATCCGCTAGC |
| LtR-Vloop-NB-R | CAGCGCGAACGCCGAGGGATTTAG |
| polyLeuRS-ins-EcoRI-R | TTATTTAGAATTCATTAAACGGGCCCGCCAACGAC |
| PytR-Twist-GGA-F | TTATTTAGGTCTCTTGTGGAAAGGACGAAACACC |
| PytR-Twist-GGA-R | TAAATAAGGTCTCGCTGCTCTCGTCGATCCGCTAGC |
| RS-PCR-F | CTTGGCAGTACATCTACGTATTCGAGCTCGGTACCATC ACCGCCTCAGAAGCCATAGAGC |

| | |
|---|---|
| RS-PCR-R | CCACAACCATGGCGTCCGGAATGGAAGAGCAATACCGCCCGG |
| RS-pIDT-AvrII-F | TTATTTACCTAGGGCGGCCGCAAATACCTGCAGGATCC |
| RS-pIDT-AvrII-R | TAAATAACCTAGGGTTCTTTCCGCCTCAGAAGCCATAGAGCC |
| tRNA-PCR-F | GGGCTATGAACTAATGACCCCGTAATTGATTACTATTCTAGAGTAAAACGACGGCCAGTTTATCTAGTCAG |
| tRNA-PCR-R | CTCGGGACCCCTCCTCTTCCTCTTCAAGGGAGCCTCAGGAAACAGCTATGACATC |

**Supplementary Table S2. tRNA$^{Leu}$ HTS Sequences**

| **LeucyltRNA** | **Acceptor Stem Sequence** | **Full tRNA sequence** | **tRNA Construct** |
|---|---|---|---|
| Wild type | GCCCGGA/TCCGGGT | **GCCCGGA**TGGTGGAATCGGTAGACACAAGGGATTCTAAATCCCTCGGCGTTCGCGCTGTGCGGGTTCAAGTCCCGC**TCCGGGT**A | Wild-type |
| HTS-1 (**LeuIGI2**) | GGGCGTG/CGCGCCC | **GGGCGTG**TGGTGGAATCGGTAGACACAAGGGATTCTAAATCCCTCGGCGTTCGCGCTGTGCGGGTTCAAGTCCCGC**CGCGCCC**A | Consensus |
| HTS-2 | GGGCGCG/CGCGCCC | **GGGCGCG**TGGTGGAATCGGTAGACACAAGGGATTCTAAATCCCTCGGCGTTCGCGCTGTGCGGGTTCAAGTCCCGC**CGCGCCC**A | |
| HTS-3 | GGGCATG/CATGCCC | **GGGCATG**TGGTGGAATCGGTAGACACAAGGGATTCTAAATCCCTCGGCGTTCGCGCTGTGCGGGTTCAAGTCCCGC**CATGCCC**A | |
| HTS-4 | GGGCACG/CGTGCCC | **GGGCACG**TGGTGGAATCGGTAGACACAAGGGATTCTAAATCCCTCGGCGTTCGCGCTGTGCGGGTTCAAGTCCCGC**CGTGCCC**A | |
| HTS-5 | GGGGGTG/CGCCCCC | **GGGGGTG**TGGTGGAATCGGTAGACACAAGGGATTCTAAATCCCTCGGCGTTCGCGCTGTGCGGGTTCAAGTCCCGC**CGCCCCC**A | |
| HTS-6 | GGGGGCG/CGTCCCC | **GGGGGCG**TGGTGGAATCGGTAGACACAAGGGATTCTAAATCCCTCGGCGTTCGCGCTGTGCGGGTTCAAGTCCCGC**CGTCCCC**A | |
| HTS-7 | GGGGATG/CGTCCCC | **GGGGATG**TGGTGGAATCGGTAGACACAAGGGATTCTAAATCCCTCGGCGTTCGCGCTGTGCGGGTTCAAGTCCCGC**CGTCCCC**A | |
| HTS-8 | GGGGACG/CGTCCCC | **GGGGACG**TGGTGGAATCGGTAGACACAAGGGATTCTAAATCCCTCGGCGTTCGCGCTGTGCGGGTTCAAGTCCCGC**CGTCCCC**A | |

| | | | |
|---|---|---|---|
| HTS-9 (**LeuIGI1**) | GCCCGTA / TGCGGGC | **GCCCGTA**TGGTGGAATCGGTAGACACAAGGGATTCTAAATCCCTCGGCGTTCGCGCTGTGCGGGTTCAAGTCCCGC**TGCGGGC**A | High Enrichment |
| HTS-10 | GGGATAG / CTATCCC | **GGGATAG**TGGTGGAATCGGTAGACACAAGGGATTCTAAATCCCTCGGCGTTCGCGCTGTGCGGGTTCAAGTCCCGC**CTATCCC**A | |
| HTS-11 | GGGCATG/ CGTGCCC | **GGGCATG**TGGTGGAATCGGTAGACACAAGGGATTCTAAATCCCTCGGCGTTCGCGCTGTGCGGGTTCAAGTCCCGC**CGTGCCC**A | |
| HTS-12 | GGGCAGA/ TCTGCCC | **GGGCAGA**TGGTGGAATCGGTAGACACAAGGGATTCTAAATCCCTCGGCGTTCGCGCTGTGCGGGTTCAAGTCCCGC**TCTGCCC**A | |
| HTS-13 | GGGCGTA/ TGCGCCC | **GGGCGTA**TGGTGGAATCGGTAGACACAAGGGATTCTAAATCCCTCGGCGTTCGCGCTGTGCGGGTTCAAGTCCCGC**TGCGCCC**A | |
| HTS-14 | GGGCAAG/ CTTGCCC | **GGGCAAG**TGGTGGAATCGGTAGACACAAGGGATTCTAAATCCCTCGGCGTTCGCGCTGTGCGGGTTCAAGTCCCGC**CTTGCCC**A | |
| HTS-15 | GCACACA / TGTGTGC | **GCACACA**TGGTGGAATCGGTAGACACAAGGGATTCTAAATCCCTCGGCGTTCGCGCTGTGCGGGTTCAAGTCCCGC**TGTGTGC**A | |

**Supplementary Table S3. Barcodes for Illumina Sequencing**

| | **Illumina Barcode** | **Sequence** |
|---|---|---|
| i5 | D501 | TATAGCCT |
| | D502 | ATAGAGGC |
| | D503 | CCTATCCT |
| i7 | D702 | TCTCCGGA |

**Supplemental information references.**


1. Zheng Y, Mukherjee R, Chin MA, Igo P, Gilgenast MJ, Chatterjee A. Expanding the Scope of Single- and Double-Noncanonical Amino Acid Mutagenesis in Mammalian Cells Using Orthogonal Polyspecific Leucyl-tRNA Synthetases. *Biochemistry.* **2018** 57(4):441-445. doi: 10.1021/acs.biochem.7b00952,

2. Mondal S, Wang S, Zheng Y, Sen S, Chatterjee A, Thompson PR. Site-specific incorporation of citrulline into proteins in mammalian cells. *Nat Commun*. **2021** 12(1):45. doi: 10.1038/s41467-020-20279-w.
3. Wang J, Xie J, Schultz PG. A genetically encoded fluorescent amino acid. *J Am Chem Soc.* **2006** 28(27):8738-9. doi: 10.1021/ja062666k. PMID: 16819861.
4. Kang JY, Kawaguchi D, Coin I, Xiang Z, O'Leary DD, Slesinger PA, Wang L. In vivo expression of a light-activatable potassium channel using unnatural amino acids. *Neuron.* **2013** 80(2):358-70. doi: 10.1016/j.neuron.2013.08.016.
5. Lemke EA, Summerer D, Geierstanger BH, Brittain SM, Schultz PG. Control of protein phosphorylation with a genetically encoded photocaged amino acid. *Nat Chem Biol.* **2007** 3(12):769-72. doi: 10.1038/nchembio.2007.44.
6. Shao S, Koh M, Schultz PG. Expanding the genetic code of the human hematopoietic system. *Proc Natl Acad Sci U S A*. **2020** 117(16):8845-8849. doi: 10.1073/pnas.1914408117.
7. Jewel D, Kelemen RE, Huang RL, Zhu Z, Sundaresh B, Cao X, Malley K, Huang Z, Pasha M, Anthony J, van Opijnen T, Chatterjee A. Virus-assisted directed evolution of enhanced suppressor tRNAs in mammalian cells. *Nat Methods*. **2023** 20(1):95-103. doi: 10.1038/s41592-022-01706-w.
8. Kelemen RE, Mukherjee R, Cao X, Erickson SB, Zheng Y, Chatterjee A. A Precise Chemical Strategy to Alter the Receptor Specificity of the Adeno-Associated Virus. *Angew Chem Int Ed Engl*. 2**016** 55(36):10645-9. doi: 10.1002/anie.201604067.
9. Kelemen RE, Erickson SB, Chatterjee A. Production and Chemoselective Modification of Adeno-Associated Virus Site-Specifically Incorporating an Unnatural Amino Acid Residue into Its Capsid. *Methods Mol Biol*. **2018** 1728:313-326. doi: 10.1007/978-1-4939-7574-7_20.
10. Erickson SB, Pham Q, Cao X, Glicksman J, Kelemen RE, Shahraeini SS, Bodkin S, Kiyam Z, Chatterjee A. Precise Manipulation of the Site and Stoichiometry of Capsid Modification Enables Optimization of Functional Adeno-Associated Virus Conjugates. *Bioconjug Chem.* **2023** doi: 10.1021/acs.bioconjchem.3c00411.
11. Kelemen, R. VADER processing pipeline (version 1.0) http:github.com/chatterjeelab2022/VADER (2018).
12. Zheng Y, Lewis TL Jr, Igo P, Polleux F, Chatterjee A. Virus-Enabled Optimization and Delivery of the Genetic Machinery for Efficient Unnatural Amino Acid Mutagenesis in Mammalian Cells and Tissues. *ACS Synth Biol*. 2**017** 6(1):13-18. doi: 10.1021/acssynbio.6b00092.
13. Chatterjee A, Xiao H, Bollong M, Ai HW, Schultz PG. Efficient viral delivery system for unnatural amino acid mutagenesis in mammalian cells. *Proc Natl Acad Sci U S A.* **2013** 110(29):11803-8. doi: 10.1073/pnas.1309584110
14. Kim SW, Li Z, Moore PS, Monaghan AP, Chang Y, Nichols M, John B. A sensitive non-radioactive northern blot method to detect small RNAs. *Nucleic Acids Res.* **2010** 38(7):e98. doi: 10.1093/nar/gkp1235

**Plasmid maps and sequences**

**Color scheme: CMV MTH aaRS EGFP mCherry PAD4 tRNA Promoter tRNA** ITR

*pAAV-ITR-CMV-mCherry-1xU6-LtR-wt(TAG)*

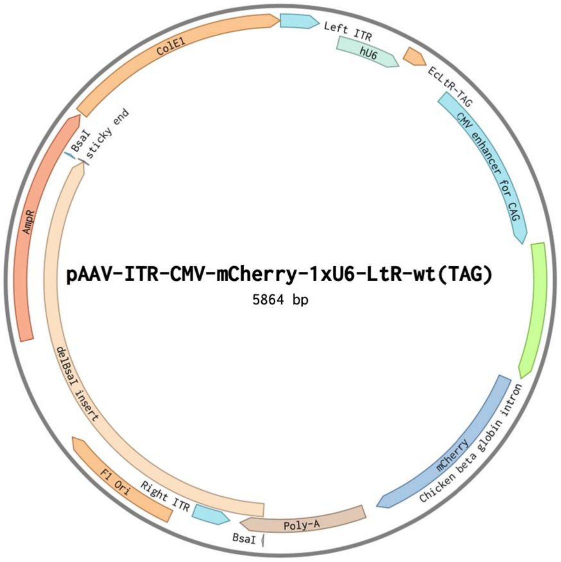


CCTGCAGGCAGCTGCGCGCTCGCTCGCTCACTGAGGCCGCCCGGGCAAAGCCCGGGCGTCGGGCGACC
TTTGGTCGCCCGGCCTCAGTGAGCGAGCGAGCGCGCAGAGAGGGAGTGGCCAACTCCATCACTAGGG
GTTCCTGCGGCCGCACGCGTCTCGCGGTCCAGTAGTGATCGACACTGCTCGATCCGCTCGCACCCCTAG
GGGTACCTCGGGCAGGAA**GAGGGCCTATTTCCCATGATTCCTTCATATTTGCATATACGATACAAG**
**GCTGTTAGAGAGATAATTAGAATTAATTTGACTGTAAACACAAAGATATTAGTACAAAATACGTG**
**ACGTAGAAAGTAATAATTTCTTGGGTAGTTTGCAGTTTTAAAATTATGTTTTAAAATGGACTATC**
**ATATGCTTACCGTAACTTGAAAGTATTTCGATTTCTTGGCTTTATATATCTTGTGGAAAGGACGA**
**AACACCGCCCGGATGGTGGAATCGGTAGACACAAGGGATTCTAAATCCCTCGGCGTTCGCGCTG**
**TGCGGGTTCAAGTCCCGCTCCGGGTA**TTTTTTGCTAGCGGATCGACGAGAGCAGCGCGACTGGATC
TGTCGCCCGTCTCAAACGCAACCCTCCGGCGGTCGCATATCATTCAGGACGAGCCTCAGACTCCAGCG
TAACACGCGT**CTAGTTATTAATAGTAATCAATTACGGGGTCATTAGTTCATAGCCCATATATGGA**
**GTTCCGCGTTACATAACTTACGGTAAATGGCCCGCCTGGCTGACCGCCCAACGACCCCCGCCCA**
**TTGACGTCAATAATGACGTATGTTCCCATAGTAACGTCAATAGGGACTTTCCATTGACGTCAATG**
**GGTGGAGTATTTACGGTAAACTGCCCACTTGGCAGTACATCAAGTGTATCATATGCCAAGTACG**
**CCCCCTATTGACGTCAATGACGGTAAATGGCCCGCCTGGCATTATGCCCAGTACATGACCTTAT**
**GGGACTTTCCTACTTGGCAGTACATCTACGTATTAGTCATCGCTATTACCATGGTGATGCGGTTT**
**TGGCAGTACATCAATGGGCGTGGATAGCGGTTTGACTCACGGGGATTTCCAAGTCTCCACCCCA**
**TTGACGTCAATGGGAGTTTGTTTTGCACCAAAATCAACGGGACTTTCCAAAATGTCGTAACAACT**
**CCGCCCCATTGACGCAAATGGGCGGTAGGCGTGTACGGTGGGAGGTCTATATAAGCAGAGCTC**
**GTTTAGTGAACCGTCAGATCGCCTGGAGACGCCATCCACGCTGTTTTGACCTCCATAGAAGACA**
**CCGGGACCGATCCAGCCTCCGCGGATTCGAATCCCGGCCGGGAACGGTGCATTGGAACGCGGA**
**TTCCCCGTGCCAAGAGTGACGTAAGTACCGCCTATAGAGTCTATAGGCCCACAAAAAATGCTTT**
**CTTCTTTTAATATACTTTTTTGTTTATCTTATTTCTAATACTTTCCCTAATCTCTTTCTTTCAGGGC**
**AATAATGATACAATGTATCATGCCTCTTTGCACCATTCTAAAGAATAACAGTGATAATTTCTGGG**
**TTAAGGCAATAGCAATATTTCTGCATATAAATATTTCTGCATATAAATTGTAACTGATGTAAGAG**
**GTTTCATATTGCTAATAGCAGCTACAATCCAGCTACCATTCTGCTTTTATTTTATGGTTGGGATA**
**AGGCTGGATTATTCTGAGTCCAAGCTAGGCCCTTTTGCTAATCATGTTCATACCTCTTATCTTCC**
**TCCCACAGCTCCTGGGCAACGTGCTGGTCTGTGTGCTGGCCCATCACTTTGGCAAAGAATTGGG**
**AT**TCGAACATCGATTGAATTC**ATGGTGAGCAAGGGCGAGGAGGATAACATGGCCATCATCAAGGA**
**GTTCATGCGCTTCAAGGTGCACATGGAGGGCTCCGTGAACGGCCACGAGTTCGAGATCGAGGG**
**CGAGGGCGAGGGCCGCCCCTACGAGGGCACCCAGACCGCCAAGCTGAAGGTGACCAAGGGTGG**

**CCCCCTGCCCTTCGCCTGGGACATCCTGTCCCCTCAGTTCATGTACGGCTCCAAGGCCTACGTG**
**AAGCACCCCGCCGACATCCCCGACTACTTGAAGCTGTCCTTCCCCGAGGGCTTCAAGTGGGAGC**
**GCGTGATGAACTTCGAGGACGGCGGCGTGGTGACCGTGACCCAGGACTCCTCCCTGCAGGACG**
**GCGAGTTCATCTACAAGGTGAAGCTGCGCGGCACCAACTTCCCCTCCGACGGCCCCGTAATGCA**
**GAAGAAGACAATGGGCTGGGAGGCCTCCTCCGAGCGGATGTACCCCGAGGACGGCGCCCTGAA**
**GGGCGAGATCAAGCAGAGGCTGAAGCTGAAGGACGGCGGCCACTACGACGCTGAGGTCAAGAC**
**CACCTACAAGGCCAAGAAGCCCGTGCAGCTGCCCGGCGCCTACAACGTCAACATCAAGTTGGAC**
**ATCACCTCCCACAACGAGGACTACACCATCGTGGAACAGTACGAACGCGCCGAGGGCCGCCACT**
**CCACCGGCGGCATGGACGAGCTGTACAAGTAA**TCTAGAGTCGACCTGCAGAAGCTTGCCTCGAGCA
GCGCTGCTCGAGAGATCTACGGGTGGCATCCCTGTGACCCCTCCCCAGTGCCTCTCCTGGCCCTGGAA
GTTGCCACTCCAGTGCCCACCAGCCTTGTCCTAATAAAATTAAGTTGCATCATTTTGTCTGACTAGGTG
TCCTTCTATAATATTATGGGGTGGAGGGGGGTGGTATGGAGCAAGGGGCAAGTTGGGAAGACAACCT
GTAGGGCCTGCGGGGTCTATTGGGAACCAAGCTGGAGTGCAGTGGCACAATCTTGGCTCACTGCAATC
TCCGCCTCCTGGGTTCAAGCGATTCTCCTGCCTCAGCCTCCCGAGTTGTTGGGATTCCAGGCATGCATG
ACCAGGCTCAGCTAATTTTTGTTTTTTTGGTAGAGACGGGGTTTCACCATATTGGCCAGGCTGGTCTCC
AACTCCTAATCTCAGGTGATCTACCCACCTTGGCCTCCCAAATTGCTGGGATTACAGGCGTGAACCACT
GCTCCCTTCCCTGTCCTTCTGATTTTGTAGGTAACCACGTGCGGACCGAGCGGCCGCAGGAACCCCTAG
TGATGGAGTTGGCCACTCCCTCTCTGCGCGCTCGCTCGCTCACTGAGGCCGGGCGACCAAAGGTCGCC
CGACGCCCGGGCTTTGCCCGGGCGGCCTCAGTGAGCGAGCGAGCGCGCAGCTGCCTGCAGGGGCGCC
TGATGCGGTATTTTCTCCTTACGCATCTGTGCGGTATTTCACACCGCATACGTCAAAGCAACCATAGTA
CGCGCCCTGTAGCGGCGCATTAAGCGCGGCGGGTGTGGTGGTTACGCGCAGCGTGACCGCTACACTTG
CCAGCGCCCTAGCGCCCGCTCCTTTCGCTTTCTTCCCTTCCTTTCTCGCCACGTTCGCCGGCTTTCCCCG
TCAAGCTCTAAATCGGGGGCTCCCTTTAGGGTTCCGATTTAGTGCTTTACGGCACCTCGACCCCAAAAA
ACTTGATTTGGGTGATGGTTCACGTAGTGGGCCATCGCCCTGATAGACGGTTTTTCGCCCTTTGACGTT
GGAGTCCACGTTCTTTAATAGTGGACTCTTGTTCCAAACTGGAACAACACTCAACCCTATCTCGGGCTA
TTCTTTTGATTTATAAGGGATTTTGCCGATTTCGGCCTATTGGTTAAAAAATGAGCTGATTTAACAAAA
ATTTAACGCGAATTTTAACAAAATATTAACGTTTACAATTTTATGGTGCACTCTCAGTACAATCTGCTC
TGATGCCGCATAGTTAAGCCAGCCCCGACACCCGCCAACACCCGCTGACGCGCCCTGACGGGCTTGTC
TGCTCCCGGCATCCGCTTACAGACAAGCTGTGACCGTCTCCGGGAGCTGCATGTGTCAGAGGTTTTCAC
CGTCATCACCGAAACGCGCGAGACGAAAGGGCCTCGTGATACGCCTATTTTTATAGGTTAATGTCATG
ATAATAATGGTTTCTTAGACGTCAGGTGGCACTTTTCGGGGAAATGTGCGCGGAACCCCTATTTGTTTA
TTTTTCTAAATACATTCAAATATGTATCCGCTCATGAGACAATAACCCTGATAAATGCTTCAATAATAT
TGAAAAAGGAAGAGTATGAGTATTCAACATTTCCGTGTCGCCCTTATTCCCTTTTTTGCGGCATTTTGC
CTTCCTGTTTTTGCTCACCCAGAAACGCTGGTGAAAGTAAAAGATGCTGAAGATCAGTTGGGTGCACG
AGTGGGTTACATCGAACTGGATCTCAACAGCGGTAAGATCCTTGAGAGTTTTCGCCCCGAAGAACGTT
TTCCAATGATGAGCACTTTTAAAGTTCTGCTATGTGGCGCGGTATTATCCCGTATTGACGCCGGGCAAG
AGCAACTCGGTCGCCGCATACACTATTCTCAGAATGACTTGGTTGAGTACTCACCAGTCACAGAAAAG
CATCTTACGGATGGCATGACAGTAAGAGAATTATGCAGTGCTGCCATAACCATGAGTGATAACACTGC
GGCCAACTTACTTCTGACAACGATCGGAGGACCGAAGGAGCTAACCGCTTTTTTGCACAACATGGGGG
ATCATGTAACTCGCCTTGATCGTTGGGAACCGGAGCTGAATGAAGCCATACCAAACGACGAGCGTGAC
ACCACGATGCCTGTAGCAATGGCAACAACGTTGCGCAAACTATTAACTGGCGAACTACTTACTCTAGC
TTCCCGGCAACAATTAATAGACTGGATGGAGGCGGATAAAGTTGCAGGACCACTTCTGCGCTCGGCCC
TTCCGGCTGGCTGGTTTATTGCTGATAAATCTGGAGCCGGTGAGCGTGGGTCTCGCGGTATCATTGCAG
CACTGGGGCCAGATGGTAAGCCCTCCCGTATCGTAGTTATCTACACGACGGGGAGTCAGGCAACTATG
GATGAACGAAATAGACAGATCGCTGAGATAGGTGCCTCACTGATTAAGCATTGGTAACTGTCAGACCA
AGTTTACTCATATATACTTTAGATTGATTTAAAACTTCATTTTTAATTTAAAAGGATCTAGGTGAAGAT
CCTTTTTGATAATCTCATGACCAAAATCCCTTAACGTGAGTTTTCGTTCCACTGAGCGTCAGACCCCGT
AGAAAAGATCAAAGGATCTTCTTGAGATCCTTTTTTTCTGCGCGTAATCTGCTGCTTGCAAACAAAAAA
ACCACCGCTACCAGCGGTGGTTTGTTTGCCGGATCAAGAGCTACCAACTCTTTTTCCGAAGGTAACTGG
CTTCAGCAGAGCGCAGATACCAAATACTGTCCTTCTAGTGTAGCCGTAGTTAGGCCACCACTTCAAGA
ACTCTGTAGCACCGCCTACATACCTCGCTCTGCTAATCCTGTTACCAGTGGCTGCTGCCAGTGGCGATA
AGTCGTGTCTTACCGGGTTGGACTCAAGACGATAGTTACCGGATAAGGCGCAGCGGTCGGGCTGAACG
GGGGGTTCGTGCACACAGCCCAGCTTGGAGCGAACGACCTACACCGAACTGAGATACCTACAGCGTG
AGCTATGAGAAAGCGCCACGCTTCCCGAAGGGAGAAAGGCGGACAGGTATCCGGTAAGCGGCAGGGT
CGGAACAGGAGAGCGCACGAGGGAGCTTCCAGGGGGAAACGCCTGGTATCTTTATAGTCCTGTCGGG

TTTCGCCACCTCTGACTTGAGCGTCGATTTTTGTGATGCTCGTCAGGGGGGCGGAGCCTATGGAAAA
CGCCAGCAACGCGGCCTTTTTACGGTTCCTGGCCTTTTGCTGGCCTTTTGCTCACATGT

*pAcBac1-CMV-EGFP(39TAG)*

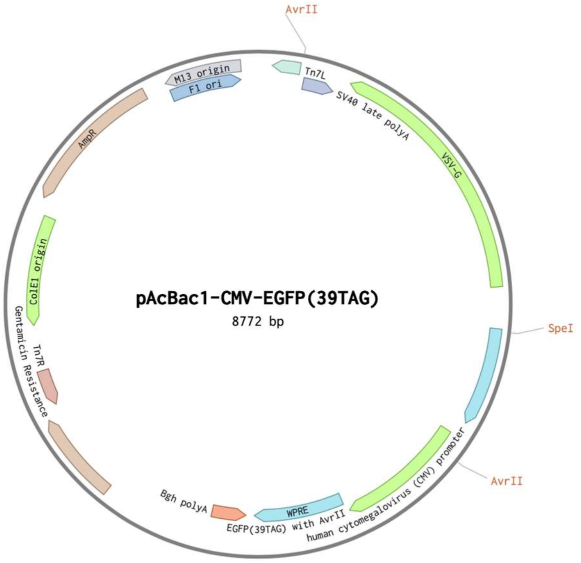


AAAGTAGCCGAAGATGACGGTTTGTCACATGGAGTTGGCAGGATGTTTGATTAAAAACATAACAGGA
AGAAAAATGCCCCGCTGTGGGCGGACAAAATAGTTGGGAACTGGGAGGGGTGGAAATGGAGTTTTTA
AGGATTATTTAGGGAAGAGTGACAAAATAGATGGGAACTGGGTGTAGCGTCGTAAGCTAATACGAAA
ATTAAAAATGACAAAATAGTTTGGAACTAGATTTCACTTATCTGGTTCGGATCTCCTAGGCTCAAGCA
GTGATCAGATCCAGACATGATAAGATACATTGATGAGTTTGGACAAACCACAACTAGAATGCAGTGA
AAAAAATGCTTTATTTGTGAAATTTGTGATGCTATTGCTTTATTTGTAACCATTATAAGCTGCAATAAA
CAAGTTAACAACAACAATTGCATTCATTTTATGTTTCAGGTTCAGGGGGAGGTGTGGGAGGTTTTTTAA
AGCAAGTAAAACCTCTACAAATGTGGTATGGCTGATTATGATCCTCTAGTACTTCTCGACAAGCTTGGC
CGGCCTGGCCTTATCACTTTCCAAGTCGGTTCATCTCTATGTCTGTATAAATCTGTCTTTTCTTGGTGTG
CTTTAATTTAATGCAAAGATGGATACCAACTCGGAGAACCAAGAATAGTCCAATGATTAACCCTATGA
TAAAGAAAAAAGAGGCAATAGAGCTTTTCCAACTACTGAACCAACCTTCTACAAGCTCGATTGGATTT
TTGGATAGCCCAGTATCACCAAAAAATAAACTCTCATCATCAGGAAGTTGCGAAGCAGCGTCTTGAAT
GTGAGGATGTTCGAACACCTGAGCCTTTGAGCTAAGATGAAGATCGGAGTCCAACATACCATGTCCAA
TCATGTATAAAGGAAACTTATATCCTGAACTGGTCCTCAGAACTCCATTGGGTCCAATTTCCACGTCTT
CATATGGTGCCCAGTCATCCCACAGTTCCCTTTCTGTGGTAGTTCCACTGATCATTCCGACCATTCTTGA
GAGGATTGGAGCAGCAATATCGACTCTGATGTATCTGGTCTCAAAGTATTTTAGGGTACCATTGATTAT
GGTGAAAGCAGGACCGGTTCCTGGGTTTTTAGGAGCAAGATAGCTGAGATCCACTGGAGAGATTGGA
AGACCCGCTCTGATTTTGCTCCAGGTTTCTTGGCAGAGGGAATAATCCAAGATCCTCTCAACGTCCTGA
ATTAGACTTACATCCACTGAGGTCTGAGATGGAGCAGAGATACTTGACCCTTCTGGGCATTCAGGGAA
TCTGGCTGCAGCAAAGAGATCCTTATCAGCCATCTCGAACCAGACACCTGATGGGAGTCTGACTCCCC
AATGCTTGCAGTATTGCATTTTGCAGGCCTTGCCTCCAGTTTCATAAGCAAAGTAGTTACTTCTGAACC
CTGTGCCCTCCTTTCCCAGGGATGATAGCTCTCCGTCCTCTGAGAAGAAGGTGATGTCCATGGAAATG
AGGTTAGAATCACATAGCCCTTTGACCTTATAGTCAGAATGCCAGGTTGTAGAGTTATGGACAGTGGG
GCATATGTAATTGCTGCATTTTCCGTTGATGAACTGTGAATCAACCCATTCTCCTGTGTATTCATCAAC
CAGCACATGGTGAGGAGTCACCTGGACAATCACTGCTTCGGCATCCGTCACAGTTGCATATCCACAAC
TTTGAGGAGGGAAGCCTGGATTCAGCCAAGTTCCTTGTTTCGTTTGTTCAATGCTTTCCTTGCATTGTTC
TACAGATGGAGTGAAGGATCGGATGGAATGTGTTATATACTTCGGTCCATACCAGCGGAAATCACAAG
TAGTGACCCATTTGGAAGCATGACACATCCAACCGTCTGCTTGAATAGCCTTGTGACTCTTGGGCATTT
TGACTTGTAAGGCTGTGCCTATTAAGTCATTATGCCAATTTAAATCTGAGCTTGACGGGCAATAATGGT
AATTAGAAGGAACATTTTTCCAGTTTCCTTTTTGGTTGTGTGGAAAAACTATGGTGAACTTGCAATTCA

CCCCAATGAATAAAAAGGCTAAGTACAAAAGGCACTTCATGGTGGCGGTTTTTAGGAGTTCGCGCCC
GATGGTGGGACGGTATGAATAATCCGGAATATTTATAGGTTTTTTTATTACAAAACTGTTACGAAAAC
AGTAAAATACTTATTTATTTGCGAGATGGTTATCATTTTAATTATCTCCATGATCTATTAATATTCCGGA
GTATACGGACCGCGGCCGCAAATACCTGCAGGATCCGTTTTGCGCTGCTTCGCGATGTACGGGCCAGA
TATACGCGTT**GACATTGATTATTGACTAGTTATTAATAGTAATCAATTACGGGGTCATTAGTTCAT**
**AGCCCATATATGGAGTTCCGCGTTACATAACTTACGGTAAATGGCCCGCCTGGCTGACCGCCCA**
**ACGACCCCCGCCCATTGACGTCAATAATGACGTATGTTCCCATAGTAACGCCAATAGGGACTTTC**
**CATTGACGTCAATGGGTGGACTATTTACGGTAAACTGCCCACTTGGCAGTACATCAAGTGTATCA**
**TATGCCAAGTACGCCCCCTATTGACGTCAATGACGGTAAATGGCCCGCCTGGCATTATGCCCAG**
**TACATGACCTTATGGGACTTTCCTACTTGGCAGTACATCTACGTATTAGTCATCGCTATTACCAT**
**GGTGATGCGGTTTTGGCAGTACATCAATGGGCGTGGATAGCGGTTTGACTCACGGGGATTTCCA**
**AGTCTCCACCCCATTGACGTCAATGGGAGTTTGTTTTGGCACCAAAATCAACGGGACTTTCCAAA**
**ATGTCGTAACAACTCCGCCCCATTGACGCAAATGGGCGGTAGGCGTGTACGGTGGGAGGTCTAT**
**ATAAGCAGAGCT**CTCTGGCTAACTAGAGAACCCACTGCTTACTGGCTTATCGAAATTAATACGACTCA
CTATAGGGAGACCCAAGCTGGCTAGCGCCGCCACC**ATGGTGAGCAAGGGCGAGGAGCTGTTCACC**
**GGGGTGGTGCCCATCCTGGTCGAGCTGGACGGCGACGTAAACGGCCACAAGTTCAGCGTGTCC**
**GGCGAGGGCGAGGGCGATGCCACCTAGGGCAAGCTGACCCTGAAGTTCATCTGCACCACCGGC**
**AAGCTGCCCGTGCCCTGGCCCACCCTCGTGACCACCCTGACCTACGGCGTGCAGTGCTTCAGCC**
**GCTACCCCGACCACATGAAGCAGCACGACTTCTTCAAGTCCGCCATGCCCGAAGGCTACGTCCA**
**GGAGCGCACCATCTTCTTCAAGGACGACGGCAACTACAAGACCCGCGCCGAGGTGAAGTTCGAG**
**GGCGACACCCTGGTGAACCGCATCGAGCTGAAGGGCATCGACTTCAAGGAGGACGGCAACATC**
**CTGGGGCACAAGCTGGAGTACAACTACAACAGCCACAACGTCTATATCATGGCCGACAAGCAGA**
**AGAACGGCATCAAGGTGAACTTCAAGATCCGCCACAACATCGAGGACGGCAGCGTGCAGCTCGC**
**CGACCACTACCAGCAGAACACCCCCATCGGCGACGGCCCCGTGCTGCTGCCCGACAACCACTAC**
**CTGAGCACCCAGTCCGCCCTGAGCAAAGACCCCAACGAGAAGCGCGATCACATGGTCCTGCTGG**
**AGTTCGTGACCGCCGCCGGGATCACTCTCGGCATGGACGAGCTGTACAAGGGGCCCTTCGAACA**
**AAAACTCATCTCAGAAGAGGATCTGAATATGCATACCGGTCATCATCACCATCACCATTGA**ATTC
AACGCGTTAAGTCGACAATCAACCTCTGGATTACAAAATTTGTGAAAGATTGACTGGTATTCTTAACT
ATGTTGCTCCTTTTACGCTATGTGGATACGCTGCTTTAATGCCTTTGTATCATGCTATTGCTTCCCGTAT
GGCTTTCATTTTCTCCTCCTTGTATAAATCCTGGTTGCTGTCTCTTTATGAGGAGTTGTGGCCCGTTGTC
AGGCAACGTGGCGTGGTGTGCACTGTGTTTGCTGACGCAACCCCCACTGGTTGGGGCATTGCCACCAC
CTGTCAGCTCCTTTCCGGGACTTTCGCTTTCCCCCTCCCTATTGCCACGGCGGAACTCATCGCCGCCTGC
CTTGCCCGCTGCTGGACAGGGGCTCGGCTGTTGGGCACTGACAATTCCGTGGTGTTGTCGGGGAAATC
ATCGTCCTTTCCTTGGCTGCTCGCCTGTGTTGCCACCTGGATTCTGCGCGGGACGTCCTTCTGCTACGTC
CCTTCGGCCCTCAATCCAGCGGACCTTCCTTCCCGCGGCCTGCTGCCGGCTCTGCGGCCTCTTCCGCGT
CTTCGCCTTCGCCCTCAGACGAGTCGGATCTCCCTTTGGGCCGCCTCCCCGCGTCGACTTTAACTCGAG
TCTAGAGGGCCCGTTTAAACCCGCTGATCAGCCTCGACTGTGCCTTCTAGTTGCCAGCCATCTGTTGTT
TGCCCCTCCCCCGTGCCTTCCTTGACCCTGGAAGGTGCCACTCCCACTGTCCTTTCCTAATAAAATGAG
GAAATTGCATCGCATTGTCTGAGTAGGTGTCATTCTATTCTGGGGGGTGGGGTGGGGCAGGACAGCAA
GGGGGAGGATTGGGAAGACAATAGCAGGCATGCTGGGGATGCGGTGGGCTCTATGGCTTCTGAGGCG
GAAAGAACCAGCTGGGGCTCTAGGGGGTATCCCCACGCGCCCTGTAGCGGCGCATTAAGCGCAAGCC
CTGCCATAGCCACTACGGGTACGTAGCCAACCACTAGAACTATAGCTAGAGTCCTGGGCGAACAAACG
ATGCTCGCCTTCCAGAAAACCGAGGATGCGAACCACTTCATCCGGGGTCAGCACCACCGGCAAGCGCC
GCGACGGCCGAGGTCTACCGATCTCCTGAAGCCAGGGCAGATCCGTGCACAGCACCTTGCCGTAGAAG
AACAGCAAGGCCGCCAATGCCTGACGATGCGTGGAGACCGAAACCTTGCGCTCGTTCGCCAGCCAGG
ACAGAAATGCCTCGACTTCGCTGCTGCCCAAGGTTGCCGGGTGACGCACACCGTGGAAACGGATGAA
GGCACGAACCCAGTTGACATAAGCCTGTTCGGTTCGTAAACTGTAATGCAAGTAGCGTATGCGCTCAC
GCAACTGGTCCAGAACCTTGACCGAACGCAGCGGTGGTAACGGCGCAGTGGCGGTTTTCATGGCTTGT
TATGACTGTTTTTTTGTACAGTCTATGCCTCGGGCATCCAAGCAGCAAGCGCGTTACGCCGTGGGTCGA
TGTTTGATGTTATGGAGCAGCAACGATGTTACGCAGCAGCAACGATGTTACGCAGCAGGGCAGTCGCC
CTAAAACAAAGTTAGGTGGCTCAAGTATGGGCATCATTCGCACATGTAGGCTCGGCCCTGACCAAGTC
AAATCCATGCGGGCTGCTCTTGATCTTTTCGGTCGTGAGTTCGGAGACGTAGCCACCTACTCCCAACAT
CAGCCGGACTCCGATTACCTCGGGAACTTGCTCCGTAGTAAGACATTCATCGCGCTTGCTGCCTTCGAC
CAAGAAGCGGTTGTTGGCGCTCTCGCGGCTTACGTTCTGCCCAGGTTTGAGCAGCCGCGTAGTGAGAT
CTATATCTATGATCTCGCAGTCTCCGGCGAGCACCGGAGGCAGGGCATTGCCACCGCGCTCATCAATC
TCCTCAAGCATGAGGCCAACGCGCTTGGTGCTTATGTGATCTACGTGCAAGCAGATTACGGTGACGAT

CCCGCAGTGGCTCTCTATACAAAGTTGGGCATACGGGAAGAAGTGATGCACTTTGATATCGACCCAAG
TACCGCCACCTAACAATTCGTTCAAGCCGAGATCGGCTTCCCGGCCGCGGAGTTGTTCGGTAAATTGTC
ACAACGCCGCGAATATAGTCTTTACCATGCCCTTGGCCACGCCCCTCTTTAATACGACGGGCAATTTGC
ACTTCAGAAAATGAAGAGTTTGCTTTAGCCATAACAAAAGTCCAGTATGCTTTTTCACAGCATAACTG
GACTGATTTCAGTTTACAACTATTCTGTCTAGTTTAAGACTTTATTGTCATAGTTTAGATCTATTTTGTT
CAGTTTAAGACTTTATTGTCCGCCCACACCCGCTTACGCAGGGCATCCATTTATTACTCAACCGTAACC
GATTTTGCCAGGTTACGCGGCTGGTCTGCGGTGTGAAATACCGCACAGATGCGTAAGGAGAAAATACC
GCATCAGGCGCTCTTCCGCTTCCTCGCTCACTGACTCGCTGCGCTCGGTCGTTCGGCTGCGGCGAGCGG
TATCAGCTCACTCAAAGGCGGTAATACGGTTATCCACAGAATCAGGGGATAACGCAGGAAAGAACAT
GTGAGCAAAAGGCCAGCAAAAGGCCAGGAACCGTAAAAAGGCCGCGTTGCTGGCGTTTTTCCATAGG
CTCCGCCCCCCTGACGAGCATCACAAAAATCGACGCTCAAGTCAGAGGTGGCGAAACCCGACAGGAC
TATAAAGATACCAGGCGTTTCCCCCTGGAAGCTCCCTCGTGCGCTCTCCTGTTCCGACCCTGCCGCTTA
CCGGATACCTGTCCGCCTTTCTCCCTTCGGGAAGCGTGGCGCTTTCTCAATGCTCACGCTGTAGGTATC
TCAGTTCGGTGTAGGTCGTTCGCTCCAAGCTGGGCTGTGTGCACGAACCCCCCGTTCAGCCCGACCGCT
GCGCCTTATCCGGTAACTATCGTCTTGAGTCCAACCCGGTAAGACACGACTTATCGCCACTGGCAGCA
GCCACTGGTAACAGGATTAGCAGAGCGAGGTATGTAGGCGGTGCTACAGAGTTCTTGAAGTGGTGGCC
TAACTACGGCTACACTAGAAGGACAGTATTTGGTATCTGCGCTCTGCTGAAGCCAGTTACCTTCGGAA
AAAGAGTTGGTAGCTCTTGATCCGGCAAACAAACCACCGCTGGTAGCGGTGGTTTTTTTGTTTGCAAG
CAGCAGATTACGCGCAGAAAAAAAGGATCTCAAGAAGATCCTTTGATCTTTTCTACGGGGTCTGACGC
TCAGTGGAACGAAAACTCACGTTAAGGGATTTTGGTCATGAGATTATCAAAAAGGATCTTCACCTAGA
TCCTTTTAAATTAAAAATGAAGTTTTAAATCAATCTAAAGTATATATGAGTAAACTTGGTCTGACAGTT
ACCAATGCTTAATCAGTGAGGCACCTATCTCAGCGATCTGTCTATTTCGTTCATCCATAGTTGCCTGAC
TCCCCGTCGTGTAGATAACTACGATACGGGAGGGCTTACCATCTGGCCCCAGTGCTGCAATGATACCG
CGAGACCCACGCTCACCGGCTCCAGATTTATCAGCAATAAACCAGCCAGCCGGAAGGGCCGAGCGCA
GAAGTGGTCCTGCAACTTTATCCGCCTCCATCCAGTCTATTAATTGTTGCCGGGAAGCTAGAGTAAGTA
GTTCGCCAGTTAATAGTTTGCGCAACGTTGTTGCCATTGCTACAGGCATCGTGGTGTCACGCTCGTCGT
TTGGTATGGCTTCATTCAGCTCCGGTTCCCAACGATCAAGGCGAGTTACATGATCCCCCATGTTGTGCA
AAAAAGCGGTTAGCTCCTTCGGTCCTCCGATCGTTGTCAGAAGTAAGTTGGCCGCAGTGTTATCACTCA
TGGTTATGGCAGCACTGCATAATTCTCTTACTGTCATGCCATCCGTAAGATGCTTTTCTGTGACTGGTG
AGTACTCAACCAAGTCATTCTGAGAATAGTGTATGCGGCGACCGAGTTGCTCTTGCCCGGCGTCAATA
CGGGATAATACCGCGCCACATAGCAGAACTTTAAAAGTGCTCATCATTGGAAAACGTTCTTCGGGGCG
AAAACTCTCAAGGATCTTACCGCTGTTGAGATCCAGTTCGATGTAACCCACTCGTGCACCCAACTGATC
TTCAGCATCTTTTACTTTCACCAGCGTTTCTGGGTGAGCAAAAACAGGAAGGCAAAATGCCGCAAAAA
AGGGAATAAGGGCGACACGGAAATGTTGAATACTCATACTCTTCCTTTTTCAATATTATTGAAGCATTT
ATCAGGGTTATTGTCTCATGAGCGGATACATATTTGAATGTATTTAGAAAAATAAACAAATAGGGGTT
CCGCGCACATTTCCCCGAAAAGTGCCACCTGAAATTGTAAACGTTAATATTTTGTTAAAATTCGCGTTA
AATTTTTGTTAAATCAGCTCATTTTTTAACCAATAGGCCGAAATCGGCAAAATCCCTTATAAATCAAAA
GAATAGACCGAGATAGGGTTGAGTGTTGTTCCAGTTTGGAACAAGAGTCCACTATTAAAGAACGTGGA
CTCCAACGTCAAAGGGCGAAAAACCGTCTATCAGGGCGATGGCCCACTACGTGAACCATCACCCTAAT
CAAGTTTTTTGGGGTCGAGGTGCCGTAAAGCACTAAATCGGAACCCTAAAGGGAGCCCCCGATTTAGA
GCTTGACGGGGAAAGCCGGCGAACGTGGCGAGAAAGGAAGGGAAGAAAGCGAAAGGAGCGGGCGCT
AGGGCGCTGGCAAGTGTAGCGGTCACGCTGCGCGTAACCACCACACCCGCCGCGCTTAATGCGCCGCT
ACAGGGCGCGTCCCATTCGCCATTCAGGCTGCAAATAAGCGTTGATATTCAGTCAATTACAAACATTA
ATAACGAAGAGATGACAGAAAAATTTTCATTCTGTGACAGAGAA

*pIDTSMART-CMV-DanRS*

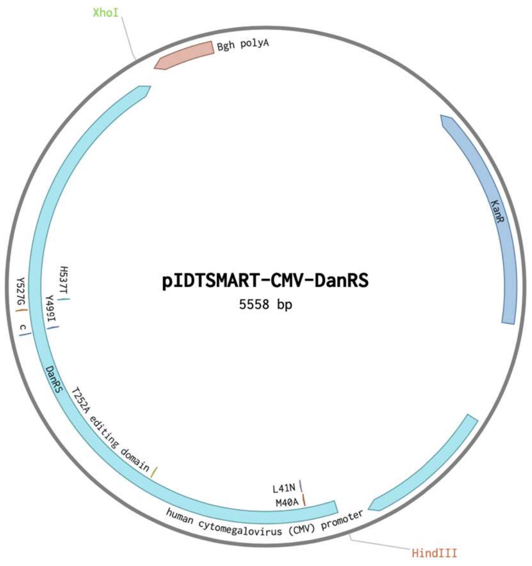


ACTGGCCGTCGTTTTACACGGGTGGGCCTTTCTTCGGTAGAAAATCAAAGGATCTTCTTGAGATCCTTT
TTTTCTGCGCGTAATCTGCTGCTTGCAAACAAAAAAACCACCGCTACCAGCGGTGGTTTGTTTGCCGGA
TCAAGAGCTACCAACTCTTTTTCCGAGGTAACTGGCTTCAGCAGAGCGCAGATACCAAATACTGTTCTT
CTAGTGTAGCCGTAGTTAGGCCACCACTTCAAGAACTCTGTAGCACCGCCTACATACCTCGCTCTGCTA
ATCCTGTTACCAGTGGCTGCTGCCAGTGGCGATAAGTCGTGTCTTACCGGGTTGGACTCAAGACGATA
GTTACCGGATAAGGCGCAGCGGTCGGGCTGAACGGGGGGTTCGTGCACACAGCCCAGCTTGGAGCGA
ACGACCTACACCGAACTGAGATACCTACAGCGTGAGCTATGAGAAAGCGCCACGCTTCCCGAAGGGA
GAAAGGCGGACAGGTATCCGGTAAGCGGCAGGGTCGGAACAGGAGAGCGCACGAGGGAGCTTCCAG
GGGGAAACGCCTGGTATCTTTATAGTCCTGTCGGGTTTCGCCACCTCTGACTTGAGCATCGATTTTTGT
GATGCTCGTCAGGGGGGCGGAGCCTATGGAAAAACGCCAGCAACGCAGAAAGGCCCACCCGAAGGTG
AGCCAGGTGATTACATTTAGGTCCTCATTAGAAAAACTCATCGAGCATCAAGTGAAACTGCAATTTAT
TCATATCAGGATTATCAATACCATATTTTTGAAAAAGCCGTTTCTGTAATGAAGGAGAAAACTCACCG
AGGCAGTTCCATAGGATGGCAAGATCCTGGTATCGGTCTGCGATTCCGACTCGTCCAACATCAATACA
ACCTATTAATTTCCCCTCGTCAAAAATAAGGTTATCAAGTGAGAAATCACCATGAGTGACGACTGAAT
CCGGTGAGAATGGCAAGAGTTTATGCATTTCTTTCCAGACTTGTTCAACAGGCCAGCCATTACGCTCGT
CATCAAAATCACTCGCACCAACCAAACCGTTATTCATTCGTGATTGCGCCTGAGCGAGACGAAATACG
CGATCGCCGTTAAAAGGACAATTACAAACAGGAATCGAATGCAACCGGCGCAGGAACACTGCCAGCG
CATCAACAATATTTTCACCTGAATCAGGATATTCTTCTAATACCTGGAATGCTGTTTTCCCTGGGATCG
CAGTGGTGAGTAACCATGCATCATCAGGAGTACGGATAAAATGCTTGATGGTCGGAAGAGGCATAAA
TTCCGTCAGCCAGTTTAGCCTGACCATCTCATCTGTAACATCATTGGCAACGCTACCTTTGCCATGTTTC
AGAAACAACTCTGGCGCATCGGGCTTCCCATACAATCGATAGATTGTCGCACCTGATTGCCCGACATT
ATCGCGAGCCCATTTATACCCATATAAATCAGCATCCATGTTGGAATTTAATCGCGGCTTCAAGCAAG
ACGTTTCCCGTTGAATATGGCTCATTTTAGCTTCCTTAGCTCCTGAAAATCTCGATAACTCAAAAAATA
CGCCCGGTAGTGATCTTATTTCATTATGGTGAAAGTTGGAACCTCTTACGTGCCGATCAAGTCAAAAGC
CTCCGGTCGGAGGCTTTTGACTTTCTGCTATGGAGGTCAGGTATGATTTAAATGGTCAGTATTGAGCCT
CAGGAAACAGCTATGACATCAAGCTGACTAGATAATCTAGCTGATCGTGGACCGATCATACGTATAAT
GCCGTAAGATCACGGGTCGCAGCACAGCTCGCGGTCCAGTAGTGATCGACACTGCTCGATCCGCTCGC
ACCCCTAGGTGCGCTGCTTCGCGATGTACGGGCCAGATATACGCGTT**GACATTGATTATTGACTAGT**
**TATTAATAGTAATCAATTACGGGGTCATTAGTTCATAGCCCATATATGGAGTTCCGCGTTACATA**
**ACTTACGGTAAATGGCCCGCCTGGCTGACCGCCCAACGACCCCCGCCCATTGACGTCAATAATG**
**ACGTATGTTCCCATAGTAACGCCAATAGGGACTTTCCATTGACGTCAATGGGTGGAGTATTTACG**
**GTAAACTGCCCACTTGGCAGTACATCAAGTGTATCATATGCCAAGTACGCCCCCTATTGACGTCA**
**ATGACGGTAAATGGCCCGCCTGGCATTATGCCCAGTACATGACCTTATGGGACTTTCCTACTTG**
**GCAGTACATCTACGTATTAGTCATCGCTATTACCATGGTGATGCGGTTTTGGCAGTACATCAATG**
**GGCGTGGATAGCGGTTTGACTCACGGGGATTTCCAAGTCTCCACCCCATTGACGCAAATGGGCG**
**GTAGGCGTGTACGGTGGGAGGTCTATATAAGCAGAGCT**CTCTGGCTAACTAGAGAACCCACTGCTT

ACTGGCTTATCGAAATTAATACGACTCACTATAGGGAGACCCAAGCTGGCTAGCGTTTAAACTTAAGC
TTGCCGCCACC**ATGGAAGAGCAATACCGCCCGGAAGAGATAGAATCCAAAGTACAGCTTCATTGG**
**GATGAGAAGCGCACATTTGAAGTAACCGAAGACGAGAGCAAAGAGAAGTATTACTGCCTGTCTG**
**CTAATCCCTATCCTTCTGGTCGACTACACATGGGCCACGTACGTAACTACACCATCGGTGACGTG**
**ATCGCCCGCTACCAGCGTATGCTGGGCAAAAACGTCCTGCAGCCGATCGGCTGGGACGCGTTTG**
**GTCTGCCTGCGGAAGGCGCGGCGGTGAAAAACAACACCGCTCCGGCACCGTGGACGTACGACA**
**ACATCGCGTATATGAAAAACCAGCTCAAAATGCTGGGCTTTGGTTATGACTGGAGCCGCGAGCT**
**GGCAACCTGTACGCCGGAATACTACCGTTGGGAACAGAAATTCTTCACCGAGCTGTATAAAAAA**
**GGCCTGGTATATAAGAAGACTTCTGCGGTCAACTGGTGTCCGAACGACCAGACCGTACTGGCGA**
**ACGAACAAGTTATCGACGGCTGCTGCTGGCGCTGCGATACCAAAGTTGAACGTAAAGAGATCCC**
**GCAGTGGTTTATCAAAATCACTGCTTACGCTGACGAGCTGCTCAACGATCTGGATAAACTGGAT**
**CACTGGCCAGACACCGTTAAAACCATGCAGCGTAACTGGATCGGTCGTTCCGAAGGCGTGGAGA**
**TCACCTTCAACGTTAACGACTATGACAACACGCTGACCGTTTACACTACCCGCCCGGACGCGTTT**
**ATGGGTTGTACCTACCTGGCGGTAGCTGCGGGTCATCCGCTGGCGCAGAAAGCGGCGGAAAAT**
**AATCCTGAACTGGCGGCCTTTATTGACGAATGCCGTAACACCAAAGTTGCCGAAGCTGAAATGG**
**CGACGATGGAGAAAAAAGGCGTCGATACTGGCTTTAAAGCGGTTCACCCATTAACGGGCGAAGA**
**AATTCCCGTTTGGGCAGCAAACTTCGTATTGATGGAGTACGGCACGGGCGCAGTTATGGCGGTA**
**CCGGGGCACGACCAGCGCGACTACGAGTTTGCCTCTAAATACGGCCTGAACATCAAACCGGTTA**
**TCCTGGCAGCTGACGGCTCTGAGCCAGATCTTTCTCAGCAAGCCCTGACTGAAAAAGGCGTGCT**
**GTTCAACTCTGGCGAGTTCAACGGTCTTGACCATGAAGCGGCCTTCAACGCCATCGCCGATAAA**
**CTGACTGCGATGGGCGTTGGCGAGCGTAAAGTGAACTACCGCCTGCGCGACTGGGGTGTTTCCC**
**GTCAGCGTTACTGGGGCGCGCCGATTCCGATGGTGACTCTAGAAGACGGTACCGTAATGCCGAC**
**CCCGGACGACCAGCTGCCGGTGATCCTGCCGGAAGATGTGGTAATGGACGGCATTACCAGCCC**
**GATTAAAGCAGATCCGGAGTGGGCGAAAACTACCGTTAACGGTATGCCAGCACTGCGTGAAACC**
**GACACTTTCGACACCTTTATGGAGTCCTCCTGGATTTATGCGCGCTACACTTGCCCGCAGTACAA**
**AGAAGGTATGCTGGATTCCGAAGCGGCTAACTACTGGCTGCCGGTGGATATCGGTATTGGTGGT**
**ATTGAACACGCCATTATGACGCTGCTCTACTTCCGCTTCTTCCACAAACTGATGCGTGATGCAGG**
**CATGGTGAACTCTGACGAACCAGCGAAACAGTTGCTGTGTCAGGGTATGGTGCTGGCAGATGCC**
**TTCTACTATGTTGGCGAAAACGGCGAACGTAACTGGGTTTCCCCGGTTGATGCTATCGTTGAAC**
**GTGACGAGAAAGGCCGTATCGTGAAAGCGAAAGATGCGGCAGGCCATGAACTGGTTTATACCG**
**GCATGAGCAAAATGTCCAAGTCGAAGAACAACGGTATCGACCCGCAGGTGATGGTTGAACGTTA**
**CGGCGCGGACACCGTTCGTCTGTTTATGATGTTTGCTTCTCCGGCTGATATGACTCTCGAATGGC**
**AGGAATCCGGTGTGGAAGGGGCTAACCGCTTCCTGAAACGTGTCTGGAAACTGGTTTACGAGCA**
**CACAGCAAAAGGTGATGTTGCGGCACTGAACGTTGATGCGCTGACTGAAAATCAGAAAGCGCTG**
**CGTCGCGATGTGCATAAAACGATCGCTAAAGTGACCGATGATATCGGCCGTCGTCAGACCTTCA**
**ACACCGCAATTGCGGCGATTATGGAGCTGATGAACAAACTGGCGAAAGCACCAACCGATGGCGA**
**GCAGGATCGCGCTCTGATGCAGGAAGCACTGCTGGCCGTTGTCCGTATGCTTAACCCGTTCACC**
**CCGCACATCTGCTTCACGCTGTGGCAGGAACTGAAAGGCGAAGGCGATATCGACAACGCGCCGT**
**GGCCGGTTGCTGACGAAAAAGCGATGGTGGAAGACTCCACGCTGGTCGTGGTGCAGGTTAACG**
**GTAAAGTCCGTGCCAAAATCACCGTTCCGGTGGACGCAACGGAAGAACAGGTTCGCGAACGTGC**
**TGGCCAGGAACATCTGGTAGCAAAATATCTTGATGGCGTTACTGTACGTAAAGTGATTTACGTAC**
**CAGGTAAACTCCTCAATCTGGTCGTTGGCGGGCCCGTTTAA**CTCGAGTCTAGAGGGCCCGTTTAAA
CCCGCTGATCAGCCTCGACTGTGCCTTCTAGTTGCCAGCCATCTGTTGTTTGCCCCTCCCCCGTGCCTTC
CTTGACCCTGGAAGGTGCCACTCCCACTGTCCTTTCCTAATAAAATGAGGAAATTGCATCGCATTGTCT
GAGTAGGTGTCATTCTATTCTGGGGGGTGGGGTGGGGCAGGACAGCAAGGGGGAGGATTGGGAAGAC
AATAGCAGGCATGCTGGGGATGCGGTGGGCTCTATGGCTTCTGAGGCGGAAAGAACCCTAGCGGATC
GACGAGAGCAGCGCGACTGGATCTGTCGCCCGTCTCAAACGCAACCCTCCGGCGGTCGCATATCATTC
AGGACGAGCCTCAGACTCCAGCGTAACTGGACTGCAATCAACTCACTGGCTCACCTTCCGGTCCACGA
TCAGCTAGAATCAAGCTGACTAGATAA

*pIDTSmart-CMV-PLRS1*

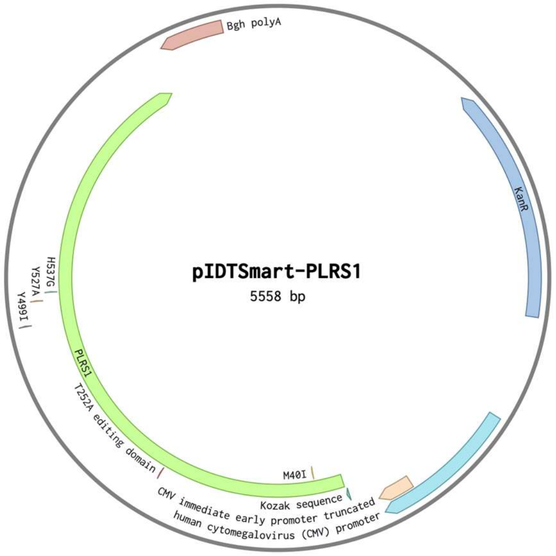


ACTGGCCGTCGTTTTACACGGGTGGGCCTTTCTTCGGTAGAAAATCAAAGGATCTTCTTGAGATCCTTT
TTTTCTGCGCGTAATCTGCTGCTTGCAAACAAAAAAACCACCGCTACCAGCGGTGGTTTGTTTGCCGGA
TCAAGAGCTACCAACTCTTTTTCCGAGGTAACTGGCTTCAGCAGAGCGCAGATACCAAATACTGTTCTT
CTAGTGTAGCCGTAGTTAGGCCACCACTTCAAGAACTCTGTAGCACCGCCTACATACCTCGCTCTGCTA
ATCCTGTTACCAGTGGCTGCTGCCAGTGGCGATAAGTCGTGTCTTACCGGGTTGGACTCAAGACGATA
GTTACCGGATAAGGCGCAGCGGTCGGGCTGAACGGGGGGTTCGTGCACACAGCCCAGCTTGGAGCGA
ACGACCTACACCGAACTGAGATACCTACAGCGTGAGCTATGAGAAAGCGCCACGCTTCCCGAAGGGA
GAAAGGCGGACAGGTATCCGGTAAGCGGCAGGGTCGGAACAGGAGAGCGCACGAGGGAGCTTCCAG
GGGGAAACGCCTGGTATCTTTATAGTCCTGTCGGGTTTCGCCACCTCTGACTTGAGCATCGATTTTTGT
GATGCTCGTCAGGGGGGCGGAGCCTATGGAAAAACGCCAGCAACGCAGAAAGGCCCACCCGAAGGTG
AGCCAGGTGATTACATTTAGGTCCTCATTAGAAAAACTCATCGAGCATCAAGTGAAACTGCAATTTAT
TCATATCAGGATTATCAATACCATATTTTTGAAAAAGCCGTTTCTGTAATGAAGGAGAAAACTCACCG
AGGCAGTTCCATAGGATGGCAAGATCCTGGTATCGGTCTGCGATTCCGACTCGTCCAACATCAATACA
ACCTATTAATTTCCCCTCGTCAAAAATAAGGTTATCAAGTGAGAAATCACCATGAGTGACGACTGAAT
CCGGTGAGAATGGCAAGAGTTTATGCATTTCTTTCCAGACTTGTTCAACAGGCCAGCCATTACGCTCGT
CATCAAAATCACTCGCACCAACCAAACCGTTATTCATTCGTGATTGCGCCTGAGCGAGACGAAATACG
CGATCGCCGTTAAAAGGACAATTACAAACAGGAATCGAATGCAACCGGCGCAGGAACACTGCCAGCG
CATCAACAATATTTTCACCTGAATCAGGATATTCTTCTAATACCTGGAATGCTGTTTTCCCTGGGATCG
CAGTGGTGAGTAACCATGCATCATCAGGAGTACGGATAAAATGCTTGATGGTCGGAAGAGGCATAAA
TTCCGTCAGCCAGTTTAGCCTGACCATCTCATCTGTAACATCATTGGCAACGCTACCTTTGCCATGTTTC
AGAAACAACTCTGGCGCATCGGGCTTCCCATACAATCGATAGATTGTCGCACCTGATTGCCCGACATT
ATCGCGAGCCCATTTATACCCATATAAATCAGCATCCATGTTGGAATTTAATCGCGGCTTCAAGCAAG
ACGTTTCCCGTTGAATATGGCTCATTTTAGCTTCCTTAGCTCCTGAAAATCTCGATAACTCAAAAAATA
CGCCCGGTAGTGATCTTATTTCATTATGGTGAAAGTTGGAACCTCTTACGTGCCGATCAAGTCAAAAGC
CTCCGGTCGGAGGCTTTTGACTTTCTGCTATGGAGGTCAGGTATGATTTAAATGGTCAGTATTGAGCCT
CAGGAAACAGCTATGACATCAAGCTGACTAGATAATCTAGCTGATCGTGGACCGATCATACGTATAAT
GCCGTAAGATCACGGGTCGCAGCACAGCTCGCGGTCCAGTAGTGATCGACACTGCTCGATCCGCTCGC
ACCCTAGGTGCGCTGCTTCGCGATGTACGGGCCAGATATACGCGTT**GACATTGATTATTGACTAGTT**
**ATTAATAGTAATCAATTACGGGGTCATTAGTTCATAGCCCATATATGGAGTTCCGCGTTACATAA**
**CTTACGGTAAATGGCCCGCCTGGCTGACCGCCCAACGACCCCCGCCCATTGACGTCAATAATGA**
**CGTATGTTCCCATAGTAACGCCAATAGGGACTTTCCATTGACGTCAATGGGTGGACTATTTACGG**
**TAAACTGCCCACTTGGCAGTACATCAAGTGTATCATATGCCAAGTACGCCCCCTATTGACGTCAA**
**TGACGGTAAATGGCCCGCCTGGCATTATGCCCAGTACATGACCTTATGGGACTTTCCTACTTGG**
**CAGTACATCTACGTATTAGTCATCGCTATTACCATGGTGATGCGGTTTTGGCAGTACATCAATGG**
**GCGTGGATAGCGGTTTGACTCACGGGGATTTCCAAGTCTCCACCCCATTGACGCAAATGGGCGG**
**TAGGCGTGTACGGTGGGAGGTCTATATAAGCAGAGCT**CTCTGGCTAACTAGAGAACCCACTGCTTA

CTGGCTTATCGAAATTAATACGACTCACTATAGGGAGACCCAAGCTGGCTAGCGTTTAAACTTAAGCT
TGCCGCCACC**ATGGAAGAGCAATACCGCCCGGAAGAGATAGAATCCAAAGTACAGCTTCATTGG**
**GATGAGAAGCGCACATTTGAAGTAACCGAAGACGAGAGCAAAGAGAAGTATTACTGCCTGTCTA**
**TCCTCCCCTATCCTTCTGGTCGACTACACATGGGCCACGTACGTAACTACACCATCGGTGACGTG**
**ATCGCCCGCTACCAGCGTATGCTGGGCAAAAACGTCCTGCAGCCGATCGGCTGGGACGCGTTTG**
**GTCTGCCTGCGGAAGGCGCGGCGGTGAAAAACAACACCGCTCCGGCACCGTGGACGTACGACA**
**ACATCGCGTATATGAAAAACCAGCTCAAAATGCTGGGCTTTGGTTATGACTGGAGCCGCGAGCT**
**GGCAACCTGTACGCCGGAATACTACCGTTGGGAACAGAAATTCTTCACCGAGCTGTATAAAAAA**
**GGCCTGGTATATAAGAAGACTTCTGCGGTCAACTGGTGTCCGAACGACCAGACCGTACTGGCGA**
**ACGAACAAGTTATCGACGGCTGCTGCTGGCGCTGCGATACCAAAGTTGAACGTAAAGAGATCCC**
**GCAGTGGTTTATCAAAATCACTGCTTACGCTGACGAGCTGCTCAACGATCTGGATAAACTGGAT**
**CACTGGCCAGACACCGTTAAAACCATGCAGCGTAACTGGATCGGTCGTTCCGAAGGCGTGGAGA**
**TCACCTTCAACGTTAACGACTATGACAACACGCTGACCGTTTACACTACCCGCCCGGACGCGTTT**
**ATGGGTTGTACCTACCTGGCGGTAGCTGCGGGTCATCCGCTGGCGCAGAAAGCGGCGGAAAAT**
**AATCCTGAACTGGCGGCCTTTATTGACGAATGCCGTAACACCAAAGTTGCCGAAGCTGAAATGG**
**CGACGATGGAGAAAAAAGGCGTCGATACTGGCTTTAAAGCGGTTCACCCATTAACGGGCGAAGA**
**AATTCCCGTTTGGGCAGCAAACTTCGTATTGATGGAGTACGGCACGGGCGCAGTTATGGCGGTA**
**CCGGGGCACGACCAGCGCGACTACGAGTTTGCCTCTAAATACGGCCTGAACATCAAACCGGTTA**
**TCCTGGCAGCTGACGGCTCTGAGCCAGATCTTTCTCAGCAAGCCCTGACTGAAAAAGGCGTGCT**
**GTTCAACTCTGGCGAGTTCAACGGTCTTGACCATGAAGCGGCCTTCAACGCCATCGCCGATAAA**
**CTGACTGCGATGGGCGTTGGCGAGCGTAAAGTGAACTACCGCCTGCGCGACTGGGGTGTTTCCC**
**GTCAGCGTTACTGGGGCGCGCCGATTCCGATGGTGACTCTAGAAGACGGTACCGTAATGCCGAC**
**CCCGGACGACCAGCTGCCGGTGATCCTGCCGGAAGATGTGGTAATGGACGGCATTACCAGCCC**
**GATTAAAGCAGATCCGGAGTGGGCGAAAACTACCGTTAACGGTATGCCAGCACTGCGTGAAACC**
**GACACTTTCGACACCTTTATGGAGTCCTCCTGGATCTATGCGCGCTACACTTGCCCGCAGTACAA**
**AGAAGGTATGCTGGATTCCGAAGCGGCTAACTACTGGCTGCCGGTGGATATCGCTATTGGTGGT**
**ATTGAACACGCCATTATGGGTCTGCTCTACTTCCGCTTCTTCCACAAACTGATGCGTGATGCAGG**
**CATGGTGAACTCTGACGAACCAGCGAAACAGTTGCTGTGTCAGGGTATGGTGCTGGCAGATGCC**
**TTCTACTATGTTGGCGAAAACGGCGAACGTAACTGGGTTTCCCCGGTTGATGCTATCGTTGAAC**
**GTGACGAGAAAGGCCGTATCGTGAAAGCGAAAGATGCGGCAGGCCATGAACTGGTTTATACCG**
**GCATGAGCAAAATGTCCAAGTCGAAGAACAACGGTATCGACCCGCAGGTGATGGTTGAACGTTA**
**CGGCGCGGACACCGTTCGTCTGTTTATGATGTTTGCTTCTCCGGCTGATATGACTCTCGAATGGC**
**AGGAATCCGGTGTGGAAGGGGCTAACCGCTTCCTGAAACGTGTCTGGAAACTGGTTTACGAGCA**
**CACAGCAAAAGGTGATGTTGCGGCACTGAACGTTGATGCGCTGACTGAAAATCAGAAAGCGCTG**
**CGTCGCGATGTGCATAAAACGATCGCTAAAGTGACCGATGATATCGGCCGTCGTCAGACCTTCA**
**ACACCGCAATTGCGGCGATTATGGAGCTGATGAACAAACTGGCGAAAGCACCAACCGATGGCGA**
**GCAGGATCGCGCTCTGATGCAGGAAGCACTGCTGGCCGTTGTCCGTATGCTTAACCCGTTCACC**
**CCGCACATCTGCTTCACGCTGTGGCAGGAACTGAAAGGCGAAGGCGATATCGACAACGCGCCGT**
**GGCCGGTTGCTGACGAAAAAGCGATGGTGGAAGACTCCACGCTGGTCGTGGTGCAGGTTAACG**
**GTAAAGTCCGTGCCAAAATCACCGTTCCGGTGGACGCAACGGAAGAACAGGTTCGCGAACGTGC**
**TGGCCAGGAACATCTGGTAGCAAAATATCTTGATGGCGTTACTGTACGTAAAGTGATTTACGTAC**
**CAGGTAAACTCCTCAATCTGGTCGTTGGCGGGCCCGTTTAA**CTCGAGTCTAGAGGGCCCGTTTAAA
CCCGCTGATCAGCCTCGACTGTGCCTTCTAGTTGCCAGCCATCTGTTGTTTGCCCCTCCCCCGTGCCTTC
CTTGACCCTGGAAGGTGCCACTCCCACTGTCCTTTCCTAATAAAATGAGGAAATTGCATCGCATTGTCT
GAGTAGGTGTCATTCTATTCTGGGGGGTGGGGTGGGGCAGGACAGCAAGGGGGAGGATTGGGAAGAC
AATAGCAGGCATGCTGGGGATGCGGTGGGCTCTATGGCTTCTGAGGCGGAAAGAACCCTAGCGGATC
GACGAGAGCAGCGCGACTGGATCTGTCGCCCGTCTCAAACGCAACCCTCCGGCGGTCGCATATCATTC
AGGACGAGCCTCAGACTCCAGCGTAACTGGACTGCAATCAACTCACTGGCTCACCTTCCGGTCCACGA
TCAGCTAGAATCAAGCTGACTAGATAA

*pIDTSmart-CMV-NvocRS*

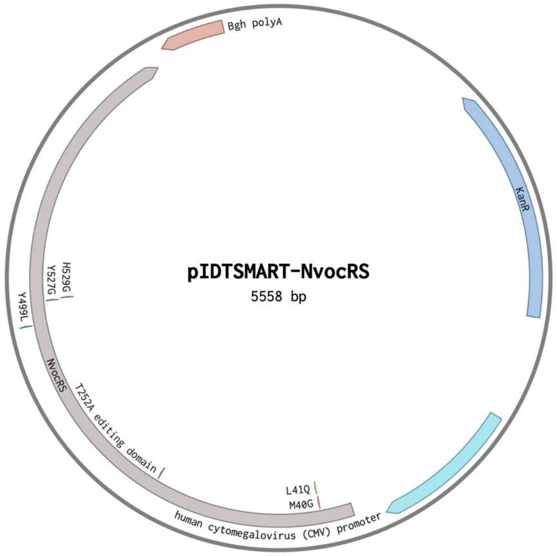


ACTGGCCGTCGTTTTACACGGGTGGGCCTTTCTTCGGTAGAAAATCAAAGGATCTTCTTGAGATCCTTT
TTTTCTGCGCGTAATCTGCTGCTTGCAAACAAAAAAACCACCGCTACCAGCGGTGGTTTGTTTGCCGGA
TCAAGAGCTACCAACTCTTTTTCCGAGGTAACTGGCTTCAGCAGAGCGCAGATACCAAATACTGTTCTT
CTAGTGTAGCCGTAGTTAGGCCACCACTTCAAGAACTCTGTAGCACCGCCTACATACCTCGCTCTGCTA
ATCCTGTTACCAGTGGCTGCTGCCAGTGGCGATAAGTCGTGTCTTACCGGGTTGGACTCAAGACGATA
GTTACCGGATAAGGCGCAGCGGTCGGGCTGAACGGGGGGTTCGTGCACACAGCCCAGCTTGGAGCGA
ACGACCTACACCGAACTGAGATACCTACAGCGTGAGCTATGAGAAAGCGCCACGCTTCCCGAAGGGA
GAAAGGCGGACAGGTATCCGGTAAGCGGCAGGGTCGGAACAGGAGAGCGCACGAGGGAGCTTCCAG
GGGGAAACGCCTGGTATCTTTATAGTCCTGTCGGGTTTCGCCACCTCTGACTTGAGCATCGATTTTTGT
GATGCTCGTCAGGGGGGCGGAGCCTATGGAAAAACGCCAGCAACGCAGAAAGGCCCACCCGAAGGTG
AGCCAGGTGATTACATTTAGGTCCTCATTAGAAAAACTCATCGAGCATCAAGTGAAACTGCAATTTAT
TCATATCAGGATTATCAATACCATATTTTTGAAAAAGCCGTTTCTGTAATGAAGGAGAAAACTCACCG
AGGCAGTTCCATAGGATGGCAAGATCCTGGTATCGGTCTGCGATTCCGACTCGTCCAACATCAATACA
ACCTATTAATTTCCCCTCGTCAAAAATAAGGTTATCAAGTGAGAAATCACCATGAGTGACGACTGAAT
CCGGTGAGAATGGCAAGAGTTTATGCATTTCTTTCCAGACTTGTTCAACAGGCCAGCCATTACGCTCGT
CATCAAAATCACTCGCACCAACCAAACCGTTATTCATTCGTGATTGCGCCTGAGCGAGACGAAATACG
CGATCGCCGTTAAAAGGACAATTACAAACAGGAATCGAATGCAACCGGCGCAGGAACACTGCCAGCG
CATCAACAATATTTTCACCTGAATCAGGATATTCTTCTAATACCTGGAATGCTGTTTTCCCTGGGATCG
CAGTGGTGAGTAACCATGCATCATCAGGAGTACGGATAAAATGCTTGATGGTCGGAAGAGGCATAAA
TTCCGTCAGCCAGTTTAGCCTGACCATCTCATCTGTAACATCATTGGCAACGCTACCTTTGCCATGTTTC
AGAAACAACTCTGGCGCATCGGGCTTCCCATACAATCGATAGATTGTCGCACCTGATTGCCCGACATT
ATCGCGAGCCCATTTATACCCATATAAATCAGCATCCATGTTGGAATTTAATCGCGGCTTCAAGCAAG
ACGTTTCCCGTTGAATATGGCTCATTTTAGCTTCCTTAGCTCCTGAAAATCTCGATAACTCAAAAAATA
CGCCCGGTAGTGATCTTATTTCATTATGGTGAAAGTTGGAACCTCTTACGTGCCGATCAAGTCAAAAGC
CTCCGGTCGGAGGCTTTTGACTTTCTGCTATGGAGGTCAGGTATGATTTAAATGGTCAGTATTGAGCCT
CAGGAAACAGCTATGACATCAAGCTGACTAGATAATCTAGCTGATCGTGGACCGATCATACGTATAAT
GCCGTAAGATCACGGGTCGCAGCACAGCTCGCGGTCCAGTAGTGATCGACACTGCTCGATCCGCTCGC
ACCCTAGGTGCGCTGCTTCGCGATGTACGGGCCAGATATACGCGTT**GACATTGATTATTGACTAGTT**
**ATTAATAGTAATCAATTACGGGGTCATTAGTTCATAGCCCATATATGGAGTTCCGCGTTACATAA**
**CTTACGGTAAATGGCCCGCCTGGCTGACCGCCCAACGACCCCCGCCCATTGACGTCAATAATGA**
**CGTATGTTCCCATAGTAACGCCAATAGGGACTTTCCATTGACGTCAATGGGTGGACTATTTACGG**
**TAAACTGCCCACTTGGCAGTACATCAAGTGTATCATATGCCAAGTACGCCCCCTATTGACGTCAA**
**TGACGGTAAATGGCCCGCCTGGCATTATGCCCAGTACATGACCTTATGGGACTTTCCTACTTGG**
**CAGTACATCTACGTATTAGTCATCGCTATTACCATGGTGATGCGGTTTTGGCAGTACATCAATGG**
**GCGTGGATAGCGGTTTGACTCACGGGGATTTCCAAGTCTCCACCCCATTGACGCAAATGGGCGG**
**TAGGCGTGTACGGTGGGAGGTCTATATAAGCAGAGCT**CTCTGGCTAACTAGAGAACCCACTGCTTA

CTGGCTTATCGAAATTAATACGACTCACTATAGGGAGACCCAAGCTGGCTAGCGTTTAAACTTAAGCT
TGCCGCCACC**ATGGAAGAGCAATACCGCCCGGAAGAGATAGAATCCAAAGTACAGCTTCATTGG**
**GATGAGAAGCGCACATTTGAAGTAACCGAAGACGAGAGCAAAGAGAAGTATTACTGCCTGTCTG**
**GGCAGCCCTATCCTTCTGGTCGACTACACATGGGCCACGTACGTAACTACACCATCGGTGACGT**
**GATCGCCCGCTACCAGCGTATGCTGGGCAAAAACGTCCTGCAGCCGATCGGCTGGGACGCGTTT**
**GGTCTGCCTGCGGAAGGCGCGGCGGTGAAAAACAACACCGCTCCGGCACCGTGGACGTACGAC**
**AACATCGCGTATATGAAAAACCAGCTCAAAATGCTGGGCTTTGGTTATGACTGGAGCCGCGAGC**
**TGGCAACCTGTACGCCGGAATACTACCGTTGGGAACAGAAATTCTTCACCGAGCTGTATAAAAA**
**AGGCCTGGTATATAAGAAGACTTCTGCGGTCAACTGGTGTCCGAACGACCAGACCGTACTGGCG**
**AACGAACAAGTTATCGACGGCTGCTGCTGGCGCTGCGATACCAAAGTTGAACGTAAAGAGATCC**
**CGCAGTGGTTTATCAAAATCACTGCTTACGCTGACGAGCTGCTCAACGATCTGGATAAACTGGA**
**TCACTGGCCAGACACCGTTAAAACCATGCAGCGTAACTGGATCGGTCGTTCCGAAGGCGTGGAG**
**ATCACCTTCAACGTTAACGACTATGACAACACGCTGACCGTTTACACTACCCGCCCGGACGCGTT**
**TATGGGTTGTACCTACCTGGCGGTAGCTGCGGGTCATCCGCTGGCGCAGAAAGCGGCGGAAAAT**
**AATCCTGAACTGGCGGCCTTTATTGACGAATGCCGTAACACCAAAGTTGCCGAAGCTGAAATGG**
**CGACGATGGAGAAAAAAGGCGTCGATACTGGCTTTAAAGCGGTTCACCCATTAACGGGCGAAGA**
**AATTCCCGTTTGGGCAGCAAACTTCGTATTGATGGAGTACGGCACGGGCGCAGTTATGGCGGTA**
**CCGGGGCACGACCAGCGCGACTACGAGTTTGCCTCTAAATACGGCCTGAACATCAAACCGGTTA**
**TCCTGGCAGCTGACGGCTCTGAGCCAGATCTTTCTCAGCAAGCCCTGACTGAAAAAGGCGTGCT**
**GTTCAACTCTGGCGAGTTCAACGGTCTTGACCATGAAGCGGCCTTCAACGCCATCGCCGATAAA**
**CTGACTGCGATGGGCGTTGGCGAGCGTAAAGTGAACTACCGCCTGCGCGACTGGGGTGTTTCCC**
**GTCAGCGTTACTGGGGCGCGCCGATTCCGATGGTGACTCTAGAAGACGGTACCGTAATGCCGAC**
**CCCGGACGACCAGCTGCCGGTGATCCTGCCGGAAGATGTGGTAATGGACGGCATTACCAGCCC**
**GATTAAAGCAGATCCGGAGTGGGCGAAAACTACCGTTAACGGTATGCCAGCACTGCGTGAAACC**
**GACACTTTCGACACCTTTATGGAGTCCTCCTGGCTGTATGCGCGCTACACTTGCCCGCAGTACAA**
**AGAAGGTATGCTGGATTCCGAAGCGGCTAACTACTGGCTGCCGGTGGATATCGGTATTGGTGGT**
**ATTGAACACGCCATTATGTTTCTGCTCTACTTCCGCTTCTTCCACAAACTGATGCGTGATGCAGG**
**CATGGTGAACTCTGACGAACCAGCGAAACAGTTGCTGTGTCAGGGTATGGTGCTGGCAGATGCC**
**TTCTACTATGTTGGCGAAAACGGCGAACGTAACTGGGTTTCCCCGGTTGATGCTATCGTTGAAC**
**GTGACGAGAAAGGCCGTATCGTGAAAGCGAAAGATGCGGCAGGCCATGAACTGGTTTATACCG**
**GCATGAGCAAAATGTCCAAGTCGAAGAACAACGGTATCGACCCGCAGGTGATGGTTGAACGTTA**
**CGGCGCGGACACCGTTCGTCTGTTTATGATGTTTGCTTCTCCGGCTGATATGACTCTCGAATGGC**
**AGGAATCCGGTGTGGAAGGGGCTAACCGCTTCCTGAAACGTGTCTGGAAACTGGTTTACGAGCA**
**CACAGCAAAAGGTGATGTTGCGGCACTGAACGTTGATGCGCTGACTGAAAATCAGAAAGCGCTG**
**CGTCGCGATGTGCATAAAACGATCGCTAAAGTGACCGATGATATCGGCCGTCGTCAGACCTTCA**
**ACACCGCAATTGCGGCGATTATGGAGCTGATGAACAAACTGGCGAAAGCACCAACCGATGGCGA**
**GCAGGATCGCGCTCTGATGCAGGAAGCACTGCTGGCCGTTGTCCGTATGCTTAACCCGTTCACC**
**CCGCACATCTGCTTCACGCTGTGGCAGGAACTGAAAGGCGAAGGCGATATCGACAACGCGCCGT**
**GGCCGGTTGCTGACGAAAAAGCGATGGTGGAAGACTCCACGCTGGTCGTGGTGCAGGTTAACG**
**GTAAAGTCCGTGCCAAAATCACCGTTCCGGTGGACGCAACGGAAGAACAGGTTCGCGAACGTGC**
**TGGCCAGGAACATCTGGTAGCAAAATATCTTGATGGCGTTACTGTACGTAAAGTGATTTACGTAC**
**CAGGTAAACTCCTCAATCTGGTCGTTGGCGGGCCCGTTTAA**CTCGAGTCTAGAGGGCCCGTTTAAA
CCCGCTGATCAGCCTCGACTGTGCCTTCTAGTTGCCAGCCATCTGTTGTTTGCCCCTCCCCCGTGCCTTC
CTTGACCCTGGAAGGTGCCACTCCCACTGTCCTTTCCTAATAAAATGAGGAAATTGCATCGCATTGTCT
GAGTAGGTGTCATTCTATTCTGGGGGGTGGGGTGGGGCAGGACAGCAAGGGGGAGGATTGGGAAGAC
AATAGCAGGCATGCTGGGGATGCGGTGGGCTCTATGGCTTCTGAGGCGGAAAGAACCCTAGCGGATC
GACGAGAGCAGCGCGACTGGATCTGTCGCCCGTCTCAAACGCAACCCTCCGGCGGTCGCATATCATTC
AGGACGAGCCTCAGACTCCAGCGTAACTGGACTGCAATCAACTCACTGGCTCACCTTCCGGTCCACGA
TCAGCTAGAATCAAGCTGACTAGATAA

*pEBV-4xU6-LtR-wt*

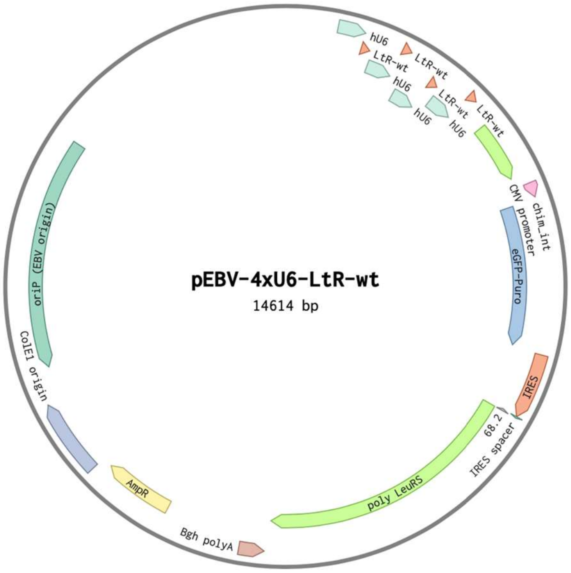


CAATAGACATCTTTATTAGACGACGCTCAGTGAATACAGGGAGTGCAGACTCCTGCCCCCTCCAACAG
CCCCCCCACCCTCATCCCCTTCATGGTCGCTGTCAGACAGATCCAGGTCTGAAAATTCCCCATCCTCCG
AACCATCCTCGTCCTCATCACCAATTACTCGCAGCCCGGAAAACTCCCGCTGAACATCCTCAAGATTTG
CGTCCTGAGCCTCAAGCCAGGCCTCAAATTCCTCGTCCCCCTTTTTGCTGGACGGTAGGGATGGGGATT
CTCGGGACCCCTCCTCTTCCTCTTCAAGGGAGCCTCAGGAAACAGCTATGACATCAAGCTGACTAGAT
AATCTAGCTGATCGTGGACCGATCATACGTATAATGCCGTAAGATCACGGGTCGCAGCACAGCTCGCG
GTCCAGTAGTGATCGACACTGCTCGATCCGCTCGCACCCCTAGGTCGGGCAGGAAGAGGGCCTATTT
CCCATGATTCCTTCATATTTGCATATACGATACAAGGCTGTTAGAGAGATAATTAGAATTAATTT
GACTGTAAACACAAAGATATTAGTACAAAATACGTGACGTAGAAAGTAATAATTTCTTGGGTAGT
TTGCAGTTTTAAAATTATGTTTTAAAATGGACTATCATATGCTTACCGTAACTTGAAAGTATTTCG
ATTTCTTGGCTTTATATATCTTGTGGAAAGGACGAAACACC**GCCCGGATGGTGGAATCGGTAGA**
**CACAAGGGATTCTAAATCCCTCGGCGTTCGCGCTGTGCGGGTTCAAGTCCCGCTCCGGGTA**TTTT
TTGCTAGGTCGGGCAGGAAGAGGGCCTATTTCCCATGATTCCTTCATATTTGCATATACGATACAA
GGCTGTTAGAGAGATAATTAGAATTAATTTGACTGTAAACACAAAGATATTAGTACAAAATACGT
GACGTAGAAAGTAATAATTTCTTGGGTAGTTTGCAGTTTTAAAATTATGTTTTAAAATGGACTAT
CATATGCTTACCGTAACTTGAAAGTATTTCGATTTCTTGGCTTTATATATCTTGTGGAAAGGACG
AAACACC**GCCCGGATGGTGGAATCGGTAGACACAAGGGATTCTAAATCCCTCGGCGTTCGCGCT**
**GTGCGGGTTCAAGTCCCGCTCCGGGTA**TTTTTTGCTAGGTCGGGCAGGAAGAGGGCCTATTTCCC
ATGATTCCTTCATATTTGCATATACGATACAAGGCTGTTAGAGAGATAATTAGAATTAATTTGAC
TGTAAACACAAAGATATTAGTACAAAATACGTGACGTAGAAAGTAATAATTTCTTGGGTAGTTTG
CAGTTTTAAAATTATGTTTTAAAATGGACTATCATATGCTTACCGTAACTTGAAAGTATTTCGATT
TCTTGGCTTTATATATCTTGTGGAAAGGACGAAACACC**GCCCGGATGGTGGAATCGGTAGACAC**
**AAGGGATTCTAAATCCCTCGGCGTTCGCGCTGTGCGGGTTCAAGTCCCGCTCCGGGTA**TTTTTTG
CTAGGTCGGGCAGGAAGAGGGCCTATTTCCCATGATTCCTTCATATTTGCATATACGATACAAGG
CTGTTAGAGAGATAATTAGAATTAATTTGACTGTAAACACAAAGATATTAGTACAAAATACGTGA
CGTAGAAAGTAATAATTTCTTGGGTAGTTTGCAGTTTTAAAATTATGTTTTAAAATGGACTATCA
TATGCTTACCGTAACTTGAAAGTATTTCGATTTCTTGGCTTTATATATCTTGTGGAAAGGACGAA
ACACC**GCCCGGATGGTGGAATCGGTAGACACAAGGGATTCTAAATCCCTCGGCGTTCGCGCTGT**
**GCGGGTTCAAGTCCCGCTCCGGGTA**TTTTTTGCTAGCGGATCGACGAGAGCAGCGCGACTGGATCT
GTCGCCCGTCTCAAACGCAACCCTCCGGCGGTCGCATATCATTCAGGACGAGCCTCAGACTCCAGCGT
AACTGGACTGCAATCAACTCACTGGCTCACCTTCCGGTCCACGATCAGCTAGAATCAAGCTGACTAGA
TAAACTGGCCGTCGTTTTACTCTAGAATAGTAATCAATTACGGGGTCATTAGTTCATAGCCCATATA
TGGAGTTCCGCGTTACATAACTTACGGTAAATGGCCCGCCTGGCTGACCGCCCAACGACCCCCG
CCCATTGACGTCAATAATGACGTATGTTCCCATAGTAACGCCAATAGGGACTTTCCATTGACGTC
AATGGGTGGAGTATTTACGGTAAACTGCCCACTTGGCAGTACATCAAGTGTATCATATGCCAAG
TCCGCCCCCTATTGACGTCAATGACGGTAAATGGCCCGCCTGGCATTATGCCCAGTACATGACC

TTACGGGACTTTCCTACTTGGCAGTACATCTACGTATTAGTCATCGCTATTACCATGGTGATGCG
GTTTTGGCAGTACACCAATGGGCGTGGATAGCGGTTTGACTCACGGGGATTTCCAAGTCTCCAC
CCCATTGACGTCAATGGGAGTTTGTTTTGGCACCAAAATCAACGGGACTTTCCAAAATGTCGTAA
TAACCCCGCCCCGTTGACGCAAATGGGCGGTAGGCGTGTACGGTGGGAGGTCTATATAAGCAGA
GGTCGTTTAGTGAACCGTCAGATCACTAGTAGCTTTATTGCGGTAGTTTATCACAGTTAAATTGCTAAC
GCAGTCAGTGCTCGACTGATCACAGGTAAGTATCAAGGTTACAAGACAGGTTTAAGGAGGCCAATAG
AAACTGGGCTTGTCGAGACAGAGAAGATTCTTGCGTTTCTGATAGGCACCTATTGGTCTTACTGACATC
CACTTTGCCTTTCTCTCCACAGGGGTAAGGCCACCATGGTGAGCAAGGGCGAGGAGCTGTTCACCG
GGGTGGTGCCCATCCTGGTCGAGCTGGACGGCGACGTAAACGGCCACAAGTTCAGCGTGTCCG
GCGAGGGCGAGGGCGATGCCACCTAGGGCAAGCTGACCCTGAAGTTCATCTGCACCACCGGCA
AGCTGCCCGTGCCCTGGCCCACCCTCGTGACCACCCTGACCTACGGCGTGCAGTGCTTCAGCCG
CTACCCCGACCACATGAAGCAGCACGACTTCTTCAAGTCCGCCATGCCCGAAGGCTACGTCCAG
GAGCGCACCATCTTCTTCAAGGACGACGGCAACTACAAGACCCGCGCCGAGGTGAAGTTCGAGG
GCGACACCCTGGTGAACCGCATCGAGCTGAAGGGCATCGACTTCAAGGAGGACGGCAACATCCT
GGGGCACAAGCTGGAGTACAACTACAACAGCCACAACGTCTATATCATGGCCGACAAGCAGAAG
AACGGCATCAAGGTGAACTTCAAGATCCGCCACAACATCGAGGACGGCAGCGTGCAGCTCGCCG
ACCACTACCAGCAGAACACCCCCATCGGCGACGGCCCCGTGCTGCTGCCCGACAACCACTACCT
GAGCACCCAGTCCGCCCTGAGCAAAGACCCCAACGAGAAGCGCGATCACATGGTCCTGCTGGA
GTTCGTGACCGCCGCCGGGATCACTCTCGGCATGGACGAGCTGTACAAGTCCGGACTCAGATCT
AGGAGACGACCTTCCATGACCGAGTACAAGCCCACGGTGCGCCTCGCCACCCGCGACGACGTCC
CCCGGGCCGTACGCACCCTCGCCGCCGCGTTCGCCGACTACCCCGCCACGCGCCACACCGTCGA
CCCGGACCGCCACATCGAGCGGGTCACCGAGCTGCAAGAACTCTTCCTCACGCGCGTCGGGCTC
GACATCGGCAAGGTGTGGGTCGCGGACGACGGCGCCGCGGTGGCGGTCTGGACCACGCCGGAG
AGCGTCGAAGCGGGGGCGGTGTTCGCCGAGATCGGCCCGCGCATGGCCGAGTTGAGCGGTTCC
CGGCTGGCCGCGCAGCAACAGATGGAAGGCCTCCTGGCGCCGCACCGGCCCAAGGAGCCCGCG
TGGTTCCTGGCCACCGTCGGCGTCTCGCCCGACCACCAGGGCAAGGGTCTGGGCAGCGCCGTC
GTGCTCCCCGGAGTGGAGGCGGCCGAGCGCGCCGGGGTGCCCGCCTTCCTGGAGACCTCCGCG
CCCCGCAACCTCCCCTTCTACGAGCGGCTCGGCTTCACCGTCACCGCCGACGTCGAGTGCCCGA
AGGACCGCGCGACCTGGTGCATGACCCGCAAGCCCGGTGCCCGGGATCCACCGGATCTAGATA
ACTGATCTAGAGTACCCGGGCTAGGATCCGCCCCTCTCCCTCCCCCCCCCCTAACGTTACTGGCCGAA
GCCGCTTGGAATAAGGCCGGTGTGCGTTTGTCTATATGTTATTTTCCACCATATTGCCGTCTTTTGGCA
ATGTGAGGGCCCGGAAACCTGGCCCTGTCTTCTTGACGAGCATTCCTAGGGGTCTTTCCCCTCTCGCCA
AAGGAATGCAAGGTCTGTTGAATGTCGTGAAGGAAGCAGTTCCTCTGGAAGCTTCTTGAAGACAAACA
ACGTCTGTAGCGACCCTTTGCAGGCAGCGGAACCCCCCACCTGGCGACAGGTGCCTCTGCGGCCAAAA
GCCACGTGTATAAGATACACCTGCAAAGGCGGCACAACCCCAGTGCCACGTTGTGAGTTGGATAGTTG
TGGAAAGAGTCAAATGGCTCTCCTCAAGCGTATTCAACAAGGGGCTGAAGGATGCCCAGAAGGTACC
CCATTGTATGGGATCTGATCTGGGGCCTCGGTGCACATGCTTTACATGTGTTTAGTCGAGGTTAAAAA
ACGTCTAGGCCCCCCGAACCACGGGGACGTGGTTTTCCTTTGAAAAACACGATGATAATATGGCCACA
ACCATGGCGTCCGGAATGGAAGAGCAATACCGCCCGGAAGAGATAGAATCCAAAGTACAGCTTCA
TTGGGATGAGAAGCGCACATTTGAAGTAACCGAAGACGAGAGCAAAGAGAAGTATTACTGCCTG
TCTATCCTCCCCTATCCTTCTGGTCGACTACACATGGGCCACGTACGTAACTACACCATCGGTGA
CGTGATCGCCCGCTACCAGCGTATGCTGGGCAAAAACGTCCTGCAGCCGATCGGCTGGGACGC
GTTTGGTCTGCCTGCGGAAGGCGCGGCGGTGAAAAACAACACCGCTCCGGCACCGTGGACGTA
CGACAACATCGCGTATATGAAAAACCAGCTCAAAATGCTGGGCTTTGGTTATGACTGGAGCCGC
GAGCTGGCAACCTGTACGCCGGAATACTACCGTTGGGAACAGAAATTCTTCACCGAGCTGTATA
AAAAAGGCCTGGTATATAAGAAGACTTCTGCGGTCAACTGGTGTCCGAACGACCAGACCGTACT
GGCGAACGAACAAGTTATCGACGGCTGCTGCTGGCGCTGCGATACCAAAGTTGAACGTAAAGAG
ATCCCGCAGTGGTTTATCAAAATCACTGCTTACGCTGACGAGCTGCTCAACGATCTGGATAAACT
GGATCACTGGCCAGACACCGTTAAAACCATGCAGCGTAACTGGATCGGTCGTTCCGAAGGCGTG
GAGATCACCTTCAACGTTAACGACTATGACAACACGCTGACCGTTTACACTACCCGCCCGGACG
CGTTTATGGGTTGTACCTACCTGGCGGTAGCTGCGGGTCATCCGCTGGCGCAGAAAGCGGCGGA
AAATAATCCTGAACTGGCGGCCTTTATTGACGAATGCCGTAACACCAAAGTTGCCGAAGCTGAA
ATGGCGACGATGGAGAAAAAGGCGTCGATACTGGCTTTAAAGCGGTTCACCCATTAACGGGCG
AAGAAATTCCCGTTTGGGCAGCAAACTTCGTATTGATGGAGTACGGCACGGGCGCAGTTATGGC
GGTACCGGGGCACGACCAGCGCGACTACGAGTTTGCCTCTAAATACGGCCTGAACATCAAACCG
GTTATCCTGGCAGCTGACGGCTCTGAGCCAGATCTTTCTCAGCAAGCCCTGACTGAAAAAGGCG

**TGCTGTTCAACTCTGGCGAGTTCAACGGTCTTGACCATGAAGCGGCCTTCAACGCCATCGCCGA**
**TAAACTGACTGCGATGGGCGTTGGCGAGCGTAAAGTGAACTACCGCCTGCGCGACTGGGGTGTT**
**TCCCGTCAGCGTTACTGGGGCGCGCCGATTCCGATGGTGACTCTAGAAGACGGTACCGTAATGC**
**CGACCCCGGACGACCAGCTGCCGGTGATCCTGCCGGAAGATGTGGTAATGGACGGCATTACCA**
**GCCCGATTAAAGCAGATCCGGAGTGGGCGAAAACTACCGTTAACGGTATGCCAGCACTGCGTGA**
**AACCGACACTTTCGACACCTTTATGGAGTCCTCCTGGATCTATGCGCGCTACACTTGCCCGCAGT**
**ACAAAGAAGGTATGCTGGATTCCGAAGCGGCTAACTACTGGCTGCCGGTGGATATCGCTATTGG**
**TGGTATTGAACACGCCATTATGGGTCTGCTCTACTTCCGCTTCTTCCACAAACTGATGCGTGATG**
**CAGGCATGGTGAACTCTGACGAACCAGCGAAACAGTTGCTGTGTCAGGGTATGGTGCTGGCAGA**
**TGCCTTCTACTATGTTGGCGAAAACGGCGAACGTAACTGGGTTTCCCCGGTTGATGCTATCGTT**
**GAACGTGACGAGAAAGGCCGTATCGTGAAAGCGAAAGATGCGGCAGGCCATGAACTGGTTTATA**
**CCGGCATGAGCAAAATGTCCAAGTCGAAGAACAACGGTATCGACCCGCAGGTGATGGTTGAACG**
**TTACGGCGCGGACACCGTTCGTCTGTTTATGATGTTTGCTTCTCCGGCTGATATGACTCTCGAAT**
**GGCAGGAATCCGGTGTGGAAGGGGCTAACCGCTTCCTGAAACGTGTCTGGAAACTGGTTTACGA**
**GCACACAGCAAAAGGTGATGTTGCGGCACTGAACGTTGATGCGCTGACTGAAAATCAGAAAGCG**
**CTGCGTCGCGATGTGCATAAAACGATCGCTAAAGTGACCGATGATATCGGCCGTCGTCAGACCT**
**TCAACACCGCAATTGCGGCGATTATGGAGCTGATGAACAAACTGGCGAAAGCACCAACCGATGG**
**CGAGCAGGATCGCGCTCTGATGCAGGAAGCACTGCTGGCCGTTGTCCGTATGCTTAACCCGTTC**
**ACCCCGCACATCTGCTTCACGCTGTGGCAGGAACTGAAAGGCGAAGGCGATATCGACAACGCGC**
**CGTGGCCGGTTGCTGACGAAAAAGCGATGGTGGAAGACTCCACGCTGGTCGTGGTGCAGGTTA**
**ACGGTAAAGTCCGTGCCAAAATCACCGTTCCGGTGGACGCAACGGAAGAACAGGTTCGCGAACG**
**TGCTGGCCAGGAACATCTGGTAGCAAAATATCTTGATGGCGTTACTGTACGTAAAGTGATTTAC**
**GTACCAGGTAAACTCCTCAATCTGGTCGTTGGCGGGCCCGTTTAA**CTCGAGTCTAGAGGGCCCGTT
TAAACCCGCTGATCAGCCTCGACTGTGCCTTCTAGTTGCCAGCCATCTGTTGTTTGCCCCTCCCCCGTG
CCTTCCTTGACCCTGGAAGGTGCCACTCCCACTGTCCTTTCCTAATAAAATGAGGAAATTGCATCGCAT
TGTCTGAGTAGGTGTCATTCTATTCTGGGGGGTGGGGTGGGGCAGGACAGCAAGGGGGAGGATTGGG
AAGACAATAGCAGGCATGCTGGGGATGCGGTGGGCTCTATGGCTTCTGAGGCGGTGATGGTACCGAG
CTCGAATACGTAGATGTACTGCCAAGTAGGAAAGTCCCATAAGGTCATGTACTGGGCATAATGCCAGG
CGGGCCATTTACCGTCATTGACGTCAATAGGGGGCGTACTTGGCATATGATACACTTGATGTACTGCC
AAGTGGGCAGTTTACCGTAAATACTCCACCCATTGACGTCAATGGAAAGTCCCTATTGGCGTTACTAT
GGGAACATACGTCATTATTGACGTCAATGGGCGGGGGTCGTTGGGCGGTCAGCCAGGCGGGCCATTTA
CCGTAAGTTATGTAACGCGGAACTCCATATATGGGCTATGAACTAATGACCCCGTAATTGATTACTATT
AATAACTAGTCAATAATCAATGTCGACCCAGGTGGCACTTTTCGGGGAAATGTGCGCGGAACCCCTAT
TTGTTTATTTTTCTAAATACATTCAAATATGTATCCGCTCATGAGACAATAACCCTGATAAATGCTTCA
ATAATATTGAAAAAGGAAGAGTATGAGTATTCAACATTTCCGTGTCGCCCTTATTCCCTTTTTTGCGGC
ATTTTGCCTTCCTGTTTTTGCTCACCCAGAAACGCTGGTGAAAGTAAAAGATGCTGAAGATCAGTTGGG
TGCACGAGTGGGTTACATCGAACTGGATCTCAACAGCGGTAAGATCCTTGAGAGTTTTCGCCCCGAAG
AACGTTTTCCAATGATGAGCACTTTTAAAGTTCTGCTATGTGGCGCGGTATTATCCCGTATTGACGCCG
GGCAAGAGCAACTCGGTCGCCGCATACACTATTCTCAGAATGACTTGGTTGAGTACTCACCAGTCACA
GAAAAGCATCTTACGGATGGCATGACAGTAAGAGAATTATGCAGTGCTGCCATAACCATGAGTGATA
ACACTGCGGCCAACTTACTTCTGACAACGATCGGAGGACCGAAGGAGCTAACCGCTTTTTTGCACAAC
ATGGGGGATCATGTAACTCGCCTTGATCGTTGGGAACCGGAGCTGAATGAAGCCATACCAAACGACG
AGCGTGACACCACGATGCCTGTAGCAATGGCAACAACGTTGCGCAAACTATTAACTGGCGAACTACTT
ACTCTAGCTTCCCGGCAACAATTAATAGACTGGATGGAGGCGGATAAAGTTGCAGGACCACTTCTGCG
CTCGGCCCTTCCGGCTGGCTGGTTTATTGCTGATAAATCTGGAGCCGGTGAGCGTGGGTCTCGCGGTAT
CATTGCAGCACTGGGGCCAGATGGTAAGCCCTCCCGTATCGTAGTTATCTACACGACGGGGAGTCAGG
CAACTATGGATGAACGAAATAGACAGATCGCTGAGATAGGTGCCTCACTGATTAAGCATTGGTAACTG
TCAGACCAAGTTTACTCATATATACTTTAGATTGATTTAAAACTTCATTTTTAATTTAAAAGGATCTAG
GTGAAGATCCTTTTTGATAATCTCATGACCAAAATCCCTTAACGTGAGTTTTCGTTCCACTGAGCGTCA
GACCCCGTAGAAAAGATCAAAGGATCTTCTTGAGATCCTTTTTTTCTGCGCGTAATCTGCTGCTTGCAA
ACAAAAAAACCACCGCTACCAGCGGTGGTTTGTTTGCCGGATCAAGAGCTACCAACTCTTTTTCCGAA
GGTAACTGGCTTCAGCAGAGCGCAGATACCAAATACTGTTCTTCTAGTGTAGCCGTAGTTAGGCCACC
ACTTCAAGAACTCTGTAGCACCGCCTACATACCTCGCTCTGCTAATCCTGTTACCAGTGGCTGCTGCCA
GTGGCGATAAGTCGTGTCTTACCGGGTTGGACTCAAGACGATAGTTACCGGATAAGGCGCAGCGGTCG
GGCTGAACGGGGGGTTCGTGCACACAGCCCAGCTTGGAGCGAACGACCTACACCGAACTGAGATACC
TACAGCGTGAGCTATGAGAAAGCGCCACGCTTCCCGAAGGGAGAAAGGCGGACAGGTATCCGGTAAG

CGGCAGGGTCGGAACAGGAGAGCGCACGAGGGAGCTTCCAGGGGGAAACGCCTGGTATCTTTATAGT
CCTGTCGGGTTTCGCCACCTCTGACTTGAGCGTCGATTTTTGTGATGCTCGTCAGGGGGGCGGAGCCTA
TGGAAAAACGCCAGCAACGCGGCCTTTTTACGGTTCCTGGCCTTTTGCTGGCCTTTTGCTCACATGTTC
TTTCCTGCGTTATCCCCTGATTCTGTGGATAACCGTATTACCGCCTTTGAGTGAGCTGATACCGCTCGC
CGCAGCCGAACGACCGAGCGCAGCGAGTCAGTGAGCGAGGAAGCGGAAGAGCGCCCAATACGCAAA
CCGCCTCTCCCCGCGCGTTGGCCGATTCATTAATGCAGCGGATCCGCTAGCGTCGACTCTAGAGGATC
GATGCCCCGCCCCGGACGAACTAAACCTGACTACGACATCTCTGCCCCTTCTTCGCGGGGCAGTGCAT
GTAATCCCTTCAGTTGGTTGGTACAACTTGCCAACTGGGCCCTGTTCCACATGTGACACGGGGGGGGA
CCAAACACAAAGGGGTTCTCTGACTGTAGTTGACATCCTTATAAATGGATGTGCACATTTGCCAACAC
TGAGTGGCTTTCATCCTGGAGCAGACTTTGCAGTCTGTGGACTGCAACACAACATTGCCTTTATGTGTA
ACTCTTGGCTGAAGCTCTTACACCAATGCTGGGGGACATGTACCTCCCAGGGGCCCAGGAAGACTACG
GGAGGCTACACCAACGTCAATCAGAGGGGCCTGTGTAGCTACCGATAAGCGGACCCTCAAGAGGGCA
TTAGCAATAGTGTTTATAAGGCCCCCTTGTTAACCCTAAACGGGTAGCATATGCTTCCCGGGTAGTAGT
ATATACTATCCAGACTAACCCTAATTCAATAGCATATGTTACCCAACGGGAAGCATATGCTATCGAAT
TAGGGTTAGTAAAAGGGTCCTAAGGAACAGCGATATCTCCCACCCCATGAGCTGTCACGGTTTTATTT
ACATGGGGTCAGGATTCCACGAGGGTAGTGAACCATTTTAGTCACAAGGGCAGTGGCTGAAGATCAA
GGAGCGGGCAGTGAACTCTCCTGAATCTTCGCCTGCTTCTTCATTCTCCTTCGTTTAGCTAATAGAATA
ACTGCTGAGTTGTGAACAGTAAGGTGTATGTGAGGTGCTCGAAAACAAGGTTTCAGGTGACGCCCCCA
GAATAAAATTTGGACGGGGGGTTCAGTGGTGGCATTGTGCTATGACACCAATATAACCCTCACAAACC
CCTTGGGCAATAAATACTAGTGTAGGAATGAAACATTCTGAATATCTTTAACAATAGAAATCCATGGG
GTGGGGACAAGCCGTAAAGACTGGATGTCCATCTCACACGAATTTATGGCTATGGGCAACACATAATC
CTAGTGCAATATGATACTGGGGTTATTAAGATGTGTCCCAGGCAGGGACCAAGACAGGTGAACCATGT
TGTTACACTCTATTTGTAACAAGGGGAAAGAGAGTGGACGCCGACAGCAGCGGACTCCACTGGTTGTC
TCTAACACCCCCGAAAATTAAACGGGGCTCCACGCCAATGGGGCCCATAAACAAAGACAAGTGGCCA
CTCTTTTTTTTGAAATTGTGGAGTGGGGGCACGCGTCAGCCCCCACACGCCGCCCTGCGGTTTTGGACT
GTAAAATAAGGGTGTAATAACTTGGCTGATTGTAACCCCGCTAACCACTGCGGTCAAACCACTTGCCC
ACAAAACCACTAATGGCACCCCGGGGAATACCTGCATAAGTAGGTGGGCGGGCCAAGATAGGGGCGC
GATTGCTGCGATCTGGAGGACAAATTACACACACTTGCGCCTGAGCGCCAAGCACAGGGTTGTTGGTC
CTCATATTCACGAGGTCGCTGAGAGCACGGTGGGCTAATGTTGCCATGGGTAGCATATACTACCCAAA
TATCTGGATAGCATATGCTATCCTAATCTATATCTGGGTAGCATAGGCTATCCTAATCTATATCTGGGT
AGCATATGCTATCCTAATCTATATCTGGGTAGTATATGCTATCCTAATTTATATCTGGGTAGCATAGGC
TATCCTAATCTATATCTGGGTAGCATATGCTATCCTAATCTATATCTGGGTAGTATATGCTATCCTAATC
TGTATCCGGGTAGCATATGCTATCCTAATAGAGATTAGGGTAGTATATGCTATCCTAATTTATATCTGG
GTAGCATATACTACCCAAATATCTGGATAGCATATGCTATCCTAATCTATATCTGGGTAGCATATGCTA
TCCTAATCTATATCTGGGTAGCATAGGCTATCCTAATCTATATCTGGGTAGCATATGCTATCCTAATCT
ATATCTGGGTAGTATATGCTATCCTAATTTATATCTGGGTAGCATAGGCTATCCTAATCTATATCTGGG
TAGCATATGCTATCCTAATCTATATCTGGGTAGTATATGCTATCCTAATCTGTATCCGGGTAGCATATG
CTATCCTCATGCATATACAGTCAGCATATGATACCCAGTAGTAGAGTGGGAGTGCTATCCTTTGCATAT
GCCGCCACCTCCCAAGGGGGCGTGAATTTTCGCTGCTTGTCCTTTTCCTGCTGGTTGCTCCCATTCTTAG
GTGAATTTAAGGAGGCCAGGCTAAAGCCGTCGCATGTCTGATTGCTCACCAGGTAAATGTCGCTAATG
TTTTCCAACGCGAGAAGGTGTTGAGCGCGGAGCTGAGTGACGTGACAACATGGGTATGCCCAATTGCC
CCATGTTGGGAGGACGAAAATGGTGACAAGACAGATGGCCAGAAATACACCAACAGCACGCATGATG
TCTACTGGGGATTTATTCTTTAGTGCGGGGGAATACACGGCTTTTAATACGATTGAGGGCGTCTCCTAA
CAAGTTACATCACTCCTGCCCTTCCTCACCCTCATCTCCATCACCTCCTTCATCTCCGTCATCTCCGTCA
TCACCCTCCGCGGCAGCCCCTTCCACCATAGGTGGAAACCAGGGAGGCAAATCTACTCCATCGTCAAA
GCTGCACACAGTCACCCTGATATTGCAGGTAGGAGCGGGCTTTGTCATAACAAGGTCCTTAATCGCAT
CCTTCAAAACCTCAGCAAATATATGAGTTTGTAAAAAGACCATGAAATAACAGACAATGGACTCCCTT
AGCGGGCCAGGTTGTGGGCCGGGTCCAGGGGCCATTCCAAAGGGGAGACGACTCAATGGTGTAAGAC
GACATTGTGGAATAGCAAGGGCAGTTCCTCGCCTTAGGTTGTAAAGGGAGGTCTTACTACCTCCATAT
ACGAACACACCGGCGACCCAAGTTCCTTCGTCGGTAGTCCTTTCTACGTGACTCCTAGCCAGGAGAGC
TCTTAAACCTTCTGCAATGTTCTCAAATTTCGGGTTGGAACCTCCTTGACCACGATGCTTTCCAAACCA
CCCTCCTTTTTTGCGCCTGCCTCCATCACCCTGACCCCGGGGTCCAGTGCTTGGGCCTTCTCCTGGGTCA
TCTGCGGGGCCCTGCTCTATCGCTCCCGGGGGCACGTCAGGCTCACCATCTGGGCCACCTTCTTGGTGG
TATTCAAAATAATCGGCTTCCCCTACAGGGTGGAAAAATGGCCTTCTACCTGGAGGGGGCCTGCGCGG
TGGAGACCCGGATGATGATGACTGACTACTGGGACTCCTGGGCCTCTTTTCTCCACGTCCACGACCTCT
CCCCCTGGCTCTTTCACGACTTCCCCCCCTGGCTCTTTCACGTCCTCTACCCCGGCGGCCTCCACTACCT

CCTCGACCCCGGCCTCCACTACCTCCTCGACCCCGGCCTCCACTGCCTCCTCGACCCCGGCCTCCACCT
CCTGCTCCTGCCCCTCCTGCTCCTGCCCCTCCTCCTGCTCCTGCCCCTCCTGCCCCTCCTGCTCCTGCCC
CTCCTGCCCCTCCTGCTCCTGCCCCTCCTGCCCCTCCTGCTCCTGCCCCTCCTGCCCCTCCTCCTGCTCCT
GCCCCTCCTGCCCCTCCTCCTGCTCCTGCCCCTCCTGCCCCTCCTGCTCCTGCCCCTCCTGCCCCTCCTG
CTCCTGCCCCTCCTGCCCCTCCTGCTCCTGCCCCTCCTGCTCCTGCCCCTCCTGCTCCTGCCCCTCCTGCT
CCTGCCCCTCCTGCCCCTCCTGCCCCTCCTCCTGCTCCTGCCCCTCCTGCTCCTGCCCCTCCTGCCCCTC
CTGCCCCTCCTGCTCCTGCCCCTCCTCCTGCTCCTGCCCCTCCTGCCCCTCCTGCCCCTCCTCCTGCTCCT
GCCCCTCCTGCCCCTCCTCCTGCTCCTGCCCCTCCTCCTGCTCCTGCCCCTCCTGCCCCTCCTGCCCCTC
CTCCTGCTCCTGCCCCTCCTGCCCCTCCTCCTGCTCCTGCCCCTCCTCCTGCTCCTGCCCCTCCTGCCCCT
CCTGCCCCTCCTCCTGCTCCTGCCCCTCCTCCTGCTCCTGCCCCTCCTGCCCCTCCTGCCCCTCCTGCCC
CTCCTCCTGCTCCTGCCCCTCCTCCTGCTCCTGCCCCTCCTGCTCCTGCCCCTCCCGCTCCTGCTCCTGCT
CCTGTTCCACCGTGGGTCCCTTTGCAGCCAATGCAACTTGGACGTTTTTGGGGTCTCCGGACACCATCT
CTATGTCTTGGCCCTGATCCTGAGCCGCCCGGGGCTCCTGGTCTTCCGCCTCCTCGTCCTCGTCCTCTTC
CCCGTCCTCGTCCATGGTTATCACCCCCTCTTCTTTGAGGTCCACTGCCGCCGGAGCCTTCTGGTCCAG
ATGTGTCTCCCTTCTCTCCTAGGCCATTTCCAGGTCCTGTACCTGGCCCCTCGTCAGACATGATTCACAC
TAAAAGAGAT

*pIDTSmart-CMV-PLRS1-AAV-RC2-TAG454*

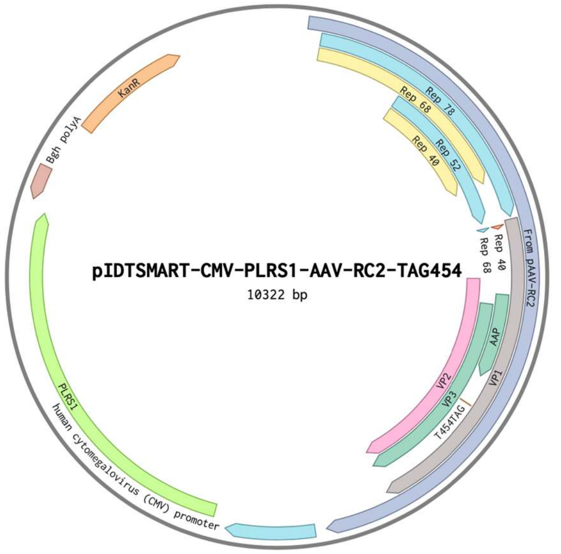


CCCGTGTAAAACGACGGCCAGTTTATCTAGTCAGCTTGATTCTAGCTGATCGTGGACCGGAAGGTGAG
CCAGTGAGTTGATTGCAGTCCAGTTACGCTGGAGTCTGAGGCTCGTCCTGAATGATATGCGACCGCCG
GAGGGTTGCGTTTGAGACGGGCGACAGATCCAGTCGCGCTGCTCTCGTCGATCCGCTAGGGCGGCCGC
TCTAGAACTAGTGGATCCCCCGGAAGATCAGAAGTTCCTATTCCGAAGTTCCTATTCTCTAGAAAGTAT
AGGAACTTCTGATCTGCGCAGCCGCCATGCCGGGGTTTTACGAGATTGTGATTAAGGTCCCCAGCGAC
CTTGACGAGCATCTGCCCGGCATTTCTGACAGCTTTGTGAACTGGGTGGCCGAGAAGGAATGGGAGTT
GCCGCCAGATTCTGACATGGATCTGAATCTGATTGAGCAGGCACCCCTGACCGTGGCCGAGAAGCTGC
AGCGCGACTTTCTGACGGAATGGCGCCGTGTGAGTAAGGCCCCGGAGGCCCTTTTCTTTGTGCAATTTG
AGAAGGGAGAGAGCTACTTCCACATGCACGTGCTCGTGGAAACCACCGGGGTGAAATCCATGGTTTTG
GGACGTTTCCTGAGTCAGATTCGCGAAAAACTGATTCAGAGAATTTACCGCGGGATCGAGCCGACTTT
GCCAAACTGGTTCGCGGTCACAAAGACCAGAAATGGCGCCGGAGGCGGGAACAAGGTGGTGGATGAG
TGCTACATCCCCAATTACTTGCTCCCCAAAACCCAGCCTGAGCTCCAGTGGGCGTGGACTAATATGGA
ACAGTATTTAAGCGCCTGTTTGAATCTCACGGAGCGTAAACGGTTGGTGGCGCAGCATCTGACGCACG
TGTCGCAGACGCAGGAGCAGAACAAAGAGAATCAGAATCCCAATTCTGATGCGCCGGTGATCAGATC
AAAAACTTCAGCCAGGTACATGGAGCTGGTCGGGTGGCTCGTGGACAAGGGGATTACCTCGGAGAAG
CAGTGGATCCAGGAGGACCAGGCCTCATACATCTCCTTCAATGCGGCCTCCAACTCGCGGTCCCAAAT
CAAGGCTGCCTTGGACAATGCGGGAAAGATTATGAGCCTGACTAAAACCGCCCCCGACTACCTGGTGG
GCCAGCAGCCCGTGGAGGACATTTCCAGCAATCGGATTTATAAAATTTTGGAACTAAACGGGTACGAT
CCCCAATATGCGGCTTCCGTCTTTCTGGGATGGGCCACGAAAAAGTTCGGCAAGAGGAACACCATCTG
GCTGTTTGGGCCTGCAACTACCGGGAAGACCAACATCGCGGAGGCCATAGCCCACACTGTGCCCTTCT
ACGGGTGCGTAAACTGGACCAATGAGAACTTTCCCTTCAACGACTGTGTCGACAAGATGGTGATCTGG
TGGGAGGAGGGGAAGATGACCGCCAAGGTCGTGGAGTCGGCCAAAGCCATTCTCGGAGGAAGCAAGG
TGCGCGTGGACCAGAAATGCAAGTCCTCGGCCCAGATAGACCCGACTCCCGTGATCGTCACCTCCAAC
ACCAACATGTGCGCCGTGATTGACGGGAACTCAACGACCTTCGAACACCAGCAGCCGTTGCAAGACCG
GATGTTCAAATTTGAACTCACCCGCCGTCTGGATCATGACTTTGGGAAGGTCACCAAGCAGGAAGTCA
AAGACTTTTTCCGGTGGGCAAAGGATCACGTGGTTGAGGTGGAGCATGAATTCTACGTCAAAAAGGGT
GGAGCCAAGAAAAGACCCGCCCCCAGTGACGCAGATATAAGTGAGCCCAAACGGGTGCGCGAGTCAG
TTGCGCAGCCATCGACGTCAGACGCGGAAGCTTCGATCAACTACGCAGACAGGTACCAAAACAAATG
TTCTCGTCACGTGGGCATGAATCTGATGCTGTTTCCCTGCAGACAATGCGAGAGAATGAATCAGAATT
CAAATATCTGCTTCACTCACGGACAGAAAGACTGTTTAGAGTGCTTTCCCGTGTCAGAATCTCAACCCG
TTTCTGTCGTCAAAAAGGCGTATCAGAAACTGTGCTACATTCATCATATCATGGGAAAGGTGCCAGAC
GCTTGCACTGCCTGCGATCTGGTCAATGTGGATTTGGATGACTGCATCTTTGAACAATAAATGATTTAA
ATCAGGTATGGCTGCCGATGGTTATCTTCCAGATTGGCTCGAGGACACTCTCTCTGAAGGAATAAGAC
AGTGGTGGAAGCTCAAACCTGGCCCACCACCACCAAAGCCCGCAGAGCGGCATAAGGACGACAGCAG
GGGTCTTGTGCTTCCTGGGTACAAGTACCTCGGACCCTTCAACGGACTCGACAAGGGAGAGCCGGTCA
ACGAGGCAGACGCCGCGGCCCTCGAGCACGACAAAGCCTACGACCGGCAGCTCGACAGCGGAGACAA

CCCGTACCTCAAGTACAACCACGCCGACGCGGAGTTTCAGGAGCGCCTTAAAGAAGATACGTCTTTTG
GGGGCAACCTCGGACGAGCAGTCTTCCAGGCGAAAAAGAGGGTTCTTGAACCTCTGGGCCTGGTTGAG
GAACCTGTTAAGACGGCTCCGGGAAAAAAGAGGCCGGTAGAGCACTCTCCTGTGGAGCCAGACTCCT
CCTCGGGAACCGGAAAGGCGGGCCAGCAGCCTGCAAGAAAAAGATTGAATTTTGGTCAGACTGGAGA
CGCAGACTCAGTACCTGACCCCCAGCCTCTCGGACAGCCACCAGCAGCCCCCTCTGGTCTGGGAACTA
ATACGATGGCTACAGGCAGTGGCGCACCAATGGCAGACAATAACGAGGGCGCCGACGGAGTGGGTAA
TTCCTCGGGAAATTGGCATTGCGATTCCACATGGATGGGCGACAGAGTCATCACCACCAGCACCCGAA
CCTGGGCCCTGCCCACCTACAACAACCACCTCTACAAACAAATTTCCAGCCAATCAGGAGCCTCGAAC
GACAATCACTACTTTGGCTACAGCACCCCTTGGGGGTATTTTGACTTCAACAGATTCCACTGCCACTTT
TCACCACGTGACTGGCAAAGACTCATCAACAACAACTGGGGATTCCGACCCAAGAGACTCAACTTCAA
GCTCTTTAACATTCAAGTCAAAGAGGTCACGCAGAATGACGGTACGACGACGATTGCCAATAACCTTA
CCAGCACGGTTCAGGTGTTTACTGACTCGGAGTACCAGCTCCCGTACGTCCTCGGCTCGGCGCATCAA
GGATGCCTCCCGCCGTTCCCAGCAGACGTCTTCATGGTGCCACAGTATGGATACCTCACCCTGAACAA
CGGGAGTCAGGCAGTAGGACGCTCTTCATTTTACTGCCTGGAGTACTTTCCTTCTCAGATGCTGCGTAC
CGGAAACAACTTTACCTTCAGCTACACTTTTGAGGACGTTCCTTTCCACAGCAGCTACGCTCACAGCCA
GAGTCTGGACCGTCTCATGAATCCTCTCATCGACCAGTACCTGTATTACTTGAGCAGAACAAACACTCC
AAGTGGATAGACCACGCAGTCAAGGCTTCAGTTTTCTCAGGCCGGAGCGAGTGACATTCGGGACCAGT
CTAGGAACTGGCTTCCTGGACCCTGTTACCGCCAGCAGCGAGTATCAAAGACATCTGCGGATAACAAC
AACAGTGAATACTCGTGGACTGGAGCTACCAAGTACCACCTCAATGGCAGAGACTCTCTGGTGAATCC
GGGCCCGGCCATGGCAAGCCACAAGGACGATGAAGAAAAGTTTTTCCTCAGAGCGGGGTTCTCATCT
TTGGGAAGCAAGGCTCAGAGAAAACAAATGTGGACATTGAAAAGGTCATGATTACAGACGAAGAGGA
AATCAGGACAACCAATCCCGTGGCTACGGAGCAGTATGGTTCTGTATCTACCAACCTCCAGAGAGGCA
ACAGACAAGCAGCTACCGCAGATGTCAACACACAAGGCGTTCTTCCAGGCATGGTCTGGCAGGACAG
AGATGTGTACCTTCAGGGGCCCATCTGGGCAAAGATTCCACACACGGACGGACATTTTCACCCCTCTC
CCCTCATGGGTGGATTCGGACTTAAACACCCTCCTCCACAGATTCTCATCAAGAACACCCCGGTACCTG
CGAATCCTTCGACCACCTTCAGTGCGGCAAAGTTTGCTTCCTTCATCACACAGTACTCCACGGGACAGG
TCAGCGTGGAGATCGAGTGGGAGCTGCAGAAGGAAAACAGCAAACGCTGGAATCCCGAAATTCAGTA
CACTTCCAACTACAACAAGTCTGTTAATGTGGACTTTACTGTGGACACTAATGGCGTGTATTCAGAGCC
TCGCCCCATTGGCACCAGATACCTGACTCGTAATCTGTAATTGCTTGTTAATCAATAAACCGTTTAATT
CGTTTCAGTTGAACTTTGGTCTCTGCGTATTTCTTTCTTATCTAGTTTCCATGGCTACGTAGATAAGTAG
CATGGCGGGTTAATCATTAACTACAGCCCGGGCGTTTAAACAGCGGGCGGAGGGGTGGAGTCGTGAC
GTGAATTACGTCATAGGGTTAGGGAGGTCCTGTATTAGAGGTCACGTGAGTGTTTTGCGACATTTTGCG
ACACCATGTGGTCTCGCTGGGGGGGGGGGCCCGAGTGAGCACGCAGGGTCTCCATTTTGAAGCGGGA
GGTTTGAACGAGCGCTGGCGCGCTCACTGGCCGTCGTTTTACAACGTCGTGACTGGGAAAACCCTGGC
GTTACCCAACTTAATCGCCTTGCAGCACATCCCCCTTTCGCCAGCTGGCGTAATAGCGAAGAGGCCCG
CACCGATCGCCCTTCCCATGCATCGGCCGCAAATACCTGCAGGATCCGTTTTGCGCTGCTTCGCGATGT
ACGGGCCAGATATACGCGTT**GACATTGATTATTGACTAGTTATTAATAGTAATCAATTACGGGGTC
ATTAGTTCATAGCCCATATATGGAGTTCCGCGTTACATAACTTACGGTAAATGGCCCGCCTGGCT
GACCGCCCAACGACCCCCGCCCATTGACGTCAATAATGACGTATGTTCCCATAGTAACGCCAAT
AGGGACTTTCCATTGACGTCAATGGGTGGACTATTTACGGTAAACTGCCCACTTGGCAGTACAT
CAAGTGTATCATATGCCAAGTACGCCCCCTATTGACGTCAATGACGGTAAATGGCCCGCCTGGC
ATTATGCCCAGTACATGACCTTATGGGACTTTCCTACTTGGCAGTACATCTACGTATTAGTCATC
GCTATTACCATGGTGATGCGGTTTTGGCAGTACATCAATGGGCGTGGATAGCGGTTTGACTCAC
GGGGATTTCCAAGTCTCCACCCCATTGACGTCAATGGGAGTTTGTTTTGGCACCAAAATCAACG
GGACTTTCCAAAATGTCGTAACAACTCCGCCCCATTGACGCAAATGGGCGGTAGGCGTGTACGG
TGGGAGGTCTATATAAGCAGAGCT**CTCTGGCTAACTAGAGAACCCACTGCTTACTGGCTTATCGAAA
TTAATACGACTCACTATAGGGAGACCCAAGCTGGCTAGCGCCACC**ATGGAAGAGCAATACCGCCCG
GAAGAGATAGAATCCAAAGTACAGCTTCATTGGGATGAGAAGCGCACATTTGAAGTAACCGAAG
ACGAGAGCAAAGAGAAGTATTACTGCCTGTCTATCCTCCCCTATCCTTCTGGTCGACTACACATG
GGCCACGTACGTAACTACACCATCGGTGACGTGATCGCCCGCTACCAGCGTATGCTGGGCAAAA
ACGTCCTGCAGCCGATCGGCTGGGACGCGTTTGGTCTGCCTGCGGAAGGCGCGGCGGTGAAAA
ACAACACCGCTCCGGCACCGTGGACGTACGACAACATCGCGTATATGAAAAACCAGCTCAAAAT
GCTGGGCTTTGGTTATGACTGGAGCCGCGAGCTGGCAACCTGTACGCCGGAATACTACCGTTGG
GAACAGAAATTCTTCACCGAGCTGTATAAAAAAGGCCTGGTATATAAGAAGACTTCTGCGGTCA
ACTGGTGTCCGAACGACCAGACCGTACTGGCGAACGAACAAGTTATCGACGGCTGCTGCTGGCG
CTGCGATACCAAAGTTGAACGTAAAGAGATCCCGCAGTGGTTTATCAAAATCACTGCTTACGCT**

**GACGAGCTGCTCAACGATCTGGATAAACTGGATCACTGGCCAGACACCGTTAAAACCATGCAGC**
**GTAACTGGATCGGTCGTTCCGAAGGCGTGGAGATCACCTTCAACGTTAACGACTATGACAACAC**
**GCTGACCGTTTACACTACCCGCCCGGACGCGTTTATGGGTTGTACCTACCTGGCGGTAGCTGCG**
**GGTCATCCGCTGGCGCAGAAAGCGGCGGAAAATAATCCTGAACTGGCGGCCTTTATTGACGAAT**
**GCCGTAACACCAAAGTTGCCGAAGCTGAAATGGCGACGATGGAGAAAAAAGGCGTCGATACTG**
**GCTTTAAAGCGGTTCACCCATTAACGGGCGAAGAAATTCCCGTTTGGGCAGCAAACTTCGTATT**
**GATGGAGTACGGCACGGGCGCAGTTATGGCGGTACCGGGGCACGACCAGCGCGACTACGAGTT**
**TGCCTCTAAATACGGCCTGAACATCAAACCGGTTATCCTGGCAGCTGACGGCTCTGAGCCAGAT**
**CTTTCTCAGCAAGCCCTGACTGAAAAAGGCGTGCTGTTCAACTCTGGCGAGTTCAACGGTCTTG**
**ACCATGAAGCGGCCTTCAACGCCATCGCCGATAAACTGACTGCGATGGGCGTTGGCGAGCGTAA**
**AGTGAACTACCGCCTGCGCGACTGGGGTGTTTCCCGTCAGCGTTACTGGGGCGCGCCGATTCCG**
**ATGGTGACTCTAGAAGACGGTACCGTAATGCCGACCCCGGACGACCAGCTGCCGGTGATCCTGC**
**CGGAAGATGTGGTAATGGACGGCATTACCAGCCCGATTAAAGCAGATCCGGAGTGGGCGAAAA**
**CTACCGTTAACGGTATGCCAGCACTGCGTGAAACCGACACTTTCGACACCTTTATGGAGTCCTCC**
**TGGATCTATGCGCGCTACACTTGCCCGCAGTACAAAGAAGGTATGCTGGATTCCGAAGCGGCTA**
**ACTACTGGCTGCCGGTGGATATCGCTATTGGTGGTATTGAACACGCCATTATGGGTCTGCTCTA**
**CTTCCGCTTCTTCCACAAACTGATGCGTGATGCAGGCATGGTGAACTCTGACGAACCAGCGAAA**
**CAGTTGCTGTGTCAGGGTATGGTGCTGGCAGATGCCTTCTACTATGTTGGCGAAAACGGCGAAC**
**GTAACTGGGTTTCCCCGGTTGATGCTATCGTTGAACGTGACGAGAAAGGCCGTATCGTGAAAGC**
**GAAAGATGCGGCAGGCCATGAACTGGTTTATACCGGCATGAGCAAAATGTCCAAGTCGAAGAAC**
**AACGGTATCGACCCGCAGGTGATGGTTGAACGTTACGGCGCGGACACCGTTCGTCTGTTTATGA**
**TGTTTGCTTCTCCGGCTGATATGACTCTCGAATGGCAGGAATCCGGTGTGGAAGGGGCTAACCG**
**CTTCCTGAAACGTGTCTGGAAACTGGTTTACGAGCACACAGCAAAAGGTGATGTTGCGGCACTG**
**AACGTTGATGCGCTGACTGAAAATCAGAAAGCGCTGCGTCGCGATGTGCATAAAACGATCGCTA**
**AAGTGACCGATGATATCGGCCGTCGTCAGACCTTCAACACCGCAATTGCGGCGATTATGGAGCT**
**GATGAACAAACTGGCGAAAGCACCAACCGATGGCGAGCAGGATCGCGCTCTGATGCAGGAAGC**
**ACTGCTGGCCGTTGTCCGTATGCTTAACCCGTTCACCCCGCACATCTGCTTCACGCTGTGGCAG**
**GAACTGAAAGGCGAAGGCGATATCGACAACGCGCCGTGGCCGGTTGCTGACGAAAAAGCGATG**
**GTGGAAGACTCCACGCTGGTCGTGGTGCAGGTTAACGGTAAAGTCCGTGCCAAAATCACCGTTC**
**CGGTGGACGCAACGGAAGAACAGGTTCGCGAACGTGCTGGCCAGGAACATCTGGTAGCAAAAT**
**ATCTTGATGGCGTTACTGTACGTAAAGTGATTTACGTACCAGGTAAACTCCTCAATCTGGTCGTT**
**GGCGGGCCCGTTTAAT**GAATTCAACGCGTTAAGTCGACTTTAACTCGAGTCTAGAGGGCCCGTTTAA
ACCCGCTGATCAGCCTCGACTGTGCCTTCTAGTTGCCAGCCATCTGTTGTTTGCCCCTCCCCCGTGCCTT
CCTTGACCCTGGAAGGTGCCACTCCCACTGTCCTTTCCTAATAAAATGAGGAAATTGCATCGCATTGTC
TGAGTAGGTGTCATTCTATTCTGGGGGGTGGGGTGGGGCAGGACAGCAAGGGGGAGGATTGGGAAGA
CAATAGCAGGCATGCTGGGGATGCGGTGGGCTCTATGGCTTCTGAGGCGGAAAGAACCCTAGGGGTG
CGAGCGGATCGAGCAGTGTCGATCACTACTGGACCGCGAGCTGTGCTGCGACCCGTGATCTTACGGCA
TTATACGTATGATCGGTCCACGATCAGCTAGATTATCTAGTCAGCTTGATGTCATAGCTGTTTCCTGAG
GCTCAATACTGACCATTTAAATCATACCTGACCTCCATAGCAGAAAGTCAAAAGCCTCCGACCGGAGG
CTTTTGACTTGATCGGCACGTAAGAGGTTCCAACTTTCACCATAATGAAATAAGATCACTACCGGGCG
TATTTTTTGAGTTATCGAGATTTTCAGGAGCTAAGGAAGCTAAAATGAGCCATATTCAACGGGAAACG
TCTTGCTTGAAGCCGCGATTAAATTCCAACATGGATGCTGATTTATATGGGTATAAATGGGCTCGCGAT
AATGTCGGGCAATCAGGTGCGACAATCTATCGATTGTATGGGAAGCCCGATGCGCCAGAGTTGTTTCT
GAAACATGGCAAAGGTAGCGTTGCCAATGATGTTACAGATGAGATGGTCAGGCTAAACTGGCTGACG
GAATTTATGCCTCTTCCGACCATCAAGCATTTTATCCGTACTCCTGATGATGCATGGTTACTCACCACT
GCGATCCCAGGGAAAACAGCATTCCAGGTATTAGAAGAATATCCTGATTCAGGTGAAAATATTGTTGA
TGCGCTGGCAGTGTTCCTGCGCCGGTTGCATTCGATTCCTGTTTGTAATTGTCCTTTTAACGGCGATCGC
GTATTTCGTCTCGCTCAGGCGCAATCACGAATGAATAACGGTTTGGTTGGTGCGAGTGATTTTGATGAC
GAGCGTAATGGCTGGCCTGTTGAACAAGTCTGGAAAGAAATGCATAAACTCTTGCCATTCTCACCGGA
TTCAGTCGTCACTCATGGTGATTTCTCACTTGATAACCTTATTTTTGACGAGGGGAAATTAATAGGTTG
TATTGATGTTGGACGAGTCGGAATCGCAGACCGATACCAGGATCTTGCCATCCTATGGAACTGCCTCG
GTGAGTTTTCTCCTTCATTACAGAAACGGCTTTTTCAAAAATATGGTATTGATAATCCTGATATGAATA
AATTGCAGTTTCACTTGATGCTCGATGAGTTTTTCTAATGAGGACCTAAATGTAATCACCTGGCTCACC
TTCGGGTGGGCCTTTCTGCGTTGCTGGCGTTTTTCCATAGGCTCCGCCCCCCTGACGAGCATCACAAAA
ATCGATGCTCAAGTCAGAGGTGGCGAAACCCGACAGGACTATAAAGATACCAGGCGTTTCCCCCTGGA
AGCTCCCTCGTGCGCTCTCCTGTTCCGACCCTGCCGCTTACCGGATACCTGTCCGCCTTTCTCCCTTCGG

GAAGCGTGGCGCTTTCTCATAGCTCACGCTGTAGGTATCTCAGTTCGGTGTAGGTCGTTCGCTCCAAGC
TGGGCTGTGTGCACGAACCCCCCGTTCAGCCCGACCGCTGCGCCTTATCCGGTAACTATCGTCTTGAGT
CCAACCCGGTAAGACACGACTTATCGCCACTGGCAGCAGCCACTGGTAACAGGATTAGCAGAGCGAG
GTATGTAGGCGGTGCTACAGAGTTCTTGAAGTGGTGGCCTAACTACGGCTACACTAGAAGAACAGTAT
TTGGTATCTGCGCTCTGCTGAAGCCAGTTACCTCGGAAAAAGAGTTGGTAGCTCTTGATCCGGCAAAC
AAACCACCGCTGGTAGCGGTGGTTTTTTTGTTTGCAAGCAGCAGATTACGCGCAGAAAAAAAGGATCT
CAAGAAGATCCTTTGATTTTCTACCGAAGAAAGGCCCA

*pAcBac2-CMV-PLRS1-CMV-EGFP(wt)*

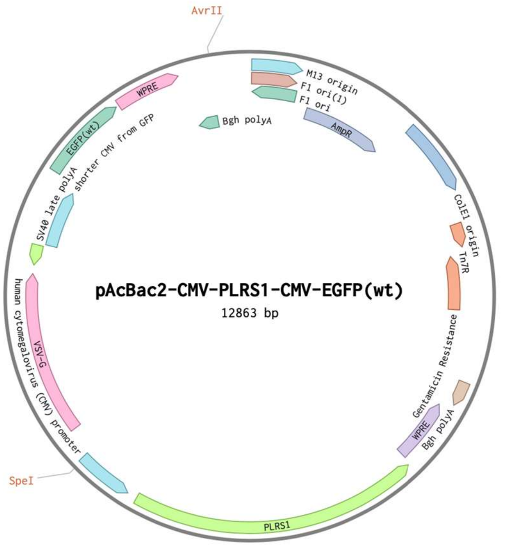


CTGAATGGCGAATGGGACGCGCCCTGTAGCGGCGCATTAAGCGCGGCGGGTGTGGTGGTTACGCGCA
GCGTGACCGCTACACTTGCCAGCGCCCTAGCGCCCGCTCCTTTCGCTTTCTTCCCTTCCTTTCTCGCCAC
GTTCGCCGGCTTTCCCCGTCAAGCTCTAAATCGGGGGCTCCCTTTAGGGTTCCGATTTAGTGCTTTACG
GCACCTCGACCCCAAAAAACTTGATTAGGGTGATGGTTCACGTAGTGGGCCATCGCCCTGATAGACGG
TTTTTCGCCCTTTGACGTTGGAGTCCACGTTCTTTAATAGTGGACTCTTGTTCCAAACTGGAACAACAC
TCAACCCTATCTCGGTCTATTCTTTTGATTTATAAGGGATTTTGCCGATTTCGGCCTATTGGTTAAAAAA
TGAGCTGATTTAACAAAAATTTAACGCGAATTTTAACAAAATATTAACGTTTACAATTTCAGGTGGCA
CTTTTCGGGGAAATGTGCGCGGAACCCCTATTTGTTTATTTTTCTAAATACATTCAAATATGTATCCGCT
CATGAGACAATAACCCTGATAAATGCTTCAATAATATTGAAAAAGGAAGAGTATGAGTATTCAACATT
TCCGTGTCGCCCTTATTCCCTTTTTTGCGGCATTTTGCCTTCCTGTTTTTGCTCACCCAGAAACGCTGGT
GAAAGTAAAAGATGCTGAAGATCAGTTGGGTGCACGAGTGGGTTACATCGAACTGGATCTCAACAGC
GGTAAGATCCTTGAGAGTTTTCGCCCCGAAGAACGTTTTCCAATGATGAGCACTTTTAAAGTTCTGCTA
TGTGGCGCGGTATTATCCCGTATTGACGCCGGGCAAGAGCAACTCGGTCGCCGCATACACTATTCTCA
GAATGACTTGGTTGAGTACTCACCAGTCACAGAAAAGCATCTTACGGATGGCATGACAGTAAGAGAAT
TATGCAGTGCTGCCATAACCATGAGTGATAACACTGCGGCCAACTTACTTCTGACAACGATCGGAGGA
CCGAAGGAGCTAACCGCTTTTTTGCACAACATGGGGGATCATGTAACTCGCCTTGATCGTTGGGAACC
GGAGCTGAATGAAGCCATACCAAACGACGAGCGTGACACCACGATGCCTGTAGCAATGGCAACAACG
TTGCGCAAACTATTAACTGGCGAACTACTTACTCTAGCTTCCCGGCAACAATTAATAGACTGGATGGA
GGCGGATAAAGTTGCAGGACCACTTCTGCGCTCGGCCCTTCCGGCTGGCTGGTTTATTGCTGATAAATC
TGGAGCCGGTGAGCGTGGGTCTCGCGGTATCATTGCAGCACTGGGGCCAGATGGTAAGCCCTCCCGTA
TCGTAGTTATCTACACGACGGGGAGTCAGGCAACTATGGATGAACGAAATAGACAGATCGCTGAGAT
AGGTGCCTCACTGATTAAGCATTGGTAACTGTCAGACCAAGTTTACTCATATATACTTTAGATTGATTT
AAAACTTCATTTTTAATTTAAAAGGATCTAGGTGAAGATCCTTTTTGATAATCTCATGACCAAAATCCC
TTAACGTGAGTTTTCGTTCCACTGAGCGTCAGACCCCGTAGAAAAGATCAAAGGATCTTCTTGAGATC
CTTTTTTTCTGCGCGTAATCTGCTGCTTGCAAACAAAAAAACCACCGCTACCAGCGGTGGTTTGTTTGC
CGGATCAAGAGCTACCAACTCTTTTTCCGAAGGTAACTGGCTTCAGCAGAGCGCAGATACCAAATACT
GTCCTTCTAGTGTAGCCGTAGTTAGGCCACCACTTCAAGAACTCTGTAGCACCGCCTACATACCTCGCT
CTGCTAATCCTGTTACCAGTGGCTGCTGCCAGTGGCGATAAGTCGTGTCTTACCGGGTTGGACTCAAGA
CGATAGTTACCGGATAAGGCGCAGCGGTCGGGCTGAACGGGGGGTTCGTGCACACAGCCCAGCTTGG
AGCGAACGACCTACACCGAACTGAGATACCTACAGCGTGAGCATTGAGAAAGCGCCACGCTTCCCGA
AGGGAGAAAGGCGGACAGGTATCCGGTAAGCGGCAGGGTCGGAACAGGAGAGCGCACGAGGGAGCT
TCCAGGGGGAAACGCCTGGTATCTTTATAGTCCTGTCGGGTTTCGCCACCTCTGACTTGAGCGTCGATT
TTTGTGATGCTCGTCAGGGGGGCGGAGCCTATGGAAAAACGCCAGCAACGCGGCCTTTTTACGGTTCC
TGGCCTTTTGCTGGCCTTTTGCTCACATGTTCTTTCCTGCGTTATCCCCTGATTCTGTGGATAACCGTAT
TACCGCCTTTGAGTGAGCTGATACCGCTCGCCGCAGCCGAACGACCGAGCGCAGCGAGTCAGTGAGCG

AGGAAGCGGAAGAGCGCCTGATGCGGTATTTTCTCCTTACGCATCTGTGCGGTATTTCACACCGCAGA
CCAGCCGCGTAACCTGGCAAAATCGGTTACGGTTGAGTAATAAATGGATGCCCTGCGTAAGCGGGTGT
GGGCGGACAATAAAGTCTTAAACTGAACAAAATAGATCTAAACTATGACAATAAAGTCTTAAACTAG
ACAGAATAGTTGTAAACTGAAATCAGTCCAGTTATGCTGTGAAAAAGCATACTGGACTTTTGTTATGG
CTAAAGCAAACTCTTCATTTTCTGAAGTGCAAATTGCCCGTCGTATTAAAGAGGGGCGTGGCCAAGGG
CATGGTAAAGACTATATTCGCGGCGTTGTGACAATTTACCGAACAACTCCGCGGCCGGGAAGCCGATC
TCGGCTTGAACGAATTGTTAGGTGGCGGTACTTGGGTCGATATCAAAGTGCATCACTTCTTCCCGTATG
CCCAACTTTGTATAGAGAGCCACTGCGGGATCGTCACCGTAATCTGCTTGCACGTAGATCACATAAGC
ACCAAGCGCGTTGGCCTCATGCTTGAGGAGATTGATGAGCGCGGTGGCAATGCCCTGCCTCCGGTGCT
CGCCGGAGACTGCGAGATCATAGATATAGATCTCACTACGCGGCTGCTCAAACCTGGGCAGAACGTAA
GCCGCGAGAGCGCCAACAACCGCTTCTTGGTCGAAGGCAGCAAGCGCGATGAATGTCTTACTACGGA
GCAAGTTCCCGAGGTAATCGGAGTCCGGCTGATGTTGGGAGTAGGTGGCTACGTCTCCGAACTCACGA
CCGAAAAGATCAAGAGCAGCCCGCATGGATTTGACTTGGTCAGGGCCGAGCCTACATGTGCGAATGAT
GCCCATACTTGAGCCACCTAACTTTGTTTTAGGGCGACTGCCCTGCTGCGTAACATCGTTGCTGCTGCG
TAACATCGTTGCTGCTCCATAACATCAAACATCGACCCACGGCGTAACGCGCTTGCTGCTTGGATGCCC
GAGGCATAGACTGTACAAAAAAACAGTCATAACAAGCCATGAAAACCGCCACTGCGCCGTTACCACC
GCTGCGTTCGGTCAAGGTTCTGGACCAGTTGCGTGAGCGCATACGCTACTTGCATTACAGTTTACGAAC
CGAACAGGCTTATGTCAACTGGGTTCGTGCCTTCATCCGTTTCCACGGTGTGCGTCACCCGGCAACCTT
GGGCAGCAGCGAAGTCGAGGCATTTCTGTCCTGGCTGGCGAACGAGCGCAAGGTTTCGGTCTCCACGC
ATCGTCAGGCATTGGCGGCCTTGCTGTTCTTCTACGGCAAGGTGCTGTGCACGGATCTGCCCTGGCTTC
AGGAGATCGGTAGACCTCGGCCGTCGCGGCGCTTGCCGGTGGTGCTGACCCCGGATGAAGTGGTTCGC
ATCCTCGGTTTTCTGGAAGGCGAGCATCGTTTGTTCGCCCAGGACTCTAGCTATAGTTCTAGTGGTTGG
CTACGTACCCGTAGTGGCTATGGCAGGGCTTGCGCTTAATGCGCCGCTACAGGGCGCGTGGGGATACC
CCCTAGAGCCCCAGCTGGTTCTTTCCGCCTCAGAAGCCATAGAGCCCACCGCATCCCCAGCATGCCTG
CTATTGTCTTCCCAATCCTCCCCCTTGCTGTCCTGCCCCACCCCACCCCCCAGAATAGAATGACACCTA
CTCAGACAATGCGATGCAATTTCCTCATTTTATTAGGAAAGGACAGTGGGAGTGGCACCTTCCAGGGT
CAAGGAAGGCACGGGGGAGGGGCAAACAACAGATGGCTGGCAACTAGAAGGCACAGTCGAGGCTGA
TCAGCGGGTTTAAACGGGCCCTCTAGACTCGAGTTAAAGTCGACGCGGGGAGGCGGCCCAAAGGGAG
ATCCGACTCGTCTGAGGGCGAAGGCGAAGACGCGGAAGAGGCCGCAGAGCCGGCAGCAGGCCGCGG
GAAGGAAGGTCCGCTGGATTGAGGGCCGAAGGGACGTAGCAGAAGGACGTCCCGCGCAGAATCCAGG
TGGCAACACAGGCGAGCAGCCAAGGAAAGGACGATGATTTCCCCGACAACACCACGGAATTGTCAGT
GCCCAACAGCCGAGCCCCTGTCCAGCAGCGGGCAAGGCAGGCGGCGATGAGTTCCGCCGTGGCAATA
GGGAGGGGGAAAGCGAAAGTCCCGGAAAGGAGCTGACAGGTGGTGGCAATGCCCCAACCAGTGGGG
GTTGCGTCAGCAAACACAGTGCACACCACGCCACGTTGCCTGACAACGGGCCACAACTCCTCATAAAG
AGACAGCAACCAGGATTTATACAAGGAGGAGAAAATGAAAGCCATACGGGAAGCAATAGCATGATAC
AAAGGCATTAAAGCAGCGTATCCACATAGCGTAAAAGGAGCAACATAGTTAAGAATACCAGTCAATC
TTTCACAAATTTTGTAATCCAGAGGTTGATTGTCGACTTAACGCGTTGAATTCA**TTAAACGGGCCCGC**
**CAACGACCAGATTGAGGAGTTTACCTGGTACGTAAATCACTTTACGTACAGTAACGCCATCAAG**
**ATATTTGCTACCAGATGTTCCTGGCCAGCACGTTCGCGAACCTGTTCTTCCGTTGCGTCCACCG**
**GAACGGTGATTTTGGCACGGACTTTACCGTTAACCTGCACCACGACCAGCGTGGAGTCTTCCAC**
**CATCGCTTTTTCGTCAGCAACCGGCCACGGCGCGTTGTCGATATCGCCTTCGCCTTTCAGTTCCT**
**GCCACAGCGTGAAGCAGATGTGCGGGGTGAACGGGTTAAGCATACGGACAACGGCCAGCAGTG**
**CTTCCTGCATCAGAGCGCGATCCTGCTCGCCATCGGTTGGTGCTTTCGCCAGTTTGTTCATCAGC**
**TCCATAATCGCCGCAATTGCGGTGTTGAAGGTCTGACGACGGCCGATATCATCGGTCACTTTAG**
**CGATCGTTTTATGCACATCGCGACGCAGCGCTTTCTGATTTTCAGTCAGCGCATCAACGTTCAGT**
**GCCGCAACATCACCTTTTGCTGTGTGCTCGTAAACCAGTTTCCAGACACGTTTCAGGAAGCGGTT**
**AGCCCCTTCCACACCGGATTCCTGCCATTCGAGAGTCATATCAGCCGGAGAAGCAAACATCATA**
**AACAGACGAACGGTGTCCGCGCCGTAACGTTCAACCATCACCTGCGGGTCGATACCGTTGTTCT**
**TCGACTTGGACATTTTGCTCATGCCGGTATAAACCAGTTCATGGCCTGCCGCATCTTTCGCTTTC**
**ACGATACGGCCTTTCTCGTCACGTTCAACGATAGCATCAACCGGGGAAACCCAGTTACGTTCGC**
**CGTTTTCGCCAACATAGTAGAAGGCATCTGCCAGCACCATACCCTGACACAGCAACTGTTTCGCT**
**GGTTCGTCAGAGTTCACCATGCCTGCATCACGCATCAGTTTGTGGAAGAAGCGGAAGTAGAGCA**
**GACCCATAATGGCGTGTTCAATACCACCAATAGCGATATCCACCGGCAGCCAGTAGTTAGCCGC**
**TTCGGAATCCAGCATACCTTCTTTGTACTGCGGGCAAGTGTAGCGCGCATAGATCCAGGAGGAC**
**TCCATAAAGGTGTCGAAAGTGTCGGTTTCACGCAGTGCTGGCATACCGTTAACGGTAGTTTTCG**
**CCCACTCCGGATCTGCTTTAATCGGGCTGGTAATGCCGTCCATTACCACATCTTCCGGCAGGATC**

**ACCGGCAGCTGGTCGTCCGGGGTCGGCATTACGGTACCGTCTTCTAGAGTCACCATCGGAATCG**
**GCGCGCCCCAGTAACGCTGACGGGAAACACCCCAGTCGCGCAGGCGGTAGTTCACTTTACGCTC**
**GCCAACGCCCATCGCAGTCAGTTTATCGGCGATGGCGTTGAAGGCCGCTTCATGGTCAAGACCG**
**TTGAACTCGCCAGAGTTGAACAGCACGCCTTTTTCAGTCAGGGCTTGCTGAGAAAGATCTGGCT**
**CAGAGCCGTCAGCTGCCAGGATAACCGGTTTGATGTTCAGGCCGTATTTAGAGGCAAACTCGTA**
**GTCGCGCTGGTCGTGCCCCGGTACCGCCATAACTGCGCCCGTGCCGTACTCCATCAATACGAAG**
**TTTGCTGCCCAAACGGGAATTTCTTCGCCCGTTAATGGGTGAACCGCTTTAAAGCCAGTATCGAC**
**GCCTTTTTTCTCCATCGTCGCCATTTCAGCTTCGGCAACTTTGGTGTTACGGCATTCGTCAATAA**
**AGGCCGCCAGTTCAGGATTATTTTCCGCCGCTTTCTGCGCCAGCGGATGACCCGCAGCTACCGC**
**CAGGTAGGTACAACCCATAAACGCGTCCGGGCGGGTAGTGTAAACGGTCAGCGTGTTGTCATAG**
**TCGTTAACGTTGAAGGTGATCTCCACGCCTTCGGAACGACCGATCCAGTTACGCTGCATGGTTTT**
**AACGGTGTCTGGCCAGTGATCCAGTTTATCCAGATCGTTGAGCAGCTCGTCAGCGTAAGCAGTG**
**ATTTTGATAAACCACTGCGGGATCTCTTTACGTTCAACTTTGGTATCGCAGCGCCAGCAGCAGCC**
**GTCGATAACTTGTTCGTTCGCCAGTACGGTCTGGTCGTTCGGACACCAGTTGACCGCAGAAGTC**
**TTCTTATATACCAGGCCTTTTTTATACAGCTCGGTGAAGAATTTCTGTTCCCAACGGTAGTATTC**
**CGGCGTACAGGTTGCCAGCTCGCGGCTCCAGTCATAACCAAAGCCCAGCATTTTGAGCTGGTTT**
**TTCATATACGCGATGTTGTCGTACGTCCACGGTGCCGGAGCGGTGTTGTTTTTCACCGCCGCGC**
**CTTCCGCAGGCAGACCAAACGCGTCCCAGCCGATCGGCTGCAGGACGTTTTTGCCCAGCATACG**
**CTGGTAGCGGGCGATCACGTCACCGATGGTGTAGTTACGTACGTGGCCCATGTGTAGTCGACCA**
**GAAGGATAGGGGAGGATAGACAGGCAGTAATACTTCTCTTTGCTCTCGTCTTCGGTTACTTCAA**
**ATGTGCGCTTCTCATCCCAATGAAGCTGTACTTTGGATTCTATCTCTTCCGGGCGGTATTGCTCT**
**TCCAT**GGTGGCGCTAGCCAGCTTGGGTCTCCCTATAGTGAGTCGTATTAATTTCGATAAGCCAGTAAG
CAGTGGGTTCTCTAGTTAGCCAGAG**AGCTCTGCTTATATAGACCTCCCACCGTACACGCCTACCGC**
**CCATTTGCGTCAATGGGGTGGAGACTTGGAAATCCCCGTGAGTCAAACCGCTATCCACGCCCAT**
**TGATGTACTGCCAAAACCGCATCACCATGGTAATAGCGATGACTAATACGTAGATGTACTGCCA**
**AGTAGGAAAGTCCCATAAGGTCATGTACTGGGCATAATGCCAGGCGGGCCATTTACCGTCATTG**
**ACGTCAATAGGGGGCGTACTTGGCATATGATACACTTGATGTACTGCCAAGTGGGCAGTTTACC**
**GTAAATAGTCCACCCATTGACGTCAATGGAAAGTCCCTATTGGCGTTACTATGGGAACATACGTC**
**ATTATTGACGTCAATGGGCGGGGGTCGTTGGGCGGTCAGCCAGGCGGGCCATTTACCGTAAGTT**
**ATGTAACGCGGAACTCCATATATGGGCTATGAACTAATGACCCCGTAATTGATTACTATTAATAA**
**CTAGTCAATAATCAATGTC**AACGCGTATATCTGGCCCGTACATCGCGAAGCAGCGCAAAACGGATCC
TGCAGGTATTTGCGGCCGCGGTCCGTATACTCCGGAATATTAATAGATCATGGAGATAATTAAAATGA
TAACCATCTCGCAAATAAATAAGTATTTTACTGTTTTCGTAACAGTTTTGTAATAAAAAAACCTATAAA
TATTCCGGATTATTCATACCGTCCCACCATCGGGCGCGAACTCCTAAAAAACCGCCACCATGAAGTGC
CTTTTGTACTTAGCCTTTTTATTCATTGGGGTGAATTGCAAGTTCACCATAGTTTTTCCACACAACCAAA
AAGGAAACTGGAAAAATGTTCCTTCTAATTACCATTATTGCCCGTCAAGCTCAGATTTAAATTGGCATA
ATGACTTAATAGGCACAGCCTTACAAGTCAAAATGCCCAAGAGTCACAAGGCTATTCAAGCAGACGGT
TGGATGTGTCATGCTTCCAAATGGGTCACTACTTGTGATTTCCGCTGGTATGGACCGAAGTATATAACA
CATTCCATCCGATCCTTCACTCCATCTGTAGAACAATGCAAGGAAAGCATTGAACAAACGAAACAAGG
AACTTGGCTGAATCCAGGCTTCCCTCCTCAAAGTTGTGGATATGCAACTGTGACGGATGCCGAAGCAG
TGATTGTCCAGGTGACTCCTCACCATGTGCTGGTTGATGAATACACAGGAGAATGGGTTGATTCACAG
TTCATCAACGGAAAATGCAGCAATTACATATGCCCCACTGTCCATAACTCTACAACCTGGCATTCTGAC
TATAAGGTCAAAGGGCTATGTGATTCTAACCTCATTTCCATGGACATCACCTTCTTCTCAGAGGACGGA
GAGCTATCATCCCTGGGAAAGGAGGGCACAGGGTTCAGAAGTAACTACTTTGCTTATGAAACTGGAGG
CAAGGCCTGCAAAATGCAATACTGCAAGCATTGGGGAGTCAGACTCCCATCAGGTGTCTGGTTCGAGA
TGGCTGATAAGGATCTCTTTGCTGCAGCCAGATTCCCTGAATGCCCAGAAGGGTCAAGTATCTCTGCTC
CATCTCAGACCTCAGTGGATGTAAGTCTAATTCAGGACGTTGAGAGGATCTTGGATTATTCCCTCTGCC
AAGAAACCTGGAGCAAAATCAGAGCGGGTCTTCCAATCTCTCCAGTGGATCTCAGCTATCTTGCTCCT
AAAAACCCAGGAACCGGTCCTGCTTTCACCATAATCAATGGTACCCTAAAATACTTTGAGACCAGATA
CATCAGAGTCGATATTGCTGCTCCAATCCTCTCAAGAATGGTCGGAATGATCAGTGGAACTACCACAG
AAAGGGAACTGTGGGATGACTGGGCACCATATGAAGACGTGGAAATTGGACCCAATGGAGTTCTGAG
GACCAGTTCAGGATATAAGTTTCCTTTATACATGATTGGACATGGTATGTTGGACTCCGATCTTCATCT
TAGCTCAAAGGCTCAGGTGTTCGAACATCCTCACATTCAAGACGCTGCTTCGCAACTTCCTGATGATGA
GAGTTTATTTTTTGGTGATACTGGGCTATCCAAAAATCCAATCGAGCTTGTAGAAGGTTGGTTCAGTAG
TTGGAAAAGCTCTATTGCCTCTTTTTTCTTTATCATAGGGTTAATCATTGGACTATTCTTGGTTCTCCGA
GTTGGTATCCATCTTTGCATTAAATTAAAGCACACCAAGAAAAGACAGATTTATACAGACATAGAGAT

GAACCGACTTGGAAAGTGATAAGGCCAGGCCGGCCAAGCTTGTCGAGAAGTACTAGAGGATCATAAT
CAGCCATACCACATTTGTAGAGGTTTTACTTGCTTTAAAAAACCTCCCACACCTCCCCCTGAACCTGAA
ACATAAAATGAATGCAATTGTTGTTGTTAACTTGTTTATTGCAGCTTATAATGGTTACAAATAAAGCAA
TAGCATCACAAATTTCACAAATAAAGCATTTTTTTCACTGCATTCTAGTTGTGGTTTGTCCAAACTCAT
CAATGTATCTTATCATGTCTGGATCTGATCACTGCTTGAGC**CTAGTTATTAATAGTAATCAATTACGG
GGTCATTAGTTCATAGCCCATATATGGAGTTCCGCGTTACATAACTTACGGTAAATGGCCCGCCT
GGCTGACCGCCCAACGACCCCCGCCCATTGACGTCAATAATGACGTATGTTCCCATAGTAACGC
CAATAGGGACTTTCCATTGACGTCAATGGGTGGACTATTTACGGTAAACTGCCCACTTGGCAGT
ACATCAAGTGTATCATATGCCAAGTACGCCCCCTATTGACGTCAATGACGGTAAATGGCCCGCC
TGGCATTATGCCCAGTACATGACCTTATGGGACTTTCCTACTTGGCAGTACATCTACGTATTAGT
CATCGCTATTACCATGGTGATGCGGTTTTGGCAGTACATCAATGGGCGTGGATAGCGGTTTGAC
TCACGGGGATTTCCAAGTCTCCACCCCATTGACGTCAATGGGAGTTTGTTTTGGCACCAAAATCA
ACGGGACTTTCCAAAATGTCGTAACAACTCCGCCCCATTGACGCAAATGGGCGGTAGGCGTGTA
CGGTGGGAGGTCTATATAAGCAGAGCTC**TCTGGCTAACTAGAGAACCCACTGCTTACTGGCTTATCG
AAATTAATACGACTCACTATAGGGAGACCCAAGCTGGCTAGCGCCGCCACC**ATGGTGAGCAAGGGC
GAGGAGCTGTTCACCGGGGTGGTGCCCATCCTGGTCGAGCTGGACGGCGACGTAAACGGCCAC
AAGTTCAGCGTGTCCGGCGAGGGCGAGGGCGATGCCACCTACGGCAAGCTGACCCTGAAGTTC
ATCTGCACCACCGGCAAGCTGCCCGTGCCCTGGCCCACCCTCGTGACCACCCTGACCTACGGCG
TGCAGTGCTTCAGCCGCTACCCCGACCACATGAAGCAGCACGACTTCTTCAAGTCCGCCATGCC
CGAAGGCTACGTCCAGGAGCGCACCATCTTCTTCAAGGACGACGGCAACTACAAGACCCGCGCC
GAGGTGAAGTTCGAGGGCGACACCCTGGTGAACCGCATCGAGCTGAAGGGCATCGACTTCAAG
GAGGACGGCAACATCCTGGGGCACAAGCTGGAGTACAACTACAACAGCCACAACGTCTATATCA
TGGCCGACAAGCAGAAGAACGGCATCAAGGTGAACTTCAAGATCCGCCACAACATCGAGGACG
GCAGCGTGCAGCTCGCCGACCACTACCAGCAGAACACCCCCATCGGCGACGGCCCCGTGCTGCT
GCCCGACAACCACTACCTGAGCACCCAGTCCGCCCTGAGCAAAGACCCCAACGAGAAGCGCGAT
CACATGGTCCTGCTGGAGTTCGTGACCGCCGCCGGGATCACTCTCGGCATGGACGAGCTGTACA
AGGGGCCCTTCGAACAAAAACTCATCTCAGAAGAGGATCTGAATATGCATACCGGTCATCATCA
CCATCACCATTGA**TAAGAATTCAACGCGTTAAGTCGACAATCAACCTCTGGATTACAAAATTTGTGAA
AGATTGACTGGTATTCTTAACTATGTTGCTCCTTTTACGCTATGTGGATACGCTGCTTTAATGCCTTTGT
ATCATGCTATTGCTTCCCGTATGGCTTTCATTTTCTCCTCCTTGTATAAATCCTGGTTGCTGTCTCTTTAT
GAGGAGTTGTGGCCCGTTGTCAGGCAACGTGGCGTGGTGTGCACTGTGTTTGCTGACGCAACCCCCAC
TGGTTGGGGCATTGCCACCACCTGTCAGCTCCTTTCCGGGACTTTCGCTTTCCCCCTCCCTATTGCCACG
GCGGAACTCATCGCCGCCTGCCTTGCCCGCTGCTGGACAGGGGCTCGGCTGTTGGGCACTGACAATTC
CGTGGTGTTGTCGGGGAAATCATCGTCCTTTCCTTGGCTGCTCGCCTGTGTTGCCACCTGGATTCTGCG
CGGGACGTCCTTCTGCTACGTCCCTTCGGCCCTCAATCCAGCGGACCTTCCTTCCCGCGGCCTGCTGCC
GGCTCTGCGGCCTCTTCCGCGTCTTCGCCTTCGCCCTCAGACGAGTCGGATCTCCCTTTGGGCCGCCTC
CCCGCGTCGACTTTAACTCGAGTCTAGAGGGCCCGTTTAAACCCGCTGATCAGCCTCGACTGTGCCTTC
TAGTTGCCAGCCATCTGTTGTTTGCCCCTCCCCCGTGCCTTCCTTGACCCTGGAAGGTGCCACTCCCACT
GTCCTTTCCTAATAAAATGAGGAAATTGCATCGCATTGTCTGAGTAGGTGTCATTCTATTCTGGGGGGT
GGGGTGGGGCAGGACAGCAAGGGGGAGGATTGGGAAGACAATAGCAGGCATGCTGGGGATGCGGTG
GGCTCTATGGCTTCTGAGGCGGAAAGAACCCTAGGAGATCCGAACCAGATAAGTGAAATCTAGTTCCA
AACTATTTTGTCATTTTTAATTTTCGTATTAGCTTACGACGCTACACCCAGTTCCCATCTATTTTGTCAC
TCTTCCCTAAATAATCCTTAAAAACTCCATTTCCACCCCTCCCAGTTCCCAACTATTTTGTCCGCCCACA
GCGGGGCATTTTTCTTCCTGTTATGTTTTAATCAAACATCCTGCCAACTCCATGTGACAAACCGTCAT
CTTCGGCTACTTTTTCTCTGTCACAGAATGAAAATTTTTCTGTCATCTCTTCGTTATTAATGTTTGTAATT
GACTGAATATCAACGCTTATTTGCAGC

*pAcBac2-MTH-mCherry-U6-LtR-wt(TAG)*

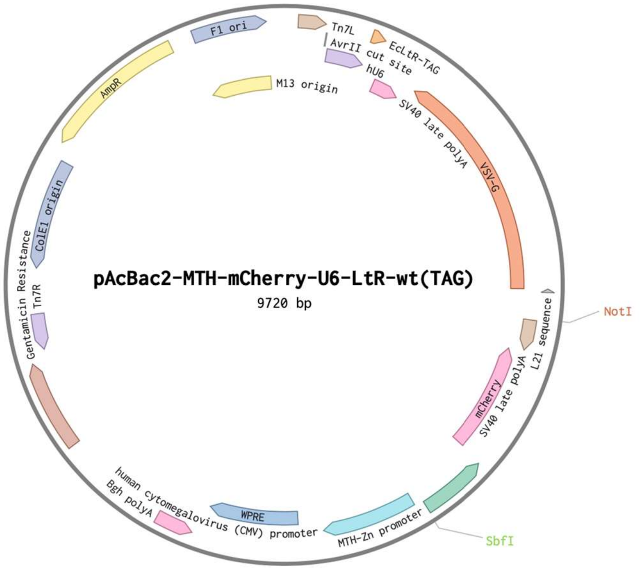


AAAGTAGCCGAAGATGACGGTTTGTCACATGGAGTTGGCAGGATGTTTGATTAAAAACATAACAGGA
AGAAAAATGCCCCGCTGTGGGCGGACAAAATAGTTGGGAACTGGGAGGGGTGGAAATGGAGTTTTTA
AGGATTATTTAGGGAAGAGTGACAAAATAGATGGGAACTGGGTGTAGCGTCGTAAGCTAATACGAAA
ATTAAAAATGACAAAATAGTTTGGAACTAGATTTCACTTATCTGGTTCGGATCTCCTAGGGGTACCTCG
GGCAGGAA**GAGGGCCTATTTCCCATGATTCCTTCATATTTGCATATACGATACAAGGCTGTTAGA
GAGATAATTAGAATTAATTTGACTGTAAACACAAAGATATTAGTACAAAATACGTGACGTAGAAA
GTAATAATTTCTTGGGTAGTTTGCAGTTTTAAAATTATGTTTTAAAATGGACTATCATATGCTTAC
CGTAACTTGAAAGTATTTCGATTTCTTGGCTTTATATATCTTGTGGAAAGGACGAAACACCGCCC
GGATGGTGGAATCGGTAGACACAAGGGATTCTAAATCCCTCGGCGTTCGCGCTGTGCGGGTTCA
AGTCCCGCTCCGGGTA**TTTTTTGCTAGGCTCAAGCAGTGATCAGATCCAGACATGATAAGATACATT
GATGAGTTTGGACAAACCACAACTAGAATGCAGTGAAAAAAATGCTTTATTTGTGAAATTTGTGATGC
TATTGCTTTATTTGTAACCATTATAAGCTGCAATAAACAAGTTAACAACAACAATTGCATTCATTTTAT
GTTTCAGGTTCAGGGGGAGGTGTGGGAGGTTTTTTAAAGCAAGTAAAACCTCTACAAATGTGGTATGG
CTGATTATGATCCTCTAGTACTTCTCGACAAGCTTGGCCGGCCTGGCCTTATCACTTTCCAAGTCGGTT
CATCTCTATGTCTGTATAAATCTGTCTTTTCTTGGTGTGCTTTAATTTAATGCAAAGATGGATACCAACT
CGGAGAACCAAGAATAGTCCAATGATTAACCCTATGATAAAGAAAAAAGAGGCAATAGAGCTTTTCC
AACTACTGAACCAACCTTCTACAAGCTCGATTGGATTTTTGGATAGCCCAGTATCACCAAAAAATAAA
CTCTCATCATCAGGAAGTTGCGAAGCAGCGTCTTGAATGTGAGGATGTTCGAACACCTGAGCCTTTGA
GCTAAGATGAAGATCGGAGTCCAACATACCATGTCCAATCATGTATAAAGGAAACTTATATCCTGAAC
TGGTCCTCAGAACTCCATTGGGTCCAATTTCCACGTCTTCATATGGTGCCCAGTCATCCCACAGTTCCC
TTTCTGTGGTAGTTCCACTGATCATTCCGACCATTCTTGAGAGGATTGGAGCAGCAATATCGACTCTGA
TGTATCTGGTCTCAAAGTATTTTAGGGTACCATTGATTATGGTGAAAGCAGGACCGGTTCCTGGGTTTT
TAGGAGCAAGATAGCTGAGATCCACTGGAGAGATTGGAAGACCCGCTCTGATTTTGCTCCAGGTTTCT
TGGCAGAGGGAATAATCCAAGATCCTCTCAACGTCCTGAATTAGACTTACATCCACTGAGGTCTGAGA
TGGAGCAGAGATACTTGACCCTTCTGGGCATTCAGGGAATCTGGCTGCAGCAAAGAGATCCTTATCAG
CCATCTCGAACCAGACACCTGATGGGAGTCTGACTCCCCAATGCTTGCAGTATTGCATTTTGCAGGCCT
TGCCTCCAGTTTCATAAGCAAAGTAGTTACTTCTGAACCCTGTGCCCTCCTTTCCCAGGGATGATAGCT
CTCCGTCCTCTGAGAAGAAGGTGATGTCCATGGAAATGAGGTTAGAATCACATAGCCCTTTGACCTTA
TAGTCAGAATGCCAGGTTGTAGAGTTATGGACAGTGGGGCATATGTAATTGCTGCATTTTCCGTTGATG
AACTGTGAATCAACCCATTCTCCTGTGTATTCATCAACCAGCACATGGTGAGGAGTCACCTGGACAAT
CACTGCTTCGGCATCCGTCACAGTTGCATATCCACAACTTTGAGGAGGGAAGCCTGGATTCAGCCAAG
TTCCTTGTTTCGTTTGTTCAATGCTTTCCTTGCATTGTTCTACAGATGGAGTGAAGGATCGGATGGAAT
GTGTTATATACTTCGGTCCATACCAGCGGAAATCACAAGTAGTGACCCATTTGGAAGCATGACACATC
CAACCGTCTGCTTGAATAGCCTTGTGACTCTTGGGCATTTTGACTTGTAAGGCTGTGCCTATTAAGTCA
TTATGCCAATTTAAATCTGAGCTTGACGGGCAATAATGGTAATTAGAAGGAACATTTTTCCAGTTTCCT
TTTTGGTTGTGTGGAAAAACTATGGTGAACTTGCAATTCACCCCAATGAATAAAAAGGCTAAGTACAA

AAGGCACTTCATGGTGGCGGTTTTTTAGGAGTTCGCGCCCGATGGTGGGACGGTATGAATAATCCGGA
ATATTTATAGGTTTTTTTATTACAAAACTGTTACGAAAACAGTAAAATACTTATTTATTTGCGAGATGG
TTATCATTTTAATTATCTCCATGATCTATTAATATTCCGGAGTATACGGACCGCGGCCGCAAATACCGC
CAGACATGATAAGATACATTGATGAGTTTGGACAAACCACAACTAGAATGCAGTGAAAAAAATGCTTT
ATTTGTGAAATTTGTGATGCTATTGCTTTATTTGTAACCATTATAAGCTGCAATAAACAAGTTAACAAC
AACAATTGCATTCATTTTATGTTTCAGGTTCAGGGGGAGGTGTGGGAGGTTTTTTAAAGCAAGTAAAA
**TTACTTGTACAGCTCGTCCATGCCGCCGGTGGAGTGGCGGCCCTCGGCGCGTTCGTACTGTTCC
ACGATGGTGTAGTCCTCGTTGTGGGAGGTGATGTCCAACTTGATGTTGACGTTGTAGGCGCCGG
GCAGCTGCACGGGCTTCTTGGCCTTGTAGGTGGTCTTGACCTCAGCGTCGTAGTGGCCGCCGTC
CTTCAGCTTCAGCCTCTGCTTGATCTCGCCCTTCAGGGCGCCGTCCTCGGGGTACATCCGCTCG
GAGGAGGCCTCCCAGCCCATGGTCTTCTTCTGCATTACGGGGCCGTCGGAGGGGAAGTTGGTGC
CGCGCAGCTTCACCTTGTAGATGAACTCGCCGTCTTGCAGGGAGGAGTCCTGGGTCACGGTCAC
CACGCCGCCGTCCTCGAAGTTCATCACGCGCTCCCACTTGAAGCCCTCGGGGAAGGACAGCTTC
AAGTAGTCGGGGATGTCGGCGGGGTGCTTCACGTAGGCCTTGGAGCCGTACATGAACTGAGGG
GACAGGATGTCCCAGGCGAAGGGCAGGGGGCCACCCTTGGTCACCTTCAGCTTGGCGGTCTGG
GTGCCCTCGTAGGGGCGGCCCTCGCCCTCGCCCTCGATCTCGAACTCGTGGCCGTTCACGGAGC
CCTCCATGTGCACCTTGAAGCGCATGAACTCCTTGATGATGGCCATGTTATCCTCCTCGCCCTTG
CTCACCAT**GGTGGCGGCCGGTA**GATCCAGACCGGCTGAGGAGCGATGAAACACAGAAACCCGAG
CGCTAGGAGAAGGCGTGATGGAGAGAAGCGTGCAGAGCGCTGCCTTTATAGAGGGAAGGATTT
GGGGCGAGCGCAAAAGGTCCGCCCAGGTTGCCCACCACCTGCATACGCCCCCGCAGACTGCAC
ACGTCCCGCAGACTGCAAGCGTCCCTCTTCCTGCACGTGGGATACCGCCTGCACACGCCTCCGC
TCTGAGTCACGCCCCTCTCCGCACGGCCCACGGACTGAGCGCAAAGGATCTGCGCTCGGACCGG
GACAGTTCCCGGAAGCTTGAGCTCGAGATCCGCGCCCGATGGTGGGACGGTATGAATAATCGG
GCCCGACGAAGACTTGATCACCCGGGATCTCGAGCCATGGT**CCTGCAGGATCCGTTTTGCGCTGCT
TCGCGATGTACGGGCCAGATATACGCGTTGACATTGATTATTGACTAGTTATTAATAGTAATCAATTAC
GGGGTCATTAGTTCATAGCCCATATATGGAGTTCCGCGTTACATAACTTACGGTAAATGGCCCGCCTG
GCTGACCGCCCAACGACCCCCGCCCATTGACGTCAATAATGACGTATGTTCCCATAGTAACGCCAATA
GGGACTTTCCATTGACGTCAATGGGTGGACTATTTACGGTAAACTGCCCACTTGGCAGTACATCAAGT
GTATCATATGCCAAGTACGCCCCCTATTGACGTCAATGACGGTAAATGGCCCGCCTGGCATTATGCCC
AGTACATGACCTTATGGGACTTTCCTACTTGGCAGTACATCTACGTATTAGTCATCGCTATTACCATGG
TGATGCGGTTTTGGCAGTACATCAATGGGCGTGGATAGCGGTTTGACTCACGGGGATTTCCAAGTCTC
CACCCCATTGACGTCAATGGGAGTTTGTTTTGGCACCAAAATCAACGGGACTTTCCAAAATGTCGTAA
CAACTCCGCCCCATTGACGCAAATGGGCGGTAGGCGTGTACGGTGGGAGGTCTATATAAGCAGAGCTC
TCTGGCTAACTAGAGAACCCACTGCTTACTGGCTTATCGAAATTAATACGACTCACTATAGGGAGACC
CAAGCTGGCTAGCGTTTAAACTTAAGCTTGGTACCGAGCTCGGATCCACTAGTCCAGTGTGGTGGAAT
TCAACGCGTTAAGTCGACAATCAACCTCTGGATTACAAAATTTGTGAAAGATTGACTGGTATTCTTAAC
TATGTTGCTCCTTTTACGCTATGTGGATACGCTGCTTTAATGCCTTTGTATCATGCTATTGCTTCCCGTA
TGGCTTTCATTTTCTCCTCCTTGTATAAATCCTGGTTGCTGTCTCTTTATGAGGAGTTGTGGCCCGTTGT
CAGGCAACGTGGCGTGGTGTGCACTGTGTTTGCTGACGCAACCCCCACTGGTTGGGGCATTGCCACCA
CCTGTCAGCTCCTTTCCGGGACTTTCGCTTTCCCCCTCCCTATTGCCACGGCGGAACTCATCGCCGCCTG
CCTTGCCCGCTGCTGGACAGGGGCTCGGCTGTTGGGCACTGACAATTCCGTGGTGTTGTCGGGGAAAT
CATCGTCCTTTCCTTGGCTGCTCGCCTGTGTTGCCACCTGGATTCTGCGCGGGACGTCCTTCTGCTACGT
CCCTTCGGCCCTCAATCCAGCGGACCTTCCTTCCCGCGGCCTGCTGCCGGCTCTGCGGCCTCTTCCGCG
TCTTCGCCTTCGCCCTCAGACGAGTCGGATCTCCCTTTGGGCCGCCTCCCCGCGTCGACTTTAACTCGA
GTCTAGAGGGCCCGTTTAAACCCGCTGATCAGCCTCGACTGTGCCTTCTAGTTGCCAGCCATCTGTTGT
TTGCCCCTCCCCCGTGCCTTCCTTGACCCTGGAAGGTGCCACTCCCACTGTCCTTTCCTAATAAAATGA
GGAAATTGCATCGCATTGTCTGAGTAGGTGTCATTCTATTCTGGGGGGTGGGGTGGGGCAGGACAGCA
AGGGGGAGGATTGGGAAGACAATAGCAGGCATGCTGGGGATGCGGTGGGCTCTATGGCTTCTGAGGC
GGAAAGAACCAGCTGGGGCTCTAGGGGGTATCCCCACGCGCCCTGTAGCGGCGCATTAAGCGCAAGC
CCTGCCATAGCCACTACGGGTACGTAGCCAACCACTAGAACTATAGCTAGAGTCCTGGGCGAACAAAC
GATGCTCGCCTTCCAGAAAACCGAGGATGCGAACCACTTCATCCGGGGTCAGCACCACCGGCAAGCGC
CGCGACGGCCGAGGTCTACCGATCTCCTGAAGCCAGGGCAGATCCGTGCACAGCACCTTGCCGTAGAA
GAACAGCAAGGCCGCCAATGCCTGACGATGCGTGGAGACCGAAACCTTGCGCTCGTTCGCCAGCCAG
GACAGAAATGCCTCGACTTCGCTGCTGCCCAAGGTTGCCGGGTGACGCACACCGTGGAAACGGATGA
AGGCACGAACCCAGTTGACATAAGCCTGTTCGGTTCGTAAACTGTAATGCAAGTAGCGTATGCGCTCA
CGCAACTGGTCCAGAACCTTGACCGAACGCAGCGGTGGTAACGGCGCAGTGGCGGTTTTCATGGCTTG

TTATGACTGTTTTTTTGTACAGTCTATGCCTCGGGCATCCAAGCAGCAAGCGCGTTACGCCGTGGGTCG
ATGTTTGATGTTATGGAGCAGCAACGATGTTACGCAGCAGCAACGATGTTACGCAGCAGGGCAGTCGC
CCTAAAACAAAGTTAGGTGGCTCAAGTATGGGCATCATTCGCACATGTAGGCTCGGCCCTGACCAAGT
CAAATCCATGCGGGCTGCTCTTGATCTTTTCGGTCGTGAGTTCGGAGACGTAGCCACCTACTCCCAACA
TCAGCCGGACTCCGATTACCTCGGGAACTTGCTCCGTAGTAAGACATTCATCGCGCTTGCTGCCTTCGA
CCAAGAAGCGGTTGTTGGCGCTCTCGCGGCTTACGTTCTGCCCAGGTTTGAGCAGCCGCGTAGTGAGA
TCTATATCTATGATCTCGCAGTCTCCGGCGAGCACCGGAGGCAGGGCATTGCCACCGCGCTCATCAAT
CTCCTCAAGCATGAGGCCAACGCGCTTGGTGCTTATGTGATCTACGTGCAAGCAGATTACGGTGACGA
TCCCGCAGTGGCTCTCTATACAAAGTTGGGCATACGGGAAGAAGTGATGCACTTTGATATCGACCCAA
GTACCGCCACCTAACAATTCGTTCAAGCCGAGATCGGCTTCCCGGCCGCGGAGTTGTTCGGTAAATTG
TCACAACGCCGCGAATATAGTCTTTACCATGCCCTTGGCCACGCCCCTCTTTAATACGACGGGCAATTT
GCACTTCAGAAAATGAAGAGTTTGCTTTAGCCATAACAAAAGTCCAGTATGCTTTTTCACAGCATAAC
TGGACTGATTTCAGTTTACAACTATTCTGTCTAGTTTAAGACTTTATTGTCATAGTTTAGATCTATTTTG
TTCAGTTTAAGACTTTATTGTCCGCCCACACCCGCTTACGCAGGGCATCCATTTATTACTCAACCGTAA
CCGATTTTGCCAGGTTACGCGGCTGGTCTGCGGTGTGAAATACCGCACAGATGCGTAAGGAGAAAATA
CCGCATCAGGCGCTCTTCCGCTTCCTCGCTCACTGACTCGCTGCGCTCGGTCGTTCGGCTGCGGCGAGC
GGTATCAGCTCACTCAAAGGCGGTAATACGGTTATCCACAGAATCAGGGGATAACGCAGGAAAGAAC
ATGTGAGCAAAAGGCCAGCAAAAGGCCAGGAACCGTAAAAAGGCCGCGTTGCTGGCGTTTTTCCATA
GGCTCCGCCCCCCTGACGAGCATCACAAAAATCGACGCTCAAGTCAGAGGTGGCGAAACCCGACAGG
ACTATAAAGATACCAGGCGTTTCCCCCTGGAAGCTCCCTCGTGCGCTCTCCTGTTCCGACCCTGCCGCT
TACCGGATACCTGTCCGCCTTTCTCCCTTCGGGAAGCGTGGCGCTTTCTCAATGCTCACGCTGTAGGTA
TCTCAGTTCGGTGTAGGTCGTTCGCTCCAAGCTGGGCTGTGTGCACGAACCCCCCGTTCAGCCCGACCG
CTGCGCCTTATCCGGTAACTATCGTCTTGAGTCCAACCCGGTAAGACACGACTTATCGCCACTGGCAGC
AGCCACTGGTAACAGGATTAGCAGAGCGAGGTATGTAGGCGGTGCTACAGAGTTCTTGAAGTGGTGG
CCTAACTACGGCTACACTAGAAGGACAGTATTTGGTATCTGCGCTCTGCTGAAGCCAGTTACCTTCGG
AAAAAGAGTTGGTAGCTCTTGATCCGGCAAACAAACCACCGCTGGTAGCGGTGGTTTTTTTGTTTGCA
AGCAGCAGATTACGCGCAGAAAAAAAGGATCTCAAGAAGATCCTTTGATCTTTTCTACGGGGTCTGAC
GCTCAGTGGAACGAAAACTCACGTTAAGGGATTTTGGTCATGAGATTATCAAAAAGGATCTTCACCTA
GATCCTTTTAAATTAAAAATGAAGTTTTAAATCAATCTAAAGTATATATGAGTAAACTTGGTCTGACAG
TTACCAATGCTTAATCAGTGAGGCACCTATCTCAGCGATCTGTCTATTTCGTTCATCCATAGTTGCCTG
ACTCCCCGTCGTGTAGATAACTACGATACGGGAGGGCTTACCATCTGGCCCCAGTGCTGCAATGATAC
CGCGAGACCCACGCTCACCGGCTCCAGATTTATCAGCAATAAACCAGCCAGCCGGAAGGGCCGAGCG
CAGAAGTGGTCCTGCAACTTTATCCGCCTCCATCCAGTCTATTAATTGTTGCCGGGAAGCTAGAGTAAG
TAGTTCGCCAGTTAATAGTTTGCGCAACGTTGTTGCCATTGCTACAGGCATCGTGGTGTCACGCTCGTC
GTTTGGTATGGCTTCATTCAGCTCCGGTTCCCAACGATCAAGGCGAGTTACATGATCCCCCATGTTGTG
CAAAAAAGCGGTTAGCTCCTTCGGTCCTCCGATCGTTGTCAGAAGTAAGTTGGCCGCAGTGTTATCACT
CATGGTTATGGCAGCACTGCATAATTCTCTTACTGTCATGCCATCCGTAAGATGCTTTTCTGTGACTGG
TGAGTACTCAACCAAGTCATTCTGAGAATAGTGTATGCGGCGACCGAGTTGCTCTTGCCCGGCGTCAA
TACGGGATAATACCGCGCCACATAGCAGAACTTTAAAAGTGCTCATCATTGGAAAACGTTCTTCGGGG
CGAAAACTCTCAAGGATCTTACCGCTGTTGAGATCCAGTTCGATGTAACCCACTCGTGCACCCAACTG
ATCTTCAGCATCTTTTACTTTCACCAGCGTTTCTGGGTGAGCAAAAACAGGAAGGCAAAATGCCGCAA
AAAAGGGAATAAGGGCGACACGGAAATGTTGAATACTCATACTCTTCCTTTTTCAATATTATTGAAGC
ATTTATCAGGGTTATTGTCTCATGAGCGGATACATATTTGAATGTATTTAGAAAAATAAACAAATAGG
GGTTCCGCGCACATTTCCCCGAAAAGTGCCACCTGAAATTGTAAACGTTAATATTTTGTTAAAATTCGC
GTTAAATTTTTGTTAAATCAGCTCATTTTTTAACCAATAGGCCGAAATCGGCAAAATCCCTTATAAATC
AAAAGAATAGACCGAGATAGGGTTGAGTGTTGTTCCAGTTTGGAACAAGAGTCCACTATTAAAGAAC
GTGGACTCCAACGTCAAAGGGCGAAAAACCGTCTATCAGGGCGATGGCCCACTACGTGAACCATCAC
CCTAATCAAGTTTTTTGGGGTCGAGGTGCCGTAAAGCACTAAATCGGAACCCTAAAGGGAGCCCCCGA
TTTAGAGCTTGACGGGGAAAGCCGGCGAACGTGGCGAGAAAGGAAGGGAAGAAAGCGAAAGGAGCG
GGCGCTAGGGCGCTGGCAAGTGTAGCGGTCACGCTGCGCGTAACCACCACACCCGCCGCGCTTAATGC
GCCGCTACAGGGCGCGTCCCATTCGCCATTCAGGCTGCAAATAAGCGTTGATATTCAGTCAATTACAA
ACATTAATAACGAAGAGATGACAGAAAAATTTTCATTCTGTGACAGAGAA

*pAcBac3-CMV-PLRS1-4xU6-LtR-wt-CAG-PAD4-R372TAG-12xHisTag*

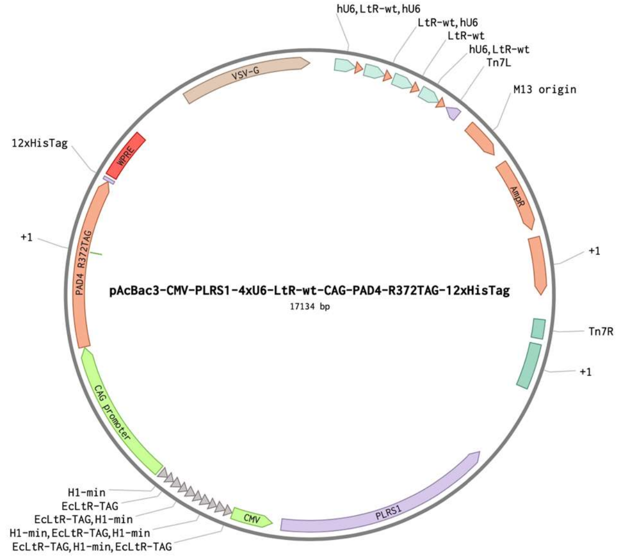


AAGCTTGTCGAGAAGTACTAGAGGATCATAATCAGCCATACCACATTTGTAGAGGTTTTACTTGCTTTA
AAAAACCTCCCACACCTCCCCCTGAACCTGAAACATAAAATGAATGCAATTGTTGTTGTTAACTTGTTT
ATTGCAGCTTATAATGGTTACAAATAAAGCAATAGCATCACAAATTTCACAAATAAAGCATTTTTTTCA
CTGCATTCTAGTTGTGGTTTGTCCAAACTCATCAATGTATCTTATCATGTCTGGATCTGATCACTGCTTG
AGCCTAGGTCGGGCAGGAA**GAGGGCCTATTTCCCATGATTCCTTCATATTTGCATATACGATACAA**
**GGCTGTTAGAGAGATAATTAGAATTAATTTGACTGTAAACACAAAGATATTAGTACAAAATACGT**
**GACGTAGAAAGTAATAATTTCTTGGGTAGTTTGCAGTTTTAAAATTATGTTTTAAAATGGACTAT**
**CATATGCTTACCGTAACTTGAAAGTATTTCGATTTCTTGGCTTTATATATCTTGTGGAAAGGACG**
**AAACACCGCCCGGATGGTGGAATCGGTAGACACAAGGGATTCTAAATCCCTCGGCGTTCGCGCT**
**GTGCGGGTTCAAGTCCCGCTCCGGGTA**TTTTTTGCTAGGTCGGGCAGGAA**GAGGGCCTATTTCCC**
**ATGATTCCTTCATATTTGCATATACGATACAAGGCTGTTAGAGAGATAATTAGAATTAATTTGAC**
**TGTAAACACAAAGATATTAGTACAAAATACGTGACGTAGAAAGTAATAATTTCTTGGGTAGTTTG**
**CAGTTTTAAAATTATGTTTTAAAATGGACTATCATATGCTTACCGTAACTTGAAAGTATTTCGATT**
**TCTTGGCTTTATATATCTTGTGGAAAGGACGAAACACCGCCCGGATGGTGGAATCGGTAGACAC**
**AAGGGATTCTAAATCCCTCGGCGTTCGCGCTGTGCGGGTTCAAGTCCCGCTCCGGGTA**TTTTTTG
CTAGGTCGGGCAGGAA**GAGGGCCTATTTCCCATGATTCCTTCATATTTGCATATACGATACAAGG**
**CTGTTAGAGAGATAATTAGAATTAATTTGACTGTAAACACAAAGATATTAGTACAAAATACGTGA**
**CGTAGAAAGTAATAATTTCTTGGGTAGTTTGCAGTTTTAAAATTATGTTTTAAAATGGACTATCA**
**TATGCTTACCGTAACTTGAAAGTATTTCGATTTCTTGGCTTTATATATCTTGTGGAAAGGACGAA**
**ACACCGCCCGGATGGTGGAATCGGTAGACACAAGGGATTCTAAATCCCTCGGCGTTCGCGCTGT**
**GCGGGTTCAAGTCCCGCTCCGGGTA**TTTTTTGCTAGGTCGGGCAGGAA**GAGGGCCTATTTCCCAT**
**GATTCCTTCATATTTGCATATACGATACAAGGCTGTTAGAGAGATAATTAGAATTAATTTGACTG**
**TAAACACAAAGATATTAGTACAAAATACGTGACGTAGAAAGTAATAATTTCTTGGGTAGTTTGCA**
**GTTTTAAAATTATGTTTTAAAATGGACTATCATATGCTTACCGTAACTTGAAAGTATTTCGATTTC**
**TTGGCTTTATATATCTTGTGGAAAGGACGAAACACCGCCCGGATGGTGGAATCGGTAGACACAA**
**GGGATTCTAAATCCCTCGGCGTTCGCGCTGTGCGGGTTCAAGTCCCGCTCCGGGTA**TTTTTTGCT
AGGAGATCCGAACCAGATAAGTGAAATCTAGTTCCAAACTATTTTGTCATTTTTAATTTTCGTATTAGC
TTACGACGCTACACCCAGTTCCCATCTATTTTGTCACTCTTCCCTAAATAATCCTTAAAAACTCCATTTC
CACCCCTCCCAGTTCCCAACTATTTTGTCCGCCCACAGCGGGGCATTTTTCTTCCTGTTATGTTTTTAAT
CAAACATCCTGCCAACTCCATGTGACAAACCGTCATCTTCGGCTACTTTTTCTCTGTCACAGAATGAAA
ATTTTTCTGTCATCTCTTCGTTATTAATGTTTGTAATTGACTGAATATCAACGCTTATTTGCAGCCTGAA
TGGCGAATGGGACGCGCCCTGTAGCGGCGCATTAAGCGCGGCGGGTGTGGTGGTTACGCGCAGCGTG
ACCGCTACACTTGCCAGCGCCCTAGCGCCCGCTCCTTTCGCTTTCTTCCCTTCCTTTCTCGCCACGTTCG
CCGGCTTTCCCCGTCAAGCTCTAAATCGGGGGCTCCCTTTAGGGTTCCGATTTAGTGCTTTACGGCACC
TCGACCCCAAAAAACTTGATTAGGGTGATGGTTCACGTAGTGGGCCATCGCCCTGATAGACGGTTTTT
CGCCCTTTGACGTTGGAGTCCACGTTCTTAATAGTGGACTCTTGTTCCAAACTGGAACAACACTCAACC

CTATCTCGGTCTATTCTTTTGATTTATAAGGGATTTTGCCGATTTCGGCCTATTGGTTAAAAAATGAGCT
GATTTAACAAAAATTTAACGCGAATTTTAACAAAATATTAACGCTTACAATTTAGGTGGCACTTTTCGG
GGAAATGTGCGCGGAACCCCTATTTGTTTATTTTTCTAAATACATTCAAATATGTATCCGCTCATGAGA
CAATAACCCTGATAAATGCTTCAATAATATTGAAAAAGGAAGAGTATGAGTATTCAACATTTCCGTGT
CGCCCTTATTCCCTTTTTTGCGGCATTTTGCCTTCCTGTTTTTGCTCACCCAGAAACGCTGGTGAAAGTA
AAAGATGCTGAAGATCAGTTGGGTGCACGAGTGGGTTACATCGAACTGGATCTCAACAGCGGTAAGA
TCCTTGAGAGTTTTCGCCCCGAAGAACGTTTTCCAATGATGAGCACTTTTAAAGTTCTGCTATGTGGCG
CGGTATTATCCCGTATTGACGCCGGGCAAGAGCAACTCGGTCGCCGCATACACTATTCTCAGAATGAC
TTGGTTGAGTACTCACCAGTCACAGAAAAGCATCTTACGGATGGCATGACAGTAAGAGAATTATGCAG
TGCTGCCATAACCATGAGTGATAACACTGCGGCCAACTTACTTCTGACAACGATCGGAGGACCGAAGG
AGCTAACCGCTTTTTTGCACAACATGGGGGATCATGTAACTCGCCTTGATCGTTGGGAACCGGAGCTG
AATGAAGCCATACCAAACGACGAGCGTGACACCACGATGCCTGTAGCAATGGCAACAACGTTGCGCA
AACTATTAACTGGCGAACTACTTACTCTAGCTTCCCGGCAACAATTAATAGACTGGATGGAGGCGGAT
AAAGTTGCAGGACCACTTCTGCGCTCGGCCCTTCCGGCTGGCTGGTTTATTGCTGATAAATCTGGAGCC
GGTGAGCGTGGGTCTCGCGGTATCATTGCAGCACTGGGGCCAGATGGTAAGCCCTCCCGTATCGTAGT
TATCTACACGACGGGGAGTCAGGCAACTATGGATGAACGAAATAGACAGATCGCTGAGATAGGTGCC
TCACTGATTAAGCATTGGTAACTGTCAGACCAAGTTTACTCATATATACTTTAGATTGATTTAAAACTT
CATTTTTAATTTAAAAGGATCTAGGTGAAGATCCTTTTTGATAATCTCATGACCAAAATCCCTTAACGT
GAGTTTTCGTTCCACTGAGCGTCAGACCCCGTAGAAAAGATCAAAGGATCTTCTTGAGATCCTTTTTTT
CTGCGCGTAATCTGCTGCTTGCAAACAAAAAAACCACCGCTACCAGCGGTGGTTTGTTTGCCGGATCA
AGAGCTACCAACTCTTTTTCCGAAGGTAACTGGCTTCAGCAGAGCGCAGATACCAAATACTGTTCTTCT
AGTGTAGCCGTAGTTAGGCCACCACTTCAAGAACTCTGTAGCACCGCCTACATACCTCGCTCTGCTAAT
CCTGTTACCAGTGGCTGCTGCCAGTGGCGATAAGTCGTGTCTTACCGGGTTGGACTCAAGACGATAGT
TACCGGATAAGGCGCAGCGGTCGGGCTGAACGGGGGGTTCGTGCACACAGCCCAGCTTGGAGCGAAC
GACCTACACCGAACTGAGATACCTACAGCGTGAGCTATGAGAAAGCGCCACGCTTCCCGAAGGGAGA
AAGGCGGACAGGTATCCGGTAAGCGGCAGGGTCGGAACAGGAGAGCGCACGAGGGAGCTTCCAGGG
GGAAACGCCTGGTATCTTTATAGTCCTGTCGGGTTTCGCCACCTCTGACTTGAGCGTCGATTTTTGTGA
TGCTCGTCAGGGGGGCGGAGCCTATGGAAAAACGCCAGCAACGCGGCCTTTTTACGGTTCCTGGCCTT
TTGCTGGCCTTTTGCTCACATGTTCTTTCCTGCGTTATCCCCTGATTCTGTGGATAACCGTATTACCGCC
TTTGAGTGAGCTGATACCGCTCGCCGCAGCCGAACGACCGAGCGCAGCGAGTCAGTGAGCGAGGAAG
CGGAAGAGCGCCTGATGCGGTATTTTCTCCTTACGCATCTGTGCGGTATTTCACACCGCATAGACCAGC
CGCGTAACCTGGCAAAATCGGTTACGGTTGAGTAATAAATGGATGCCCTGCGTAAGCGGGTGTGGGCG
GACAATAAAGTCTTAAACTGAACAAAATAGATCTAAACTATGACAATAAAGTCTTAAACTAGACAGA
ATAGTTGTAAACTGAAATCAGTCCAGTTATGCTGTGAAAAAGCATACTGGACTTTTGTTATGGCTAAA
GCAAACTCTTCATTTTCTGAAGTGCAAATTGCCCGTCGTATTAAAGAGGGGCGTGGCCAAGGGCATGG
TAAAGACTATATTCGCGGCGTTGTGACAATTTACCGAACAACTCCGCGGCCGGGAAGCCGATCTCGGC
TTGAACGAATTGTTAGGTGGCGGTACTTGGGTCGATATCAAAGTGCATCACTTCTTCCCGTATGCCCAA
CTTTGTATAGAGAGCCACTGCGGGATCGTCACCGTAATCTGCTTGCACGTAGATCACATAAGCACCAA
GCGCGTTGGCCTCATGCTTGAGGAGATTGATGAGCGCGGTGGCAATGCCCTGCCTCCGGTGCTCGCCG
GAGACTGCGAGATCATAGATATAGATCTCACTACGCGGCTGCTCAAACTTGGGCAGAACGTAAGCCGC
GAGAGCGCCAACAACCGCTTCTTGGTCGAAGGCAGCAAGCGCGATGAATGTCTTACTACGGAGCAAG
TTCCCGAGGTAATCGGAGTCCGGCTGATGTTGGGAGTAGGTGGCTACGTCTCCGAACTCACGACCGAA
AAGATCAAGAGCAGCCCGCATGGATTTGACTTGGTCAGGGCCGAGCCTACATGTGCGAATGATGCCCA
TACTTGAGCCACCTAACTTTGTTTTAGGGCGACTGCCCTGCTGCGTAACATCGTTGCTGCTGCGTAACA
TCGTTGCTGCTCCATAACATCAAACATCGACCCACGGCGTAACGCGCTTGCTGCTTGGATGCCCGAGG
CATAGACTGTACAAAAAAACAGTCATAACAAGCCATGAAAACCGCCACTGCGCCGTTACCACCGCTGC
GTTCGGTCAAGGTTCTGGACCAGTTGCGTGAGCGCATACGCTACTTGCATTACAGTTTACGAACCGAA
CAGGCTTATGTCAACTGGGTTCGTGCCTTCATCCGTTTCCACGGTGTGCGTCACCCGGCAACCTTGGGC
AGCAGCGAAGTCGAGGCATTTCTGTCCTGGCTGGCGAACGAGCGCAAGGTTTCGGTCTCCACGCATCG
TCAGGCATTGGCGGCCTTGCTGTTCTTCTACGGCAAGGTGCTGTGCACGGATCTGCCCTTGCTTCAGGA
GATCGGTAGACCTCGGCCGTCGCGGCGCTTGCCGGTGGTGCTGACCCCGGATGAAGTGGTTCGCATCC
TCGGTTTTCTGGAAGGCGAGCATCGTTTGTTCGCCCAGGACTCTAGCTATAGTTCTAGTGGTTGGCTAC
GTACCCGTAGTGGCTATGGCAGGGCTTGCGCTTAATGCGCCGCTACAGGGCGCGTGGGGATACCCCCT
AGAGCCCCAGCTGGTTCTTTCCGCCTCAGAAGCCATAGAGCCCACCGCATCCCCAGCATGCCTGCTATT
GTCTTCCCAATCCTCCCCCTTGCTGTCCTGCCCCACCCCACCCCCCAGAATAGAATGACACCTACTCAG
ACAATGCGATGCAATTTCCTCATTTTATTAGGAAAGGACAGTGGGAGTGGCACCTTCCAGGGTCAAGG

AAGGCACGGGGGAGGGGCAAACAACAGATGGCTGGCAACTAGAAGGCACAGTCGAGGCTGATCAGC
GGGTTTAAACGGGCCCTCTAGACTCGAGTTAAACGGGCCCGCCAACGACCAGATTGAGGAGTTTAC
CTGGTACGTAAATCACTTTACGTACAGTAACGCCATCAAGATATTTTGCTACCAGATGTTCCTGG
CCAGCACGTTCGCGAACCTGTTCTTCCGTTGCGTCCACCGGAACGGTGATTTTGGCACGGACTT
TACCGTTAACCTGCACCACGACCAGCGTGGAGTCTTCCACCATCGCTTTTTCGTCAGCAACCGG
CCACGGCGCGTTGTCGATATCGCCTTCGCCTTTCAGTTCCTGCCACAGCGTGAAGCAGATGTGC
GGGGTGAACGGGTTAAGCATACGGACAACGGCCAGCAGTGCTTCCTGCATCAGAGCGCGATCCT
GCTCGCCATCGGTTGGTGCTTTCGCCAGTTTGTTCATCAGCTCCATAATCGCCGCAATTGCGGTG
TTGAAGGTCTGACGACGGCCGATATCATCGGTCACTTTAGCGATCGTTTTATGCACATCGCGAC
GCAGCGCTTTCTGATTTTCAGTCAGCGCATCAACGTTCAGTGCCGCAACATCACCTTTTGCTGTG
TGCTCGTAAACCAGTTTCCAGACACGTTTCAGGAAGCGGTTAGCCCCTTCCACACCGGATTCCT
GCCATTCGAGAGTCATATCAGCCGGAGAAGCAAACATCATAAACAGACGAACGGTGTCCGCGCC
GTAACGTTCAACCATCACCTGCGGGTCGATACCGTTGTTCTTCGACTTGGACATTTTGCTCATGC
CGGTATAAACCAGTTCATGGCCTGCCGCATCTTTCGCTTTCACGATACGGCCTTTCTCGTCACGT
TCAACGATAGCATCAACCGGGGAAACCCAGTTACGTTCGCCGTTTTCGCCAACATAGTAGAAGG
CATCTGCCAGCACCATACCCTGACACAGCAACTGTTTCGCTGGTTCGTCAGAGTTCACCATGCCT
GCATCACGCATCAGTTTGTGGAAGAAGCGGAAGTAGAGCAGACCCATAATGGCGTGTTCAATAC
CACCAATAGCGATATCCACCGGCAGCCAGTAGTTAGCCGCTTCGGAATCCAGCATACCTTCTTT
GTACTGCGGGCAAGTGTAGCGCGCATAGATCCAGGAGGACTCCATAAAGGTGTCGAAAGTGTCG
GTTTCACGCAGTGCTGGCATACCGTTAACGGTAGTTTTCGCCCACTCCGGATCTGCTTTAATCGG
GCTGGTAATGCCGTCCATTACCACATCTTCCGGCAGGATCACCGGCAGCTGGTCGTCCGGGGTC
GGCATTACGGTACCGTCTTCTAGAGTCACCATCGGAATCGGCGCGCCCCAGTAACGCTGACGGG
AAACACCCCAGTCGCGCAGGCGGTAGTTCACTTTACGCTCGCCAACGCCCATCGCAGTCAGTTT
ATCGGCGATGGCGTTGAAGGCCGCTTCATGGTCAAGACCGTTGAACTCGCCAGAGTTGAACAGC
ACGCCTTTTTCAGTCAGGGCTTGCTGAGAAAGATCTGGCTCAGAGCCGTCAGCTGCCAGGATAA
CCGGTTTGATGTTCAGGCCGTATTTAGAGGCAAACTCGTAGTCGCGCTGGTCGTGCCCCGGTAC
CGCCATAACTGCGCCCGTGCCGTACTCCATCAATACGAAGTTTGCTGCCCAAACGGAATTTCTT
CGCCCGTTAATGGGTGAACCGCTTTAAAGCCAGTATCGACGCCTTTTTTCTCCATCGTCGCCATT
TCAGCTTCGGCAACTTTGGTGTTACGGCATTCGTCAATAAAGGCCGCCAGTTCAGGATTATTTTC
CGCCGCTTTCTGCGCCAGCGGATGACCCGCAGCTACCGCCAGGTAGGTACAACCCATAAACGCG
TCCGGGCGGGTAGTGTAAACGGTCAGCGTGTTGTCATAGTCGTTAACGTTGAAGGTGATCTCCA
CGCCTTCGGAACGACCGATCCAGTTACGCTGCATGGTTTTAACGGTGTCTGGCCAGTGATCCAG
TTTATCCAGATCGTTGAGCAGCTCGTCAGCGTAAGCAGTGATTTTGATAAACCACTGCGGGATCT
CTTTACGTTCAACTTTGGTATCGCAGCGCCAGCAGCAGCCGTCGATAACTTGTTCGTTCGCCAGT
ACGGTCTGGTCGTTCGGACACCAGTTGACCGCAGAAGTCTTCTTATATACCAGGCCTTTTTTATA
CAGCTCGGTGAAGAATTTCTGTTCCCAACGGTAGTATTCCGGCGTACAGGTTGCCAGCTCGCGG
CTCCAGTCATAACCAAAGCCCAGCATTTTGAGCTGGTTTTTCATATACGCGATGTTGTCGTACGT
CCACGGTGCCGGAGCGGTGTTGTTTTTCACCGCCGCGCCTTCCGCAGGCAGACCAAACGCGTCC
CAGCCGATCGGCTGCAGGACGTTTTTGCCCAGCATACGCTGGTAGCGGGCGATCACGTCACCGA
TGGTGTAGTTACGTACGTGGCCCATGTGTAGTCGACCAGAAGGATAGGGGAGGATAGACAGGC
AGTAATACTTCTCTTTGCTCTCGTCTTCGGTTACTTCAAATGTGCGCTTCTCATCCCAATGAAGC
TGTACTTTGGATTCTATCTCTTCCGGGCGGTATTGCTCTTCCATGGTGGCGGCAAGCTTAAGTTTA
AACGCTAGCCAGCTTGGGTCTCCCTATAGTGAGTCGTATTAATTTCGATAAGCCAGTAAGCAGTGGGT
TCTCTAGTTAGCCAGAGAGCTCTAGACCAAGTGACGATCACAGCGATCCACAAACAAGAACCGCG
ACCCAAATCCCGGCTGCGACGGAACTAGCTGTGCCACACCCGGCGCGTCCTTATATAATCATCG
GCGTTCACCGCCCCACGGAGATCCCTCCGCAGAATCGCCGAGAAGGGACTACTTTTCCTCGCCT
GTTCCGCTCTCTGGAAAGAAAACCAGTGCCCTAGAGTCACCCAAGTCCCGTCCTAAAATGTCCTT
CTGCTGATACTGGGGTTCTAAGGCCGAGTCTTATGAGCAGCGGGCCGCTGTCCTGAGCGTCCGG
GCGGAAGGATCAGGACGCTCGCTGCGCCCTTCGTCTGACGTGGCAGCGCTCGCCGTGAGGAGG
GGGGCGCCCGCGGGAGGCGCCAAAACCCGGCGCGGAGGCCTTCGAACGGCCACTAGTCAATAA
TCAATGTCAACGCGTATATCTGGCCCGTACATCGCGAAGCAGCGCAAAACGGATCCAAAAAATA
CCCGGAGCGGGACTTGAACCCGCACAGCGCGAACGCCGAGGGATTTAGAATCCCTTGTGTCTAC
CGATTCCACCATCCGGGCGGTGTCTCTATCACTGATAGGGAACTTATAAGTCTCTATCACTGATA
GGGATTTCACGTTTATGGTGATTTCCCAGAACACATAGCGACATGCAAATATTAGATCCAAAAAA
TACCCGGAGCGGGACTTGAACCCGCACAGCGCGAACGCCGAGGGATTTAGAATCCCTTGTGTCT
ACCGATTCCACCATCCGGGCGGTGTCTCTATCACTGATAGGGAACTTATAAGTCTCTATCACTGA

TAGGGATTTCACGTTTATGGTGATTTCCCAGAACACATAGCGACATGCAAATATTAGATCCAAAA
AATACCCGGAGCGGGACTTGAACCCGCACAGCGCGAACGCCGAGGGATTTAGAATCCCTTGTGT
CTACCGATTCCACCATCCGGGGGTGTCTCTATCACTGATAGGGAACTTATAAGTCTCTATCACTG
ATAGGGATTTCACGTTTATGGTGATTTCCCAGAACACATAGCGACATGCAAATATTAGATCCAAA
AAATACCCGGAGCGGGACTTGAACCCGCACAGCGCGAACGCCGAGGGATTTAGAATCCCTTGTG
TCTACCGATTCCACCATCCGGGCGGTGTCTCTATCACTGATAGGGAACTTATAAGTCTCTATCAC
TGATAGGGATTTCACGTTTATGGTGATTTCCCAGAACACATAGCGACATGCAAATATTAGATCCA
AAAAATACCCGGAGCGGGACTTGAACCCGCACAGCGCGAACGCCGAGGGATTTAGAATCCCTTG
TGTCTACCGATTCCACCATCCGGGCGGTGTCTCTATCACTGATAGGGAACTTATAAGTCTCTATC
ACTGATAGGGATTTCACGTTTATGGTGATTTCCCAGAACACATAGCGACATGCAAATATTAGATC
CTGCAGGCTGATCACTGCTTGAGCCTAGTTATTAATAGTAATCAATTACGGGGTCATTAGTTCATA
GCCCATATATGGAGTTCCGCGTTACATAACTTACGGTAAATGGCCCGCCTGGCTGACCGCCCAA
CGACCCCCGCCCATTGACGTCAATAATGACGTATGTTCCCATAGTAACGCCAATAGGGACTTTCC
ATTGACGTCAATGGGTGGAGTATTTACGGTAAACTGCCCACTTGGCAGTACATCAAGTGTATCAT
ATGCCAAGTACGCCCCCTATTGACGTCAATGACGGTAAATGGCCCGCCTGGCATTATGCCCAGT
ACATGACCTTATGGGACTTTCCTACTTGGCAGTACATCTACGTATTAGTCATCGCTATTACCATG
GTCGAGGTGAGCCCCACGTTCTGCTTCACTCTCCCCATCTCCCCCCCCTCCCCACCCCCAATTTT
GTATTTATTTATTTTTTAATTATTTTGTGCAGCGATGGGGGCGGGGGGGGGGGGGGCGCGCGCC
AGGCGGGGCGGGGCGGGGCGAGGGGCGGGGCGGGGCGAGGCGGAGAGGTGCGGCGGCAGCC
AATCAGAGCGGCGCGCTCCGAAAGTTTCCTTTTATGGCGAGGCGGCGGCGGCGGCGGCCCTAT
AAAAAGCGAAGCGCGCGGCGGGCGGGAGTCGCTGCGCGCTGCCTTCGCCCCGTGCCCCGCTCC
GCCGCCGCCTCGCGCCGCCCGCCCCGGCTCTGACTGACCGCGTTACTCCCACAGGTGAGCGGG
CGGGACGGCCCTTCTCCTCCGGGCTGTAATTAGCGCTTGGTTTAATGACGGCTTGTTTCTTTTCT
GTGGCTGCGTGAAAGCCTTGAGGGGCTCCGGGAGGGCCCTTTGTGCGGGGGGAGCGGCTCGGG
GGGTGCGTGCGTGTGTGTGTGCGTGGGGAGCGCCGCGTGCGGCTCCGCGCTGCCCGGCGGCTG
TGAGCGCTGCGGGCGCGGCGCGGGGCTTTGTGCGCTCCGCAGTGTGCGCGAGGGGAGCGCGGC
CGGGGGCGGTGCCCCGCGGTGCGGGGGGGGCTGCGAGGGGAACAAAGGCTGCGTGCGGGGTG
TGTGCGTGGGGGGGTGAGCAGGGGGTGTGGGCGCGTCGGTCGGGCTGCAACCCCCCCTGCACC
CCCCTCCCCGAGTTGCTGAGCACGGCCCGGCTTCGGGTGCGGGGCTCCGTACGGGGCGTGGCG
CGGGGCTCGCCGTGCCGGGCGGGGGGTGGCGGCAGGTGGGGGTGCCGGGCGGGGCGGGGCCG
CCTCGGGCCGGGGAGGGCTCGGGGGAGGGGCGCGGCGGCCCCCGGAGCGCCGGCGGCTGTCG
AGGCGCGGCGAGCCGCAGCCATTGCCTTTTATGGTAATCGTGCGAGAGGGCGCAGGGACTTCCT
TTGTCCCAAATCTGTGCGGAGCCGAAATCTGGGAGGCGCCGCCGCACCCCCTCTAGCGGGCGC
GGGGCGAAGCGGTGCGGCGCCGGCAGGAAGGAAATGGGCGGGGAGGGCCTTCGTGCGTCGCC
GCGCCGCCGTCCCCTTCTCCCTCTCCAGCCTCGGGGCTGTCCGCGGGGGGACGGCTGCCTTCGG
GGGGGACGGGGCAGGGCGGGGTTCGGCTTCTGGCGTGTGACCGGCGGCTCTAGAGCCTCTGCT
AACCATGTTCATGCCTTCTTCTTTTTCCTACAGCTCCTGGGCAACGTGCTGGTTATTGTGCTGTC
TCATCATTTTGGCAAAGAATTGGCCAAGGAGGCCACCATGGACTACAAGGACGACGACGACAAG
GCCCAGGGGACATTGATCCGTGTGACCCCAGAGCAGCCCACCCATGCCGTGTGTGTGCTGGGCA
CCTTGACTCAGCTTGACATATGCAGCTCTGCCCCTGAGGACTGCACGTCCTTCAGCATCAACGC
CTCCCCAGGGGTGGTCGTGGATATTGCCCACAGCCCTCCAGCCAAGAAGAAATCCACAGGTTCC
TCCACATGGCCCCTGGACCCTGGGGTAGAGGTGACCCTGACGATGAAAGCGGCCAGTGGTAGC
ACAGGCGACCAGAAGGTTCAGATTTCATACTACGGACCCAAGACTCCACCAGTCAAAGCTCTAC
TCTACCTCACCGCGGTGGAAATCTCCCTGTGCGCAGACATCACCCGCACCGGCAAAGTGAAGCC
AACCAGAGCTGTGAAAGATCAGAGGACCTGGACCTGGGGCCCTTGTGGACAGGGTGCCATCCT
GCTGGTGAACTGTGACAGAGACAATCTCGAATCTTCTGCCATGGACTGCGAGGATGATGAAGTG
CTTGACAGCGAAGACCTGCAGGACATGTCGCTGATGACCCTGAGCACGAAGACCCCCAAGGACT
TCTTCACAAACCATACACTGGTGCTACACGTGGCCAGGTCTGAGATGGACAAAGTGAGGGTGTT
TCAGGCCACACGGGGCAAACTGTCCTCCAAGTGCAGCGTAGTCTTGGGTCCCAAGTGGCCCTCT
CACTACCTGATGGTCCCCGGTGGAAAGCACAACATGGACTTCTACGTGGAGGCCCTCGCTTTCC
CGGACACCGACTTCCCGGGGCTCATTACCCTCACCATCTCCCTGCTGGACACGTCCAACCTGGA
GCTCCCCGAGGCTGTGGTGTTCCAAGACAGCGTGGTCTTCCGCGTGGCGCCCTGGATCATGACC
CCCAACACCCAGCCCCCGCAGGAGGTGTACGCGTGCAGTATTTTTGAAAATGAGGACTTCCTGA
AGTCAGTGACTACTCTGGCCATGAAAGCCAAGTGCAAGCTGACCATCTGCCCTGAGGAGGAGAA
CATGGATGACCAGTGGATGCAGGATGAAATGGAGATCGGCTACATCCAAGCCCCACACAAAACA
CTGCCCGTGGTCTTCGACTCTCCTTAGAACAGAGGCCTGAAGGAGTTTCCCATCAAACGAGTGA

**TGGGTCCAGATTTTGGCTATGTAACTCGAGGGCCCCAAACAGGGGGTATCAGTGGACTGGACTC
CTTTGGGAACCTGGAAGTGAGCCCCCCAGTCACAGTCAGGGGCAAGGAATACCCGCTGGGCAG
GATTCTCTTCGGGGACAGCTGTTATCCCAGCAATGACAGCCGGCAGATGCACCAGGCCCTGCAG
GACTTCCTCAGTGCCCAGCAGGTGCAGGCCCCTGTGAAGCTCTATTCTGACTGGCTGTCCGTGG
GCCACGTGGACGAGTTCCTGAGCTTTGTGCCAGCACCCGACAGGAAGGGCTTCCGGCTGCTCCT
GGCCAGCCCCAGGTCCTGCTACAAACTGTTCCAGGAGCAGCAGAATGAGGGCCACGGGGAGGC
CCTGCTGTTCGAAGGGATCAAGAAAAAAAAACAGCAGAAAATAAAGAACATTCTGTCAAACAAG
ACATTGAGAGAACATAATTCATTTGTGGAGAGATGCATCGACTGGAACCGCGAGCTGCTGAAGC
GGGAGCTGGGCCTGGCCGAGAGTGACATCATTGACATCCCGCAGCTCTTCAAGCTCAAAGAGTT
CTCTAAGGCGGAAGCTTTTTCCCCAACATGGTGAACATGCTGGTGCTAGGGAAGCACCTGGGC
ATCCCCAAGCCCTTCGGGCCAGTCATCAACGGCCGCTGCTGCCTGGAGGAGAAGGTGTGTTCCC
TGCTGGAGCCACTGGGCCTCCAGTGCACCTTCATCAACGACTTCTTCACCTACCACATCAGGCAT
GGGGAGGTGCACTGCGGCACCAACGTGCGCAGAAAGCCCTTCTCCTTCAAGTGGTGGAACATG
GTGCCCCACCACCACCACCACCACCACCACCACCACCACCACTGA**GGCCTCTAAGGCCGAATTCA
ACGCGTTAAGTCGACAATCAACCTCTGGATTACAAAATTTGTGAAAGATTGACTGGTATTCTTAACTAT
GTTGCTCCTTTTACGCTATGTGGATACGCTGCTTTAATGCCTTTGTATCATGCTATTGCTTCCCGTATGG
CTTTCATTTTCTCCTCCTTGTATAAATCCTGGTTGCTGTCTCTTTATGAGGAGTTGTGGCCCGTTGTCAG
GCAACGTGGCGTGGTGTGCACTGTGTTTGCTGACGCAACCCCCACTGGTTGGGGCATTGCCACCACCT
GTCAGCTCCTTTCCGGGACTTTCGCTTTCCCCCTCCCTATTGCCACGGCGGAACTCATCGCCGCCTGCCT
TGCCCGCTGCTGGACAGGGGCTCGGCTGTTGGGCACTGACAATTCCGTGGTGTTGTCGGGGAAATCAT
CGTCCTTTCCTTGGCTGCTCGCCTGTGTTGCCACCTGGATTCTGCGCGGGACGTCCTTCTGCTACGTCCC
TTCGGCCCTCAATCCAGCGGACCTTCCTTCCCGCGGCCTGCTGCCGGCTCTGCGGCCTCTTCCGCGTCT
TCGCCTTCGCCCTCAGACGAGTCGGATCTCCCTTTGGGCCGCCTCCCCGCGTCGACTTTAACTCGGCCA
GCACAGTGGTCGATCGACCAATGCCCTGGCTCACAAATACCACTGAGATCTTTTTCCCTCTGCCAAAA
ATTATGGGGACATCATGAAGCCCCTTGAGCATCTGACTTCTGGCTAATAAAGGAAATTTATTTTCATTG
CAATAGTGTGTTGGAATTTTTTGTGTCTCTCACTCGGAAGGACATATGGGAGGGCAAATCATTTAAAA
CATCAGAATGAGTATTTGGTTTAGAGTTTGGCAACATATGCCCATATGCTGGCTGCCATGAACAAAGG
TTGGCTATAAAGAGGTCATCAGTATATGAAACAGCCCCCTGCTGTCCATTCCTTATTCCATAGAAAAGC
CTTGACTTGAGGTTAGATTTTTTTTATATTTTGTTTTGTGTTATTTTTTTCTTTAACATCCCTAAAATTTT
CCTTACATGTTTTACTAGCCAGATTTTTCCTCCTCTCCTGACTACTCCCAGTCATAGCTGTCCCTCTTCTT
TGCGGCCGCGGTCCGTATACTCCGGAATATTAATAGATCATGGAGATAATTAAAATGATAACCATCTC
GCAAATAAATAAGTATTTTACTGTTTTCGTAACAGTTTTGTAATAAAAAAACCTATAAATATTCCGGAT
TATTCATACCGTCCCACCATCGGGCGCGAACTCCTAAAAAACCGCCACCATGAAGTGCCTTTTGTACTT
AGCCTTTTTATTCATTGGGGTGAATTGCAAGTTCACCATAGTTTTTCCACACAACCAAAAAGGAAACTG
GAAAAATGTTCCTTCTAATTACCATTATTGCCCGTCAAGCTCAGATTTAAATTGGCATAATGACTTAAT
AGGCACAGCCTTACAAGTCAAAATGCCCAAGAGTCACAAGGCTATTCAAGCAGACGGTTGGATGTGTC
ATGCTTCCAAATGGGTCACTACTTGTGATTTCCGCTGGTATGGACCGAAGTATATAACACATTCCATCC
GATCCTTCACTCCATCTGTAGAACAATGCAAGGAAAGCATTGAACAAACGAAACAAGGAACTTGGCT
GAATCCAGGCTTCCCTCCTCAAAGTTGTGGATATGCAACTGTGACGGATGCCGAAGCAGTGATTGTCC
AGGTGACTCCTCACCATGTGCTGGTTGATGAATACACAGGAGAATGGGTTGATTCACAGTTCATCAAC
GGAAAATGCAGCAATTACATATGCCCCACTGTCCATAACTCTACAACCTGGCATTCTGACTATAAGGT
CAAAGGGCTATGTGATTCTAACCTCATTTCCATGGACATCACCTTCTTCTCAGAGGACGGAGAGCTATC
ATCCCTGGGAAAGGAGGGCACAGGGTTCAGAAGTAACTACTTTGCTTATGAAACTGGAGGCAAGGCC
TGCAAAATGCAATACTGCAAGCATTGGGGAGTCAGACTCCCATCAGGTGTCTGGTTCGAGATGGCTGA
TAAGGATCTCTTTGCTGCAGCCAGATTCCCTGAATGCCCAGAAGGGTCAAGTATCTCTGCTCCATCTCA
GACCTCAGTGGATGTAAGTCTAATTCAGGACGTTGAGAGGATCTTGGATTATTCCCTCTGCCAAGAAA
CCTGGAGCAAAATCAGAGCGGGTCTTCCAATCTCTCCAGTGGATCTCAGCTATCTTGCTCCTAAAAACC
CAGGAACCGGTCCTGCTTTCACCATAATCAATGGTACCCTAAAATACTTTGAGACCAGATACATCAGA
GTCGATATTGCTGCTCCAATCCTCTCAAGAATGGTCGGAATGATCAGTGGAACTACCACAGAAAGGGA
ACTGTGGGATGACTGGGCACCATATGAAGACGTGGAAATTGGACCCAATGGAGTTCTGAGGACCAGTT
CAGGATATAAGTTTCCTTTATACATGATTGGACATGGTATGTTGGACTCCGATCTTCATCTTAGCTCAA
AGGCTCAGGTGTTCGAACATCCTCACATTCAAGACGCTGCTTCGCAACTTCCTGATGATGAGAGTTTAT
TTTTTGGTGATACTGGGCTATCCAAAAATCCAATCGAGCTTGTAGAAGGTTGGTTCAGTAGTTGGAAA
AGCTCTATTGCCTCTTTTTTCTTTATCATAGGGTTAATCATTGGACTATTCTTGGTTCTCCGAGTTGGTA
TCCATCTTTGCATTAAATTAAAGCACACCAAGAAAAGACAGATTTATACAGACATAGAGATGAACCGA
CTTGGAAAGTGATAA
